# Predictive genome-wide CRISPR-mediated gene downregulation for enhanced bioproduction

**DOI:** 10.1101/2025.04.25.650723

**Authors:** Ian S. Yunus, David N. Carruthers, Yan Chen, Jennifer W. Gin, Edward E. K. Baidoo, Christopher J. Petzold, Hector Garcia Martin, Paul D. Adams, Aindrila Mukhopadhyay, Taek Soon Lee

## Abstract

CRISPR interference (CRISPRi) has emerged as a valuable tool for redirecting metabolic flux to enhance bioproduction. However, conventional approaches for identifying target genes for CRISPRi-mediated downregulation have largely relied on heuristic methods and trial and error, which are labor-intensive and time-consuming. Additionally, the ability to achieve multigene knockdowns is limited by the constraints of molecular cloning techniques required for building multiplexed CRISPRi systems. In this study, we describe two novel methodologies FluxRETAP (Flux-REaction TArget Prioritization, a Genome-Scale Modeling Technique) and VAMMPIRE (a Versatile Assembly Method for MultiPlexing CRISPRi-mediated downREgulation). FluxRETAP accurately identified gene targets whose knockdown led to substantial increase of isoprenol titers, outperforming traditional heuristic selection. The use of VAMMPIRE enabled accurate and position-independent assembly of CRISPRi constructs containing up to five sgRNA arrays. The integration of FluxRETAP and VAMMPIRE has the potential to advance metabolic engineering by rapidly identifying CRISPRi-mediated knockdowns and knockdown combinations that enhance bioproduction titers, with potential applicability to other microbial systems.

## 1. Introduction

The production of high-value chemicals is critical to the biotechnology industry, providing essential precursors for a wide range of products, including pharmaceuticals, cosmetics, and industrial chemicals (Chen et al., 2017; d’Espaux et al., 2017; Liu et al., 2018; Luo et al., 2019; Palmer et al., 2020; Sattayawat et al., 2020; Shukal et al., 2019; Tai et al., 2015). One such high-value chemical is isoprenol, a highly versatile molecule with a wide range of applications, including as a precursor for the jet fuel 1,4-dimethylcyclooctane (DMCO) (Baral et al., 2021). However, the production of isoprenol is often limited by the efficiency of the biosynthetic pathways involved and the capacity of high titer production in a selected host organism.

CRISPR-Cas9 has emerged as a powerful tool for gene editing, offering a highly selective approach improving biochemical through targeted gene knockouts. Recently, CRISPR interference (CRISPRi) arose as a variant of the CRISPR-Cas9 system that instead utilizes a deactivated Cas9 protein. This dCas9 enzyme enables the repression of gene expression without permanent modification of the genome (Zhao et al., 2021). In the context of isoprenol biosynthesis, where current titers remain too low for commercial viability, CRISPRi offers a promising strategy to enhance yields by targeting host genes that negatively affect production.

Target gene selection for CRISPRi downregulation has, historically, followed one of the three possible approaches: heuristic, empirical, and computational. The heuristic approach involves selecting genes hypothesized to play crucial roles in increasing metabolite titers when complete information is unavailable. Early studies often rely on heuristic methods. For example, Fang and colleagues identified 108 genes related to free fatty acid (FFA) metabolism in *E. coli* for downregulation, leading to high-titer FFA production (Fang et al., 2021). Similarly, Tian and colleagues targeted 15 genes involved in lycopene biosynthesis – sharing the same precursor, IPP, with isoprenol biosynthesis – hypothesizing that their downregulation would enhance IPP and DMAPP accumulation, ultimately improving isoprenol production (Tian et al., 2019). Alternatively, empirical approaches identify genes based on experimental data. For example, Yao and colleagues downregulated multiple genes encoding aldehyde reductases to increase the aldehyde pool in *Synechocystis* sp. PCC 6803 for decarbonylation to alkanes (Yao et al., 2016). The same downregulation mechanisms were used to increase the butyraldehyde pool in *Synechocystis* for production of propane (Yunus et al., 2022). Lastly, Banerjee et al. leveraged computational tools using a constrained minimal cut set (cMCS) approach to predict 14 genes for knockdown and successfully improved indigoidine production in *P. putida* KT2440 (Banerjee et al., 2020).

The two principal challenges that hinder a wider application of CRISPRi to improve bioproduction are the identification of target genes and the construction of multiplexed CRISPRi arrays. Methods for identifying target genes for CRISPRi-mediated downregulation, for example, have primarily relied on heuristic reasoning and trial and error approaches which can be laborious and time-consuming. Additionally, computational approaches such as cMCS often assume a binary on-off logic for gene interactions, which might oversimplify the complex nature of biological networks (Banerjee et al., 2025; Klamt and Gilles, 2004; Schneider et al., 2020). In reality, gene interactions are often more nuanced, involving gradients of expression, feedback loops, and non-linear interactions (Mousavi and Lobo, 2024; Nordick and Hong, 2021). Secondly, methods to achieve multigene knockdowns have been severely limited by the existing molecular cloning methods to build multiplex CRISPRi. For example, CRISPRi arrays are prone to homologous recombination events during cloning due to the presence of repeated DNA sequences (Jiang et al., 2013; Su et al., 2016). This challenge can be minimized by creating multiple sgRNAs with distinct promoters, terminators, and sgRNA handles (Reis et al., 2019). However, the design of multiple sgRNAs can be expensive to synthesize and not universally implementable in different organisms.

To address these challenges, we leveraged FluxRETAP (Flux-REaction TArget Prioritization), a technique based on genome-scale models and introduced VAMMPIRE (Versatile Assembly Method for MultiPlexing CRISPRi-mediated downREgulation), a systematic gene assembly method. FluxRETAP is a simple and computationally inexpensive method to prioritize genes for up or downregulation when optimizing production. This computational method leverages the COBRA python (COBRApy) library (Heirendt et al., 2019) to identify target genes likely to increase production of a desired metabolite. FluxRETAP has been tested with data from the literature and shown to recover several targets experimentally tested to improve production in *E. coli* and *P. putida* (Czajka et al., 2025), showcasing its effectiveness in guiding metabolic engineering efforts *in silico*. However, its practical implementation in actual metabolic engineering studies has yet to be fully realized. To this end, VAMMPIRE leverages a linker-based modular cloning assembly method (Storch et al., 2015) to streamline the assembly of sgRNAs arrays. This facilitates the design and construction of CRISPRi gene knockdown strain with up to five sgRNAs in a single-pot assembly. To demonstrate the effectiveness of both tools, we showcase how these two methods improve the production of isoprenol in *P. putida* KT2440.

To achieve this goal, we first applied FluxRETAP to predict gene targets whose downregulation would enhance isoprenol biosynthesis, identifying 57 candidates across diverse metabolic pathways. In parallel, we used VAMMPIRE to rapidly and accurately construct CRISPRi plasmids targeting up to five genes. Through systematic screening, we demonstrated that nearly half of the FluxRETAP-predicted targets led to significantly increased isoprenol titers, outperforming genes selected by heuristic reasoning. These results validate the utility of combining FluxRETAP’s predictive power with VAMMPIRE’s streamlined assembly method to accelerate strain optimization processes, laying the groundwork for future efforts aimed at optimizing other bioproducts in industrial microorganisms.

## 2. Results

### 2.1 Identification of target genes for CRISPRi downregulation to improve isoprenol production

The highest isoprenol titer to date in *Pseudomonas* was achieved by heterologous expression of the IPP-bypass pathway in a highly engineered *Pseudomonas putida* KT2440 strain developed by our group (Banerjee et al., 2024). Given its high baseline titer, this strain was selected for our genome-wide screening efforts. The isoprenol biosynthetic pathway commences with three acetyl-CoA molecules which are condensed into 3-hydroxy-3-methylglutaryl coenzyme A (HMG-CoA), followed by four consecutive enzymatic reactions to convert HMG-CoA to isoprenol (Fig. 1A). Given the pivotal role of acetyl-CoA, NADPH, and ATP in this pathway, we heuristically targeted 51 genes involved in acetyl-CoA, cofactors, and energy metabolism in *P. putida* KT2440 for downregulation. These heuristic target genes are distributed across various metabolic pathways and are described in Supplementary Table S1 and Supplementary Fig. S1.

**Figure 1.**
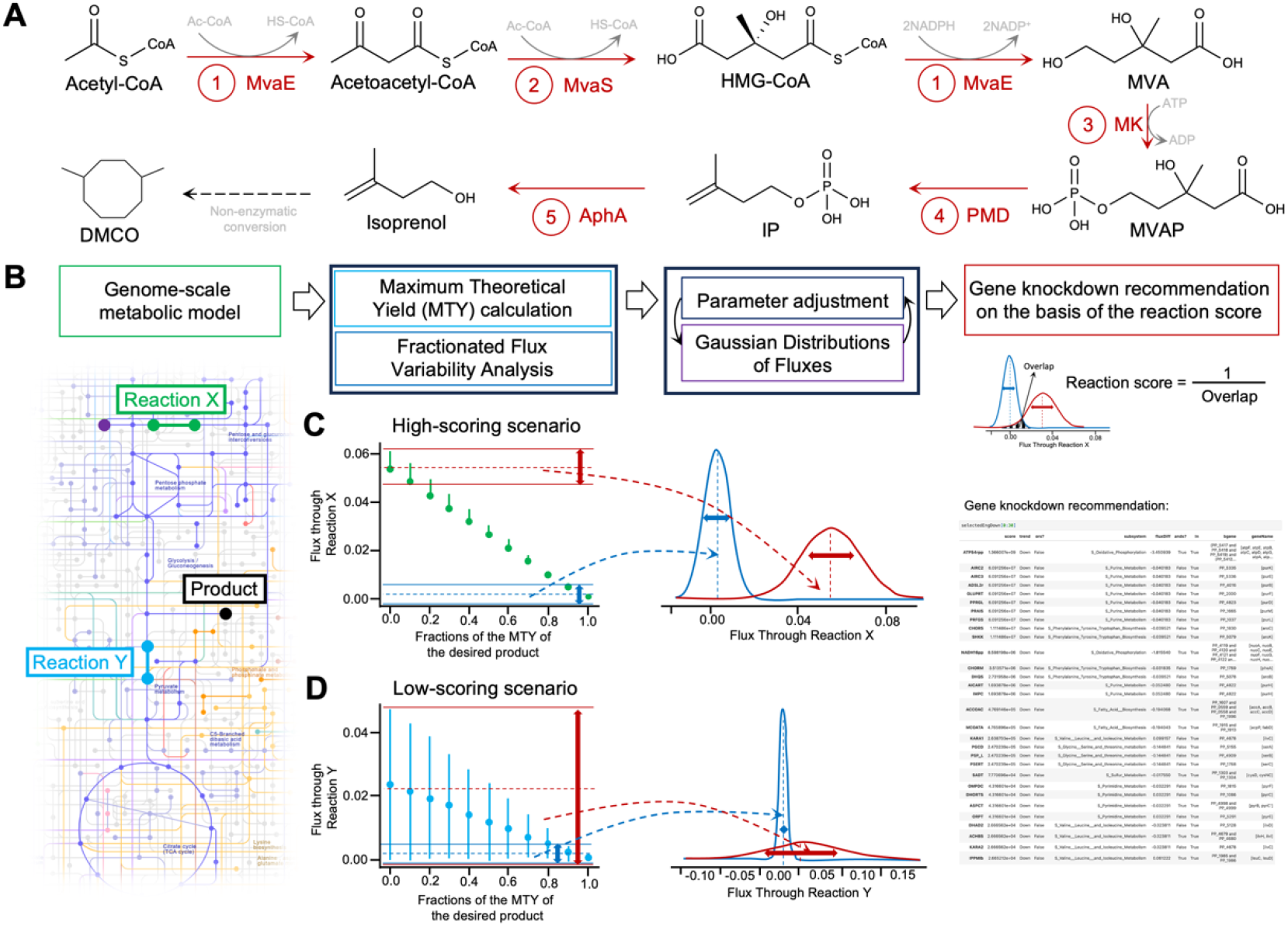
Schematic diagram of gene knockdown identification for improved production of isoprenol in *Pseudomonas putida* KT2440. (A) Metabolic pathway for production of isoprenol. (B) Workflow detailing FluxRETAP. The FluxRETAP algorithm identifies key reactions for potential genetic downregulation using Flux Variability Analysis (FVA) and Gaussian distributions of reaction fluxes. Initially, the Maximum Theoretical Yield (MTY) of the reaction corresponding to the final product in a genome-scale metabolic model is determined, then different levels of production (e.g., 10%, 20%, up to 100% of the MTY) are simulated to find the flux values compatible with each level through FVA. The flux values are represented via Gaussian distributions. Separate distributions are calculated for low production levels (e.g., 10%, 20%) and high production levels (e.g., 90%, 100%). The algorithm then calculates the overlap between these distributions. (C) Reactions with smaller overlaps (indicating large differences in flux between low and high production levels) receive higher scores, making them prime candidates for genetic downregulation. (D) Conversely, a large overlap results in a low score and indicates that the reaction’s flux doesn’t change significantly between low and high production yield. Gene knockdown recommendations are ranked based on the reaction score. A high reaction score indicates a high priority for a gene knockdown.

Next, we leveraged FluxRETAP to identify genes for knockdown that increase isoprenol production in *P. putida* KT2440 as well as to evaluate its effectiveness compared to heuristic target genes. FluxRETAP was designed to streamline the identification of knockdown targets genes in various microbes (Fig. 1B-D). By using FluxRETAP, we identified 57 genes (Fig. 2, Supplementary Table S2) and ranked them based on the inverse value of the overlap area. These genes primarily play roles in amino acid metabolism, metabolism of cofactors and vitamins, carbohydrate metabolism, transport ABC metabolism, energy metabolism, nucleotide metabolism, and lipid metabolism.

**Figure 2.**
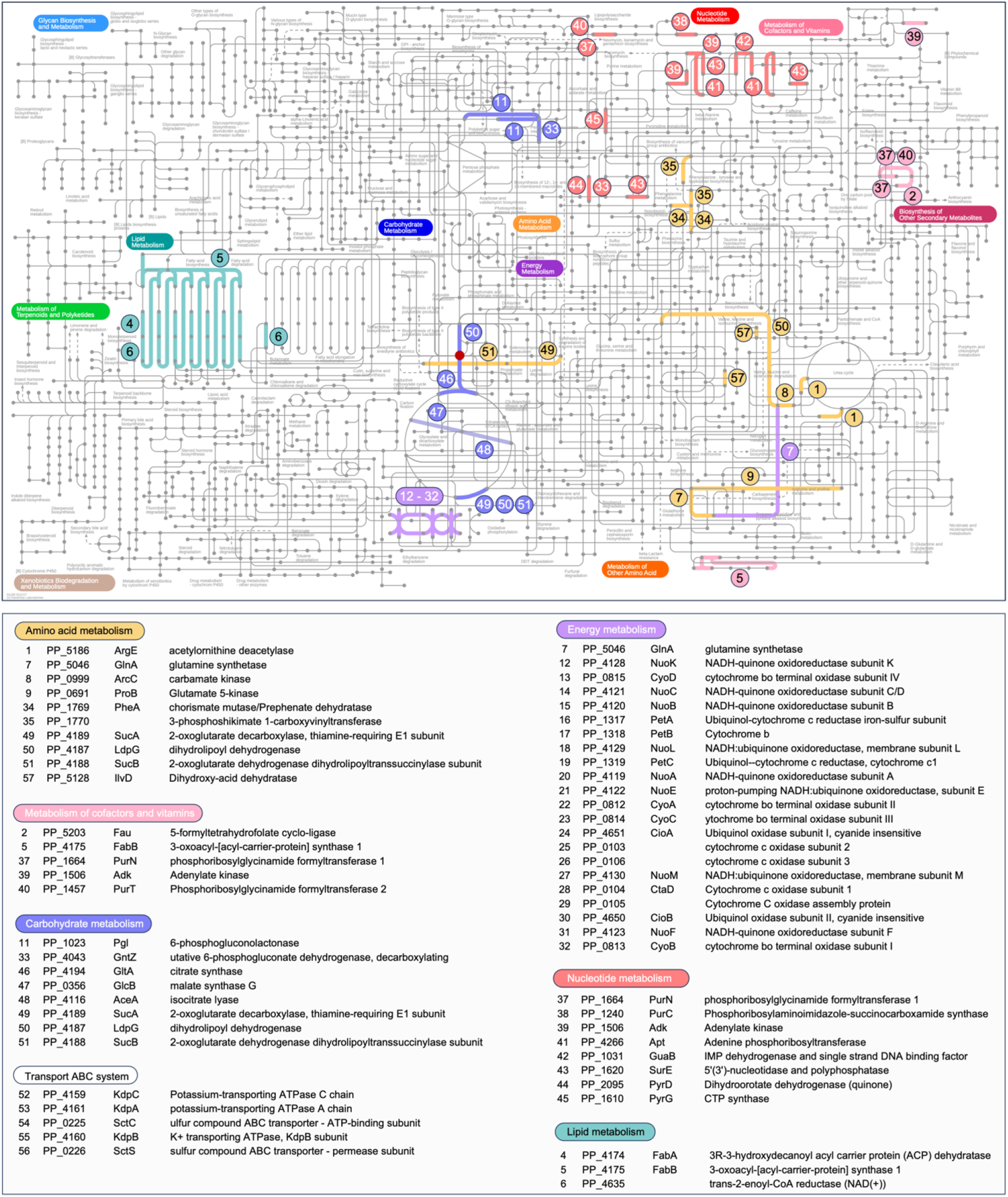
Identification of FluxRETAP target genes for CRISPRi downregulation. Figure generated using iPath3.0 (https://pathways.embl.de/) (Darzi et al., 2018). KEGG Orthology alphanumerical code for each gene was used as an input in element selection. Complete details of FluxRETAP target genes can be found in the Supplementary Table S2. Red dot represents acetyl-CoA.

### 2.2 VAMMPIRE: A Versatile Assembly Method for MultiPlexing CRISPRi-mediated downREgulation

Once we identified the target genes for downregulation, we continued to develop a genetic assembly method to rapidly and accurately create sgRNA constructs: VAMMPIRE, a Versatile Assembly Method for MultiPlexing CRISPRi-mediated downREgulation. VAMMPIRE streamlines the construction of sgRNA harboring plasmids for multi-gene downregulation through two simplified steps (Fig. 3A). First, an sgRNA oligonucleotide was digested and ligated into an sgRNA storage plasmid using a Type IIS restriction enzyme (*Bsa*I). Second, the sgRNA(s) were digested and ligated into a CRISPRi plasmid by employing a linker-based modular cloning method (Storch et al., 2015). Here, we used a constitutive J23119 promoter to control the transcription of each sgRNA as it has been shown to function in a wide range of prokaryotes (Hawkins et al., 2015; Li et al., 2023). A P_nagAa_ promoter (Banerjee et al., 2024) was used to drive dCas9 expression as it is functional in *P. putida* KT2440 but inactive in *E. coli*, thereby minimizing the expression burden of dCas9 during cloning. Using this method, we constructed 197 CRISPRi plasmids harboring a sgRNA, 41 plasmids with two sgRNAs, 24 plasmids with three sgRNAs, 11 plasmids with four sgRNAs, and two plasmids with five sgRNAs. All plasmids were constructed in a one-pot assembly and validated by PCR (Supplementary Fig. S2) and Sanger sequencing indicated 100% assembly accuracy, showcasing the versatility of this genetic assembly method compared to others (Supplementary Fig. S3, S4).

**Figure 3.**
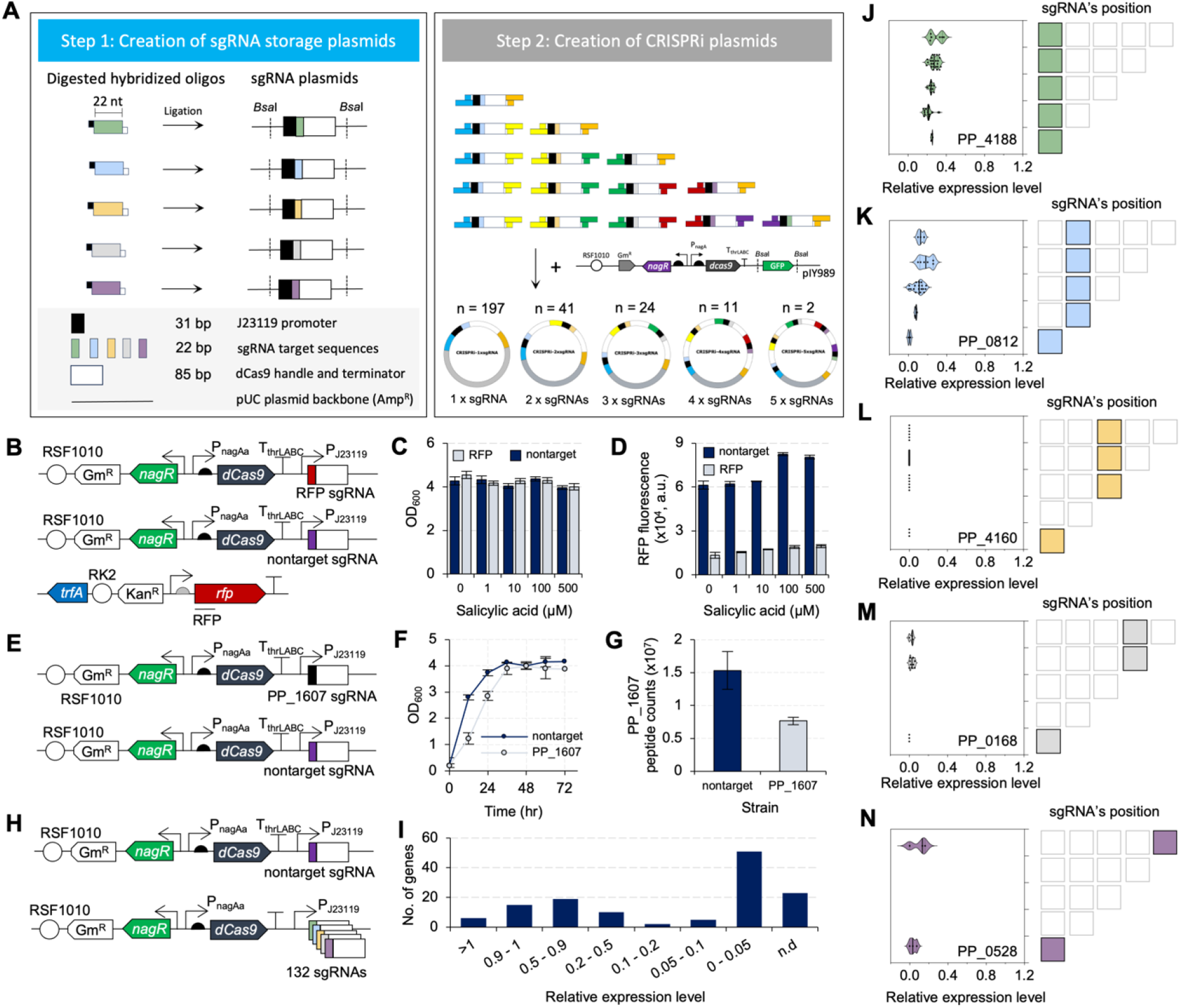
Development of VAMMPIRE, a Versatile Assembly Method for MultiPlexing CRISPRi-mediated downREgulation. (A) A two-step assembly method for multiplexing CRISPRi (detailed methods are provided in the Supplementary Note 2). (B) Plasmid diagrams used for downregulation of RFP. (C) Terminal OD_600_ and (D) RFP fluorescence measured at 48 hr from samples expressing RFP. (E) Plasmid maps for CRISPRi downregulation of *accA* (PP_1607). (F) Growth curve of nontarget and PP_1607 strains. (G) Protein counts of PP_1607 in both nontarget (control) and PP_1607 strains. (H) Schematics of plasmids used for gene downregulation in *P. putida* KT2440. (I) Summary of relative expression levels of target genes in comparison to the control strains. (J-N) Relative expression levels of target genes when used in sgRNA arrays that harbor different number of sgRNAs. Error bars represent standard deviation from three biological replicates.

To test the functionality of CRISPRi in *P. putida* KT2440, we first used the CRISPRi system to downregulate the expression of a red fluorescent protein (Fig. 3B-D) and an essential gene *accA* (encoding acetyl-CoA carboxylase carboxyl transferase subunit alpha) (Fig. 3E-G). The results indicated that the expression of RFP was not only effectively repressed, but the repression was stably maintained for at least 72 hr (Fig. 3D). Conversely, strains with CRISPRi-mediated repression of the essential gene *accA* (PP_1607) exhibited a slower growth phenotype (Fig. 3F). Proteomics analysis revealed 50% downregulation of PP_1607 by the CRISPRi (Fig. 3G). To show the effectiveness of downregulation of different genes, we performed shotgun proteomics on 132 samples carrying different sgRNAs. Proteomics analysis revealed that 68 genes were downregulated by at least 50%, 51 of which were downregulated by more than 95% (Fig. 3). Nineteen genes were downregulated by 10-50%. Fifteen genes were downregulated only by 10%. Six genes were upregulated. The expression levels of twenty-three genes could not be determined as their gene expression was not detected in the control strain. Overall, the results indicate the reliability of CRISPRi in downregulating single genes in *P. putida* KT2440.

The gene downregulation efficiency of CRISPRi arrays has been linked to the position of the target sequences in the CRISPR array (Liao et al., 2019). We hypothesized that since our sgRNA arrays are composed of an array of single transcribed sgRNAs with identical regulatory elements, each guide RNA will generate relatively constant downregulation strength regardless of its position in the array. To investigate the position dependency of our CRISPRi within the sgRNA arrays, we fixed the position of one sgRNA while varying the position of other sgRNAs in different arrays (Fig. 3J-N). We also varied the position of one sgRNA in different arrays while keeping the position of other sgRNAs unmodified (Supplementary Fig. S5). The results from both experiments indicate that the variability of gene downregulation among samples was narrow and suggest that our sgRNA arrays were less prone to context dependency for gene downregulation. Our findings are of importance for metabolic engineering studies where the Design-Build-Test-Learn (DBTL) cycle is commonly used as an iterative process to improve product titers.

### 2.3 Genome-wide CRISPRi downregulation for improved isoprenol production

To compare the efficacy of FluxRETAP and heuristic-based gene selection upon isoprenol production, we transformed the CRISPRi plasmid into a highly genetically engineered isoprenol-producing strain (IY1452) (Banerjee et al., 2024) (Fig. 4A). We then compared the isoprenol titers from all strains based on how the knockdown target genes were selected (i.e., FluxRETAP vs. heuristic). We found that the average isoprenol titers from FluxRETAP recommendations were significantly improved compared to that of the control (Fig. 4B). Moreover, these isoprenol titers were significantly higher than the titers obtained through the heuristic approach. In contrast, isoprenol titers obtained from the heuristic group were not statistically different from the control strain. Overall, among the 56 genes selected by FluxRETAP, 46.4% yielded significantly higher isoprenol titers compared to the control strain (Fig. 4C-J). Conversely, heuristic reasoning-based target gene selection resulted in only 15.7% of genes exhibiting higher isoprenol titers than the control (Supplementary Fig. S6). The highest recorded isoprenol titer of 1469 mg/L (Fig. 4E) was achieved by downregulating PP_4188, a gene identified by FluxRETAP, while in the heuristic group, the highest titer was 958 mg/L obtained from the PP_0168 downregulated strain (Supplementary Fig. S6). These results underscore the advantage of using FluxRETAP over heuristic reasoning approaches to identify genes for CRISPRi-mediated downregulation.

**Figure 4.**
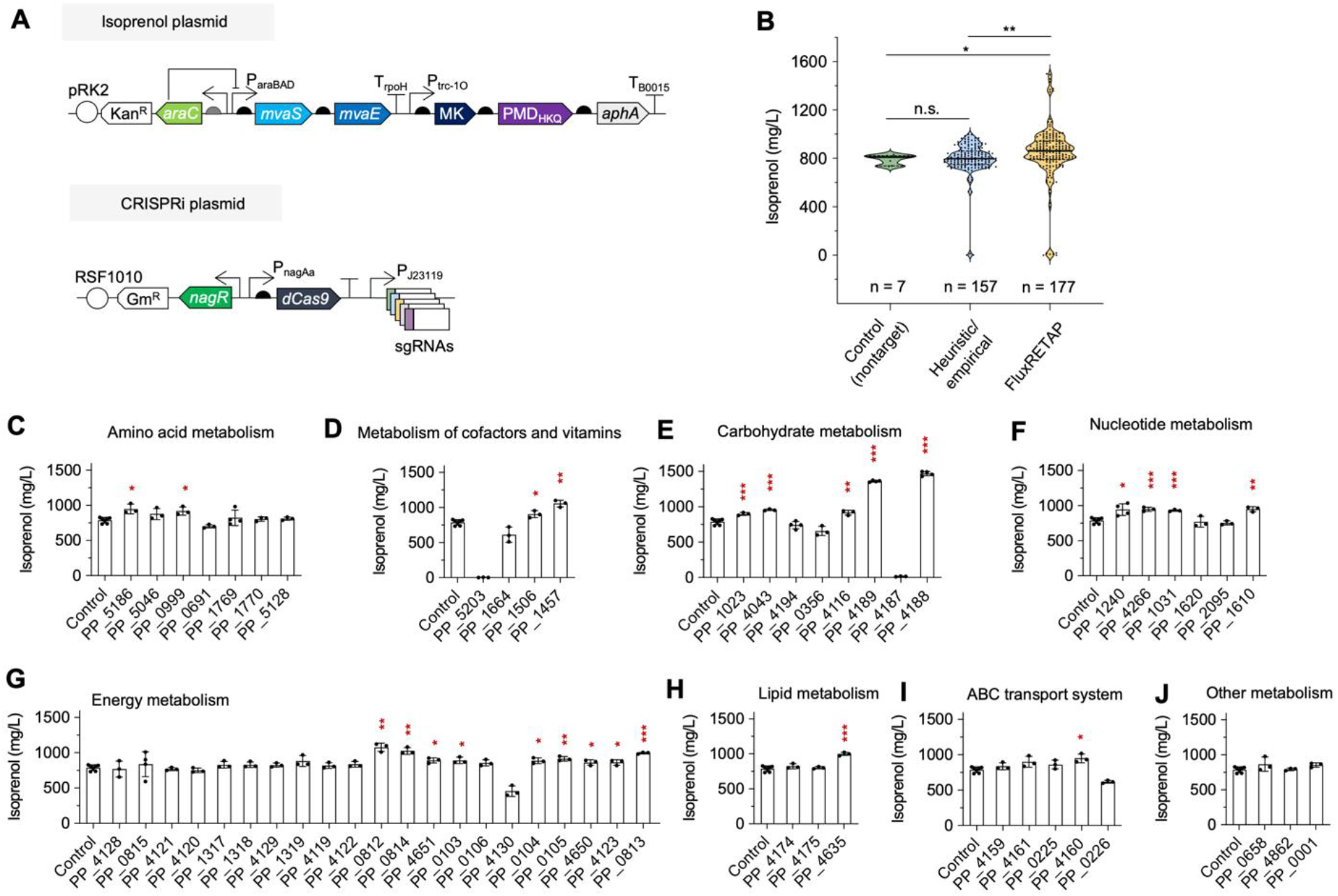
Downregulatiion of specific genes significantly improved isoprenol production. (A) Schematic diagram of plasmid used for isoprenol production and CRISPRi downregulation. (B) Comparison of isoprenol titers from heuristic group and FluxRETAP. Isoprenol titers obtained by downregulation of genes involved in various metabolisms: amino acid (C), cofactors and vitamins (D), carbohydrate (E), nucleotide (F), energy (G), lipid (H), transport (I), other (J). All samples were extracted at 48 hr. OD600 at 48 hr is shown in Supplementary Fig. S7. Error bars represent standard deviation from 3 to 6 biological replicates. Asterisks represent student’s t-test (n.s., P > 0.05; *, P ≤ 0.05; **, P ≤ 0.01; ***, P ≤ 0.001).

After analyzing isoprenol, we conducted proteomics and metabolomics studies on the PP_4188 strain to gain biological insights into its high titer. PP_4188 encodes α-ketoglutarate dehydrogenase, a key enzyme in the TCA cycle. PP_4188 appeared to be downregulated by 75% (Supplementary Fig. S8), leading to a higher accumulation of α-ketoglutarate within the cells (Fig. 5A). Proteomics analysis revealed significant changes in the proteome (Fig. 5B). Approximately, 145 genes were downregulated (Supplementary Table S3), and 193 genes were upregulated (Supplementary Table S4). Among the upregulated genes is the transglutaminase-like superfamily domain protein, encoded by PP_2686 (UniprotKB ID Q88JG3), which exhibited over 450-fold increase in expression. Transglutaminase acts on glutamine, forming γ-glutamyl thioester and releasing ammonia. In the absence of a peptide-bound lysine ε-amino group or a primary amine, this acyl-enzyme intermediate may react with water to form glutamic acid. We hypothesize that glutamic acid plays an important role in isoprenol metabolism, consistent with previous observations where glutamic acid supplementation in *Pseudomonas* growth medium reduced isoprenol catabolism (Wang et al., 2022). Pearson correlation analysis of all samples also revealed a positive correlation between the upregulation of PP_2686 and high isoprenol titer (Pearson correlation value = 0.4593, p-value = 3*10^−8^) (Supplementary Note 1), suggesting that the upregulation of PP_2686 contributing to higher isoprenol titer in the PP_4188 downregulated strain was not an isolated phenomenon.

**Figure 5.**
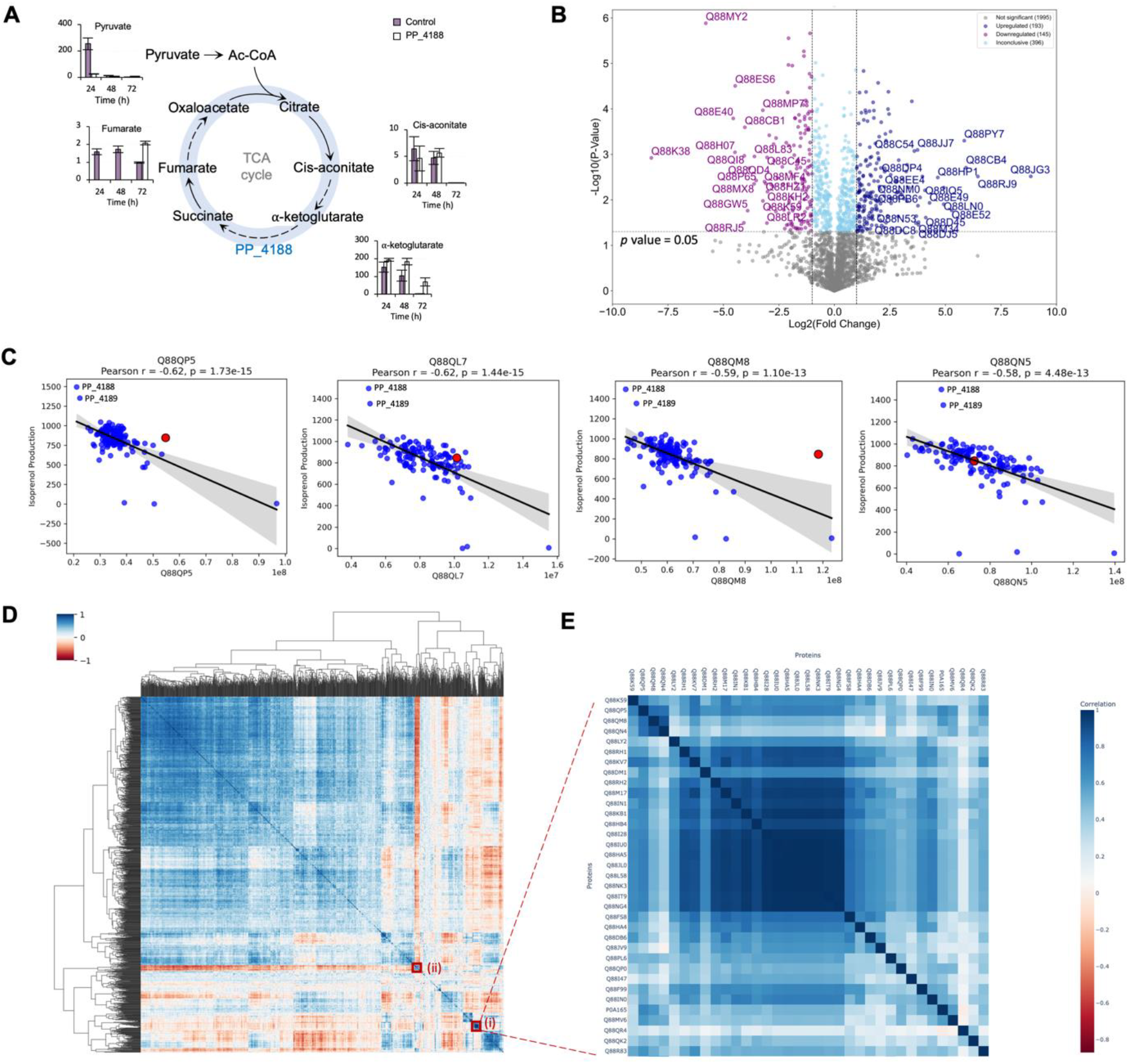
Shotgun proteomics and assessment of key metabolites in the TCA cycle. (A) Concentrations of pyruvate, fumarate, cis-aconitate, and α-ketoglutarate in μM from PP_4188 strain extracted at 24 h, 48 h, and 72 h. (B) Volcano plot representing the results of shotgun proteomic analysis from strain PP_4188 strain in comparison to the control strain. Horizontal dashed line represents the applied significance threshold of a student’s t-test *p*-value = 0.05. Vertical dashed lines represent the applied thresholds of an absolute fold change ≥ 1. (C) Pearson’s correlation analysis of ribosomal subunit proteins and isoprenol production. Red dots represent a control strain. (D) Correlation heatmap with hierarchical clustering of gene expression and isoprenol production. (E) Magnified view of the clustered heatmap from panel D (i). Magnified view of cluster (ii) is shown in Supplementary Fig. S11. Error bars represent standard deviation from three biological replicates.

Following improved isoprenol production from successful sgRNA downregulation, we then sought to maximize titers by downregulating multiple genes (Banerjee et al., 2020; Fang et al., 2021). Here, we selected PP_4188 as one of the target genes for multiplex CRISPRi downregulation and paired it with previously identified targets that had shown potential for improving isoprenol production (Fig. 4C-J). The results indicated that while the downregulation of these genes was successful, none of the engineered strains produced higher isoprenol titer than PP_4188 downregulation alone (Supplementary Fig. S9). A possible explanation for this outcome is that we observed reduced expression levels of genes encoding ribosomal subunits in the strain with singular PP_4188 downregulation (Fig. 5C, Supplementary Table S3, and Supplementary Fig. S10). An analysis using correlation heatmap and hierarchical clustering revealed that ribosomal subunits co-clustered with the RNA-binding protein Hfq and membrane transport proteins (Fig. 5D and E, Supplementary Fig. S11). This may suggest that PP_4188 downregulation triggers a cellular stress response or diverts resources away from ribosome biogenesis, likely due to disruptions in central metabolism (Hawkins et al., 2015; Liao et al., 2019). As ribosome biogenesis is critical for protein synthesis and cellular growth, its downregulation could limit the cell’s ability to support additional metabolic burdens. Therefore, further downregulation of multiple genes did not result in higher isoprenol production, possibly due to the compounded effects on cellular resource allocation and stress.

## 3. Discussion

The findings from this study underscore the transformative potential of computationally guided target gene selection in metabolic engineering, as demonstrated by the superior performance of FluxRETAP compared to heuristic-based approaches. By leveraging FluxRETAP to systematically identify gene targets for knockdown, we achieved significantly higher isoprenol titers in *Pseudomonas putida* KT2440. Notably, 46.4% of the genes identified by FluxRETAP resulted in improved isoprenol titers, compared to only 15.7% of those in the heuristic gene selection. The highest titer of 1469 mg/L, obtained by downregulating PP_4188, far surpassed the best titer of 958 mg/L from the heuristic group. These results demonstrate that computational approaches can uncover nonintuitive gene targets that significantly impact product yields and thereby provide a powerful alternative to traditional heuristic methods.

The development and successful implementation of VAMMPIRE represents an advancement in metabolic engineering and synthetic biology by offering a streamlined and efficient approach to constructing sgRNA arrays for CRISPRi. In doing so, the approach addresses several common challenges associated with multiplexed gene downregulation (Adiego-Pérez et al., 2019). Through its linker-based modular cloning strategy, VAMMPIRE simplifies the design and assembly process and facilitates the construction of up to five sgRNAs in a one-pot assembly with high accuracy, as demonstrated by the creation of 275 CRISPRi plasmids in this study. By using distinct promoters like the constitutive J23119 and the inducible P_nagAa_, VAMMPIRE ensures that sgRNAs are expressed efficiently while reducing the burden of dCas9 expression on host cells. The ease with which multiple sgRNAs were constructed and validated the 100% assembly accuracy highlights the robustness and versatility of the method.

The analysis of sgRNA arrays provided valuable insights into the potential context dependency of gene repression within multiplexed sgRNA constructs. Previous studies have reported that CRISPR arrays are often prone to variability in downregulation efficiency based on the position of individual sgRNAs within the array (Liao et al., 2019). In contrast, the results from this study demonstrated that VAMMPIRE’s use of independently transcribed sgRNAs largely mitigated this issue, leading to consistent gene repression across different sgRNA positions. This is particularly important in the context of metabolic engineering, where the reproducibility of gene knockdown is crucial for iterative Design-Build-Test-Learn (DBTL) cycles aimed at optimizing metabolic pathways.

While we anticipate that our evidence substantiates the effectiveness of FluxRETAP and VAMMPIRE for metabolic engineering, it is important to acknowledge certain limitations in our research. Despite the success of single-gene downregulation, the multiplexing CRISPRi arrays with PP_4188 did not yield further improvements in isoprenol titers. These results emphasize the complexity of metabolic engineering, where improving the production of a target compound often involves trade-offs in cellular growth and resource management. While FluxRETAP identified PP_4188 as a high-impact target for isoprenol production, further optimization of the strain may require strategies that mitigate cellular stress and improve resource allocation. For instance, tuning the level of PP_4188 downregulation could help optimize metabolic flux, reducing unintended disruptions to ribosome biogenesis and cellular fitness, potentially making it feasible to implement multiple, carefully calibrated gene downregulations.

In conclusion, this study demonstrates the effectiveness of FluxRETAP in identifying key gene targets for metabolic engineering, particularly for enhancing isoprenol production in *P. putida*. The downregulation of PP_4188 was especially impactful, revealing new insights into the relationship between central metabolism and product yields. The challenges encountered with multiplexed gene downregulation highlight the need for a deeper understanding of how cellular stress responses and resource allocation impact pathway flux. To address this complex issue, a strategy that integrates -omics data, metabolic modeling, and machine learning will be essential for further optimization of these complex metabolic networks to maximize production titers

## 4. Materials and Methods

### 4.1 Strains, plasmids, media, and growth conditions

*E. coli* XL1-Blue (Thermo Fisher Scientific) was used to propagate all plasmids in this study. The strain was grown in LB medium (Sigma Aldrich) at 37 °C, 180 rpm, with antibiotics (gentamicin 10 μg/mL, kanamycin 50 μg/mL). Plasmids were assembled using a modified BASIC method (Storch et al., 2015) and are listed in Supplementary Table S5.

Self replicating plasmids (RK2 and RSF1010-based plasmids) were transformed into *Pseudomonas putida* KT2440 by electroporation following the previous study (Yunus et al., 2024). Briefly, a fresh P. putida KT2440 colony was grown in 5 mL LB overnight at 30 °C, 200 rpm. The culture was centrifuged (1 min at 13,000 x g), washed three times with 1 mL of 10% glycerol, and resuspended in 500 μL of 10% glycerol at room temperature. For electroporation, ∼100 ng of DNA was added to 100 μL of cells and pulsed using a Bio-Rad GenePulser II (1.8 kV, 200 Ω, 1 mm cuvette). Afterward, 1 mL LB was added, and the mixture was transferred to a microtube and incubated at 30 °C, 200 rpm for 1 hour. Finally, 20 μL was plated on selective agar with antibiotic and incubated overnight at 30 °C.

*P. putida* KT2440 seed cultures were grown from single colonies in 5 mL LB at 30 °C, 180 rpm, overnight. Unless specified, 100 μL of the overnight culture was transferred to 5 mL M9 minimal medium and incubated under the same conditions overnight. This step was repeated once to adapt the cells to M9 medium. For isoprenol production, cultures were inoculated at an OD_600_ of 0.2 in 5 mL M9 medium with 2% glucose and antibiotics (kanamycin 50 mg/L, gentamicin 10 mg/L) and induced with 0.2% L-arabinose 4 hours after inoculation.

### 4.2 Routine isoprenol extraction and analysis

To analyze isoprenol, 100 μL of liquid culture was mixed with 400 μL of ethyl acetate, vortexed for 5 minutes at 3,000 rpm, and centrifuged for 1 minute at 13,000 x g. Isoprenol was sampled from the top ethyl acetate layer and measured using GC-FID, with concentration determined using serially diluted isoprenol standards.

### 4.3 Construction of CRISPRi plasmids

For CRISPRi-mediated gene downregulation, single guide RNAs (sgRNAs) with 22 nucleotides were designed using the web tool CRISPOR (Concordet and Haeussler, 2018) to target the non-template strand with 3’-NGG-5’ protospacer adjacent motif (PAM) sequence. A pair of complementary DNA oligos was ordered from Integrated DNA Technologies, hybridized at 95°C for 10 min, and cooled down to room temperature in a heating block. Hybridized oligo was digested with *Bsa*I (New England Biolabs) and ligated to *PaqC*I-digested pIY993 plasmid (JBx_249165). The resulting plasmid was digested with *Bsa*I and cloned into pIY989 plasmid (JBx_249567). Hybridized oligos used for CRISPRi-mediated gene downregulation are listed in Supplementary Table S6. Detail protocol for plasmid construction is described in Supplementary Note 2.

### 4.4 FluxRETAP

The source code of FluxRETAP is available under an open-source license and can be accessed at https://github.com/JBEI/FluxRETAP (see license details herein).

### 4.5 Fluorescence measurement

Fluorescence measurements were taken using a flow cytometer based on a previous study (Yunus et al., 2024). To start, 2 mL of LB medium was inoculated with 0.1% (v/v) overnight culture and supplemented with adequate antibiotics. After 24 hours, 1-3 μL of the sample was added to 150 μL of 1X phosphate-buffered saline. Single-cell RFP and GFP fluorescence from at least 30,000 cells was recorded using a BD C6 Accuri flow cytometer. GFP and RFP fluorescence were measured with FL1 and FL4 detectors, respectively. Protein fluorescence levels were determined by averaging the fluorescence distribution.

### 4.6 Proteomics analysis

Protein was extracted from cell pellets and tryptic peptides were prepared by following established proteomic sample preparation protocol (Chen et al., 2023). Cell pellets were resuspended in Qiagen P2 Lysis Buffer to lyse cells. Proteins were precipitated by adding 1 mM NaCl and 4 times the volume of acetone, followed by two washes with 80% acetone in water. The protein pellet was homogenized with 100 mM ammonium bicarbonate in 20% methanol. Protein concentration was measured using the DC protein assay (BioRad). Reduction was done with 5 mM TCEP (tris 2-(carboxyethyl) phosphine) for 30 min at room temperature, and alkylation with 10 mM iodoacetamide for 30 min in the dark. Proteins were digested overnight with trypsin at a 1:50 ratio. The resulting peptides were analyzed on an Agilent 1290 UHPLC system coupled to a Thermo Orbitrap Exploris 480 mass spectrometer (Chen et al., 2020).

Peptide samples were loaded onto an Ascentis^®^ ES-C18 Column and eluted with a 10-minute gradient from 98% solvent A (0.1% FA in H_2_O) and 2% solvent B (0.1% FA in ACN) to 65% solvent A and 35% solvent B. Eluting peptides were introduced to the mass spectrometer in positive-ion mode and measured in data-independent acquisition (DIA) mode, with 3 survey scans (m/z 380–985) and 45 MS2 scans. DIA raw data were analyzed using the DIA-NN software. (Demichev et al., 2020). The DIA-NN search used the latest *P. putida* KT2440 Uniprot proteome FASTA sequences, along with sequences for heterologous proteins and common contaminants. DIA-NN automatically determined mass tolerances and optimal mass accuracies based on initial sample analysis. The retention time extraction window was optimized for each MS run. Protein inference was enabled, and quantification used the Robust LC = High Accuracy method. The main DIA-NN reports were filtered with a global FDR = 0.01 at both the precursor and protein group levels. Protein quantities were plotted using the Top3 method, which averages the MS signal of the three most intense tryptic peptides. (Ahrné et al., 2013; Silva et al., 2006). The mass spectrometry proteomics data generated have been deposited to the ProteomeXchange Consortium via the PRIDE partner repository with the dataset identifier PXD062697 (Perez-Riverol et al., 2022).

### 4.7 Metabolomics analysis

Measurement of metabolites in the TCA cycle was conducted as described in a previous study (Banerjee et al., 2025).

## 5. Conclusion

In this study, we present an integrated strategy for improving isoprenol production in *P. putida* that combines predictive modeling with modular CRISPRi tools. FluxRETAP effectively prioritized gene knockdown targets that enhanced isoprenol production, while VAMMPIRE enabled easy, accurate, and position-independent assembly of CRISPRi constructs. Our observation that the guide RNA position had minimal influence on gene repression expands the utility of CRISPRi arrays in multiplex applications. Together, these tools provide a streamlined and reliable pipeline for testing gene perturbations at scale. The combination of predictive modeling and positionally robust gene knockdowns paves the way for more efficient DBTL cycles and the rational engineering of high-performing microbial cell factories.

## Acknowledgements

This work was supported by the US Department of Energy, Office of Science, Office of Biological and Environmental Research, through Contract DE-AC0205CH11231 between Lawrence Berkeley National Laboratory and the US Department of Energy.

## Notes

### Competing Interest Statement

The authors have declared no competing interest.

