## Supporting Information for "Predictive genome-wide CRISPR-mediated gene downregulation for enhanced bioproduction"

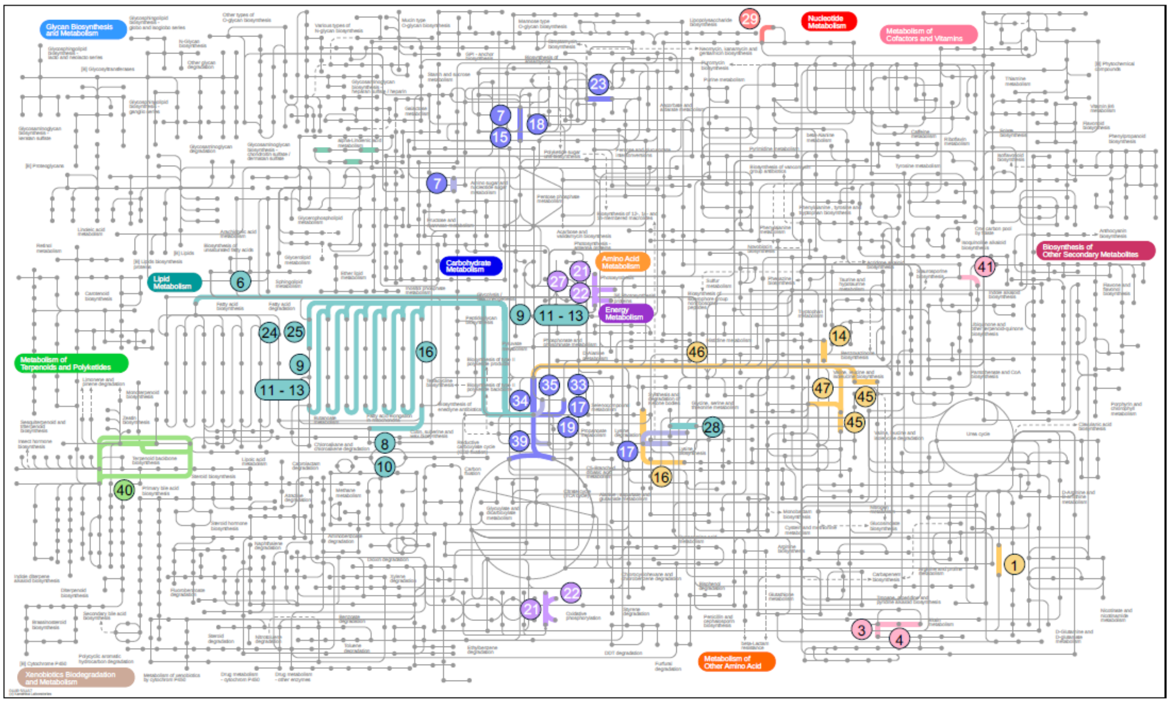


**Amino acid metabolism**

| | | | |
|---|---|---|---|
| 1 | PP_2589 | | Aldehyde dehydrogenase family protein |
| 14 | PP_5064 | BetA-II | choline dehydrogenase |
| 16 | PP_2137 | PcaF-II | eta-ketoadipyl CoA thiolase, beta component |
| 45 | PP_4678 | IlvC | Ketol-acid reductoisomerase |
| 46 | PP_1157 | | Acetolactate synthase |
| 47 | PP_4679 | IlvH | acetohydroxybutanoate synthase / acetolactate synthase |

**Metabolism of cofactors and vitamins**

| | | | |
|---|---|---|---|
| 3 | PP_0362 | BioB | Biotin synthase |
| 4 | PP_0437 | BirA | bifunctional biotin-[acetyl-CoA-carboxylase] ligase/biotin operon repressor BirA |
| 41 | PP_0548 | PadA1 | putative phenylacrylic acid decarboxylase |

**Carbohydrate metabolism**

| | | | |
|---|---|---|---|
| 7 | PP_5288 | AlgC | Phosphomannomutase/phosphoglucomutase |
| 15 | PP_3578 | Pgm | phosphoglucomutase |
| 17 | PP_4487 | AcsA-I | acetyl-CoA synthetase |
| 18 | PP_1777 | CpsG | phosphomannomutase |
| 19 | PP_2213 | | putative acyl-CoA synthetase |
| 23 | PP_1444 | Gcd | quinoprotein glucose dehydrogenase |
| 33 | PP_4702 | AcsA-II | acetyl-CoA synthetase |
| 34 | PP_0338 | AceF | AceF-S-acetyldihydrolipoate |
| 35 | PP_0774 | Pta | Phosphate acetyltransferase |
| 39 | PP_4194 | GltA | citrate synthase |

**Energy metabolism**

| | | | |
|---|---|---|---|
| 21 | PP_5417 | AtpF | ATP synthase subunit b |
| 22 | PP_5416 | AtpH | ATP synthase subunit delta |
| 27 | PP_5419 | AtpB | ATP synthase subunit a |

**Nucleotide metabolism**

| | | | |
|---|---|---|---|
| 29 | PP_5335 | PurK | N5-carboxyaminoimidazole ribonucleotide synthase |

**Lipid metabolism**

| | | | |
|---|---|---|---|
| 6 | PP_1607 | AccA | acetyl-coenzyme A carboxylase carboxyl transferase |
| 8 | PP_4549 | FadD-I | long-chain-fatty-acid/CoA ligase |
| 9 | PP_4636 | YqeF | acetyl-CoA acetyltransferase |
| 10 | PP_4550 | FadD-II | long-chain-fatty-acid/CoA ligase |
| 11 | PP_3754 | BktB | Beta-ketothiolase |
| 12 | PP_3355 | | Beta-ketothiolase |
| 13 | PP_0582 | | Thiolase family protein |
| 16 | PP_2137 | PcaF-II | eta-ketoadipyl CoA thiolase, beta component |
| 24 | PP_3755 | Hbd | 3-hydroxybutyryl-CoA dehydrogenase |
| 25 | PP_3282 | PaaH | 3-hydroxyadipyl-CoA dehydrogenase |
| 28 | PP_3394 | | putative 3-hydroxy-3-methylglutaryl-CoA lyase |

Supplementary Figure S1. Metabolic map depicting heuristic target genes for CRISPRi downregulation. (https://pathways.embl.de/) (Darzi et al., 2018). KEGG Orthology alphanumerical code for each gene was used as an input in element selection. A tabulated list of CRISPRi heuristic target genes can be found in Supplementary Table S1.

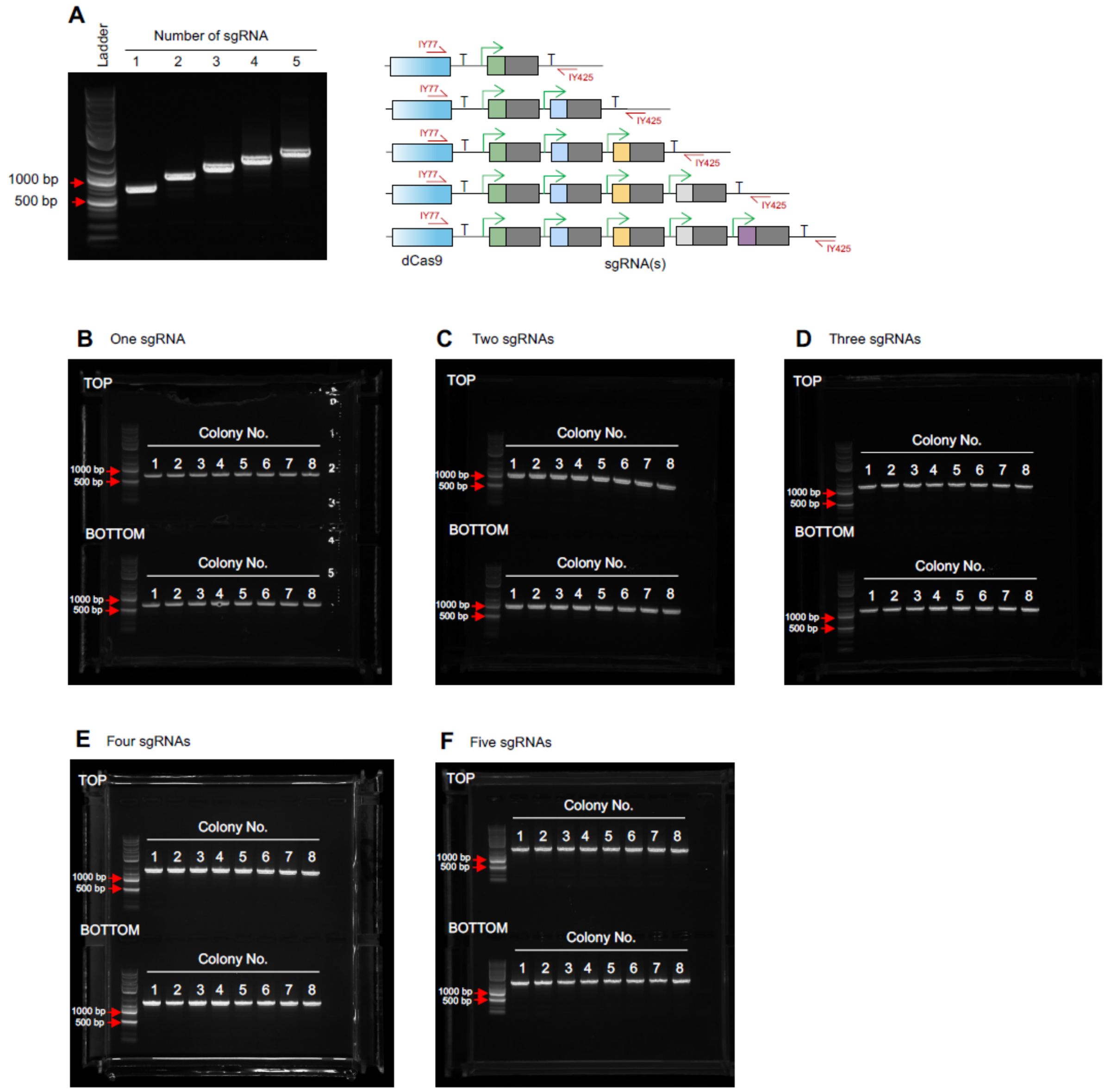


Supplementary Figure S2. Assessment of cloning accuracy. (A) Gel images showing sequence-verified plasmids harboring one, two, three, four, and five sgRNAs. (B-F) Gel images of colonies carrying different number of sgRNAs. For each gel, two sets of eight colonies, each containing one of two different plasmid constructs (depicted at the top and bottom of the gels), were assessed. All results were obtained through PCR amplification using primers IY77 and IY425

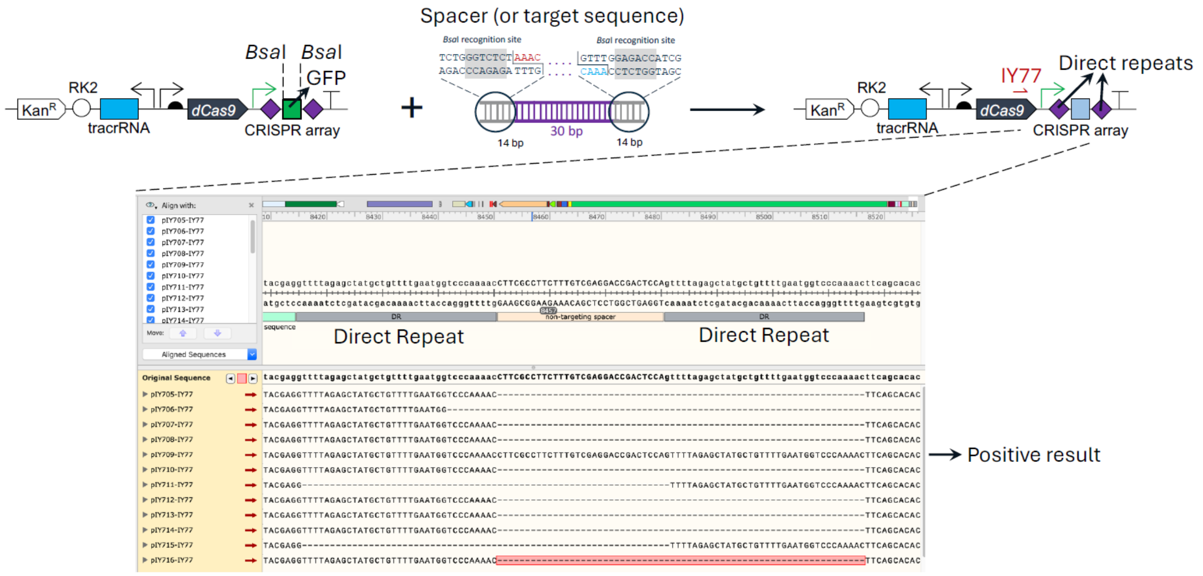


Supplementary Figure S3. Cloning of CRISPRi array using Golden Gate method (Aparicio et al., 2019). Plasmid and spacer were both digested with *Bsa*I. Subsequently, digested spacer was ligated to the plasmid. Sanger sequencing using primer IY77 revealed that fewer than 5% of the plasmids were correctly assembled. Incorrectly assembled plasmids exhibited homologous recombination of the direct repeats.

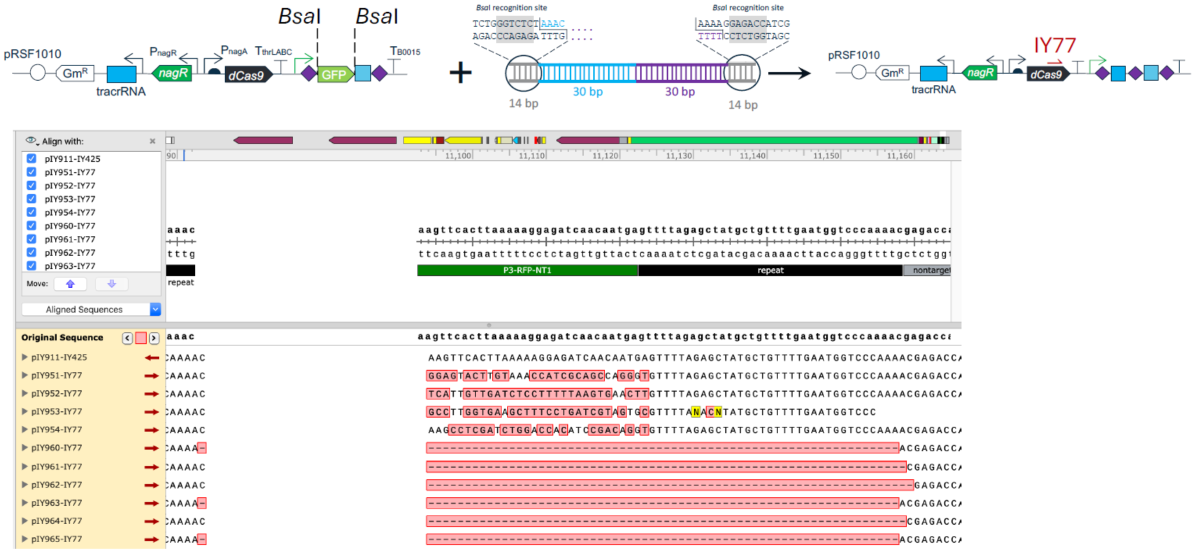


Supplementary Figure S4. Cloning of CRISPRi array using CRISPathBrick method (Cress et al., 2015). Sequencing results revealed that approximately 45% (5 out of 11) constructs were correctly cloned.

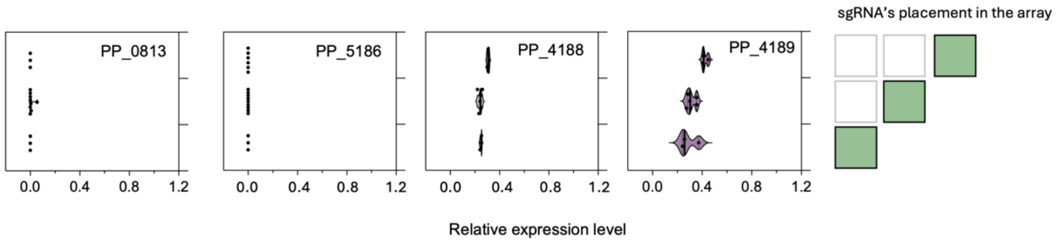


Supplementary Figure S5. Relative expression levels of PP_0813, PP_5186, PP_4188, and PP_4189 genes when the sgRNA targeting that specific gene is placed at different position in the array. All samples were extracted at 48 hr.

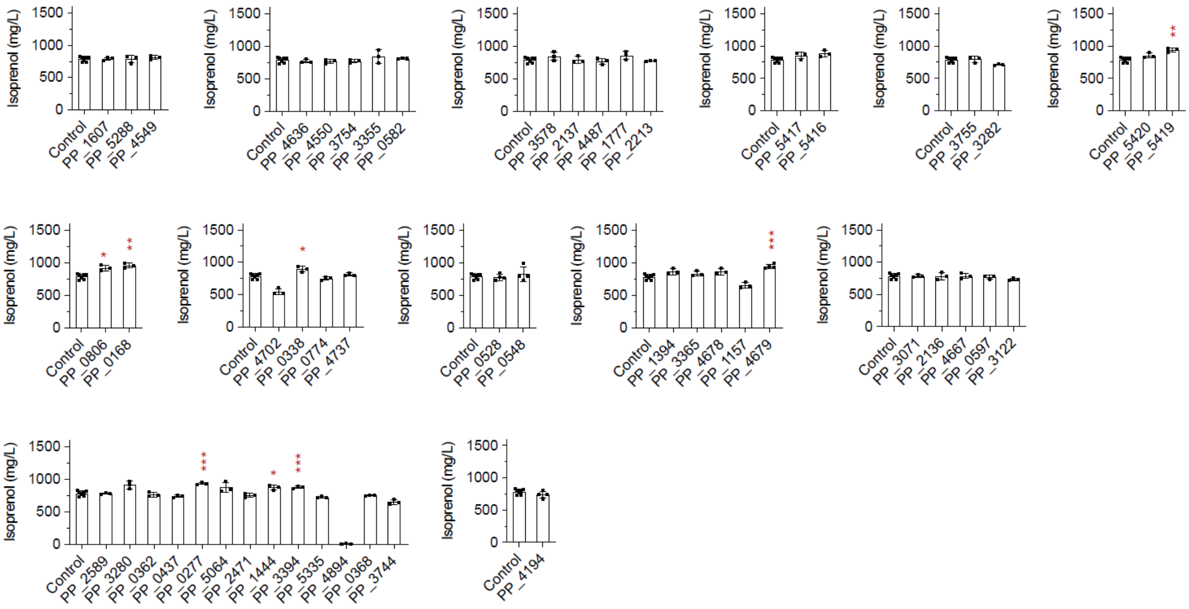


Supplementary Figure S6. Isoprenol titers obtained from heuristic target genes. All samples were extracted at 48 hr. Error bars represent standard deviation from 3 to 6 biological replicates.

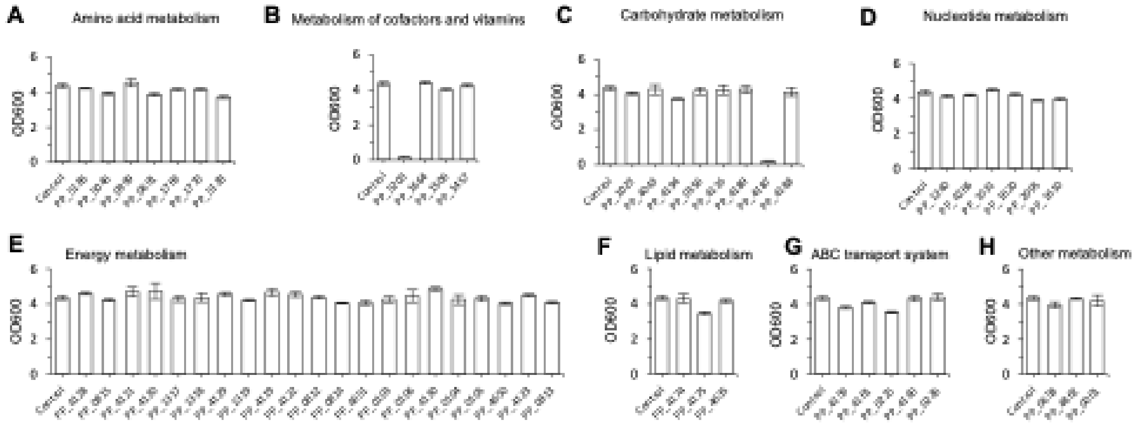


Supplementary Figure S7. $OD_{600}$ of strains shown in Fig. 4. $OD_{600}$ was sampled at 48 hr. Error bars represent standard deviation from 3 to 6 biological replicates.

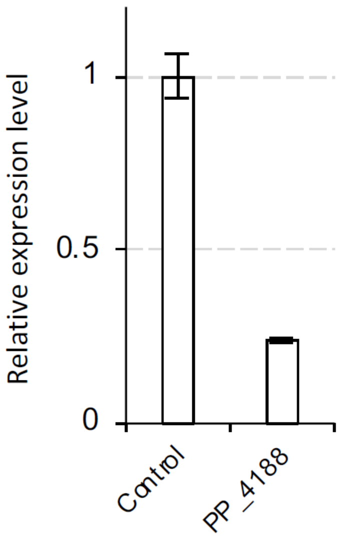


Supplementary Figure S8. Relative expression level of PP_4188 gene in the control and PP_4188 strains. Proteins were extracted at 48 h. Error bars represent standard deviation from three biological replicates.

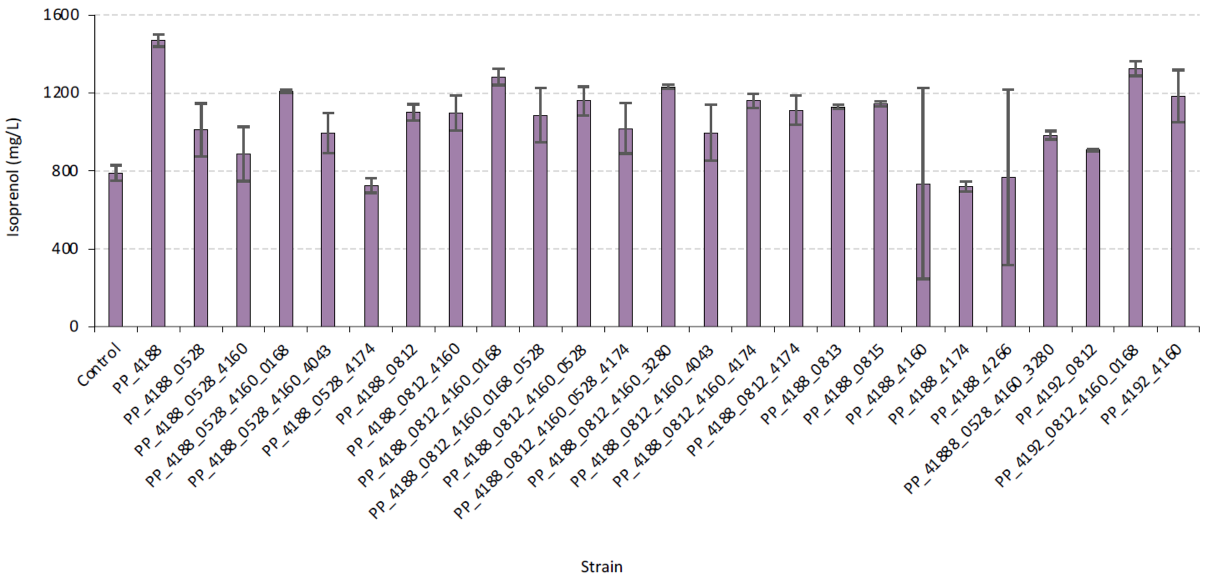


Supplementary Figure S9. Isoprenol production from strains carrying multiple sgRNAs. All samples were extracted at 48 hr. Error bars represent standard deviation from 3~6 biological replicates.

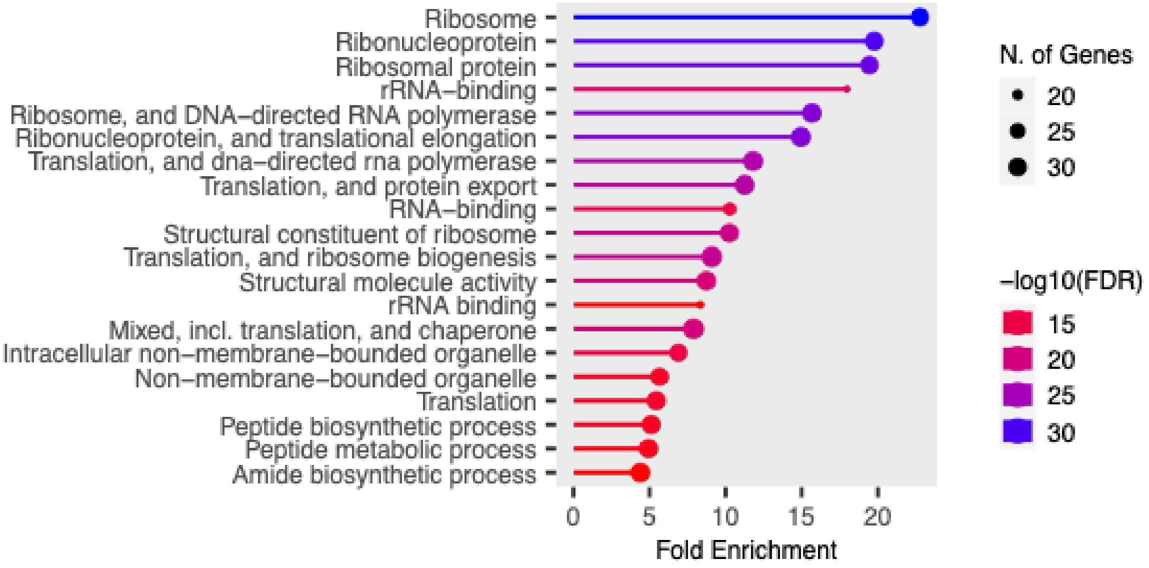


Figure S10. Gene enrichment analysis of PP_4188 strain. A list of significantly downregulated genes obtained from Supplementary Table 3 was used as an input for gene enrichment analysis. Gene enrichment analysis was performed using ShinyGO 0.82 (Ge et al., 2020) with false discovery rate (FDR) cutoff 0.05. The result indicates enrichment of many ribosome-related proteins which suggests that translation machinery is downregulated.

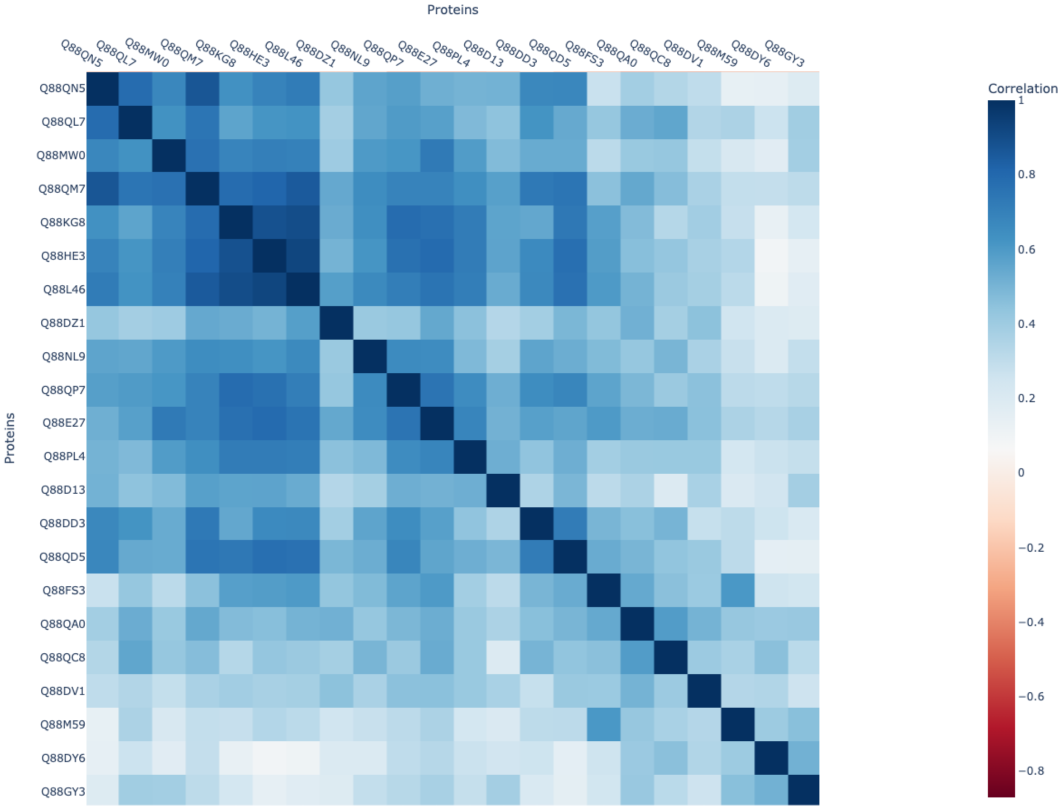


Supplementary Figure S11. A correlation heatmap with hierarchical clustering (Figure 5D, (ii)) showing clustered of ribosomal subunit proteins, a small heat shock protein, and membrane transport proteins with high correlation values. Ribosomal subunit proteins (Q88QN5, Q88QM7, Q88QL7), RNA-binding protein Hfq (Q88DD3), membrane transport proteins (Q88D13, Q88PL4, Q88E27, Q88QP7, Q88NL9, Q88L46, D88HE3, Q88KG8, Q88MW0).

Supplementary Table S1. List of CRISPRi heuristic target genes

| No | Target Gene | Enzyme | Enzyme description | KEGG Orthology | Metabolic Pathway | iPath3 |
|---|---|---|---|---|---|---|
| 1 | PP_2589 | | Aldehyde dehydrogenase family protein | K09472 | Arginine and proline metabolism | |
| 2 | PP_3280 | PaaJ | 3-oxoadipyl-CoA/3-oxo-5,6-dehydrosuberyl-CoA thiolase | K07823 | Benzoate degradation paaJ | NOT FOUND |
| 3 | PP_0362 | BioB | Biotin synthase | K01012 | Biotin metabolism | |
| 4 | PP_0437 | BirA | bifunctional biotin-[acetyl-CoA-carboxylase] ligase/biotin operon repressor BirA | K03524 | BirA family transcriptional regulator, biotin operon repressor | |
| 5 | PP_0277 | | conserved exported protein of unknown function | NOT AVAILABLE | conserved exported protein of unknown function | NOT FOUND |
| 6 | PP_1607 | AccA | acetyl-coenzyme A carboxylase carboxyl transferase subunit alpha | K01962 | Fatty acid biosynthesis | |
| 7 | PP_5288 | AlgC | Phosphomannomutase/phosphoglucomutase | K15778 | Fatty acid biosynthesis | |
| 8 | PP_4549 | FadD-I | long-chain-fatty-acid/CoA ligase | K01897 | Fatty acid biosynthesis | |
| 9 | PP_4636 | YqeF | acetyl-CoA acetyltransferase | K00626 | Fatty acid degradation | |
| 10 | PP_4550 | FadD-II | long-chain-fatty-acid/CoA ligase | K01897 | Fatty acid degradation | |
| 11 | PP_3754 | BktB | Beta-ketothiolase | K00626 | Fatty acid degradation | |
| 12 | PP_3355 | | Beta-ketothiolase | K00626 | Fatty acid degradation | |
| 13 | PP_0582 | | Thiolase family protein | K00626 | Fatty acid degradation | |
| 14 | PP_5064 | BetA-II | choline dehydrogenase | K00108 | Glycine, serine and threonine metabolism | |
| 15 | PP_3578 | Pgm | phosphoglucomutase | K01835 | Glycolysis / Gluconeogenesis | |
| 16 | PP_2137 | PcaF-II | eta-ketoadipyl CoA thiolase, beta component | K00632 | Glycolysis / Gluconeogenesis | |
| 17 | PP_4487 | AcsA-I | acetyl-CoA synthetase | K01895 | Glycolysis / Gluconeogenesis | |
| 18 | PP_1777 | CpsG | phosphomannomutase | K15778 | Glycolysis / Gluconeogenesis | |
| 19 | PP_2213 | | putative acyl-CoA synthetase | K01895 | Glycolysis / Gluconeogenesis | |
| 20 | PP_2471 | IhfA | Integration host factor subunit alpha | K04764 | Integration host factor subunit alpha | NOT FOUND |
| 21 | PP_5417 | AtpF | ATP synthase subunit b | K02109 | Oxidative phosphorylation | |
| 22 | PP_5416 | AtpH | ATP synthase subunit delta | K02113 | Oxidative phosphorylation | |
| 23 | PP_1444 | Gcd | quinoprotein glucose dehydrogenase | K00117 | Pentose phosphate pathway | |
| 24 | PP_3755 | Hbd | 3-hydroxybutyryl-CoA dehydrogenase | K00074 | Phenylalanine metabolism | |
| 25 | PP_3282 | PaaH | 3-hydroxyadipyl-CoA dehydrogenase | K00074 | Phenylalanine metabolism | |
| 26 | PP_5420 | AtpI | ATP synthase protein I | K02116 | Photosystem and electron transport system | NOT FOUND |
| 27 | PP_5419 | AtpB | ATP synthase subunit a | K02108 | Photosystem and electron transport system | |

| | | | | | | |
|---|---|---|---|---|---|---|
| 28 | PP_3394 | | putative 3-hydroxy-3-methylglutaryl-CoA lyase | K01640 | Pinene, camphor and geraniol degradation | |
| 29 | PP_5335 | PurK | N5-carboxyaminoimidazole ribonucleotide synthase | K01589 | Purine metabolism | |
| 30 | PP_3123 | AtoB | acetoacetyl CoA-transferase (subunit B) | K01029 | putative Polysaccharide transporter | NOT FOUND |
| 31 | PP_0806 | | utative Surface adhesion protein | K20276 | putative Surface adhesion protein | NOT FOUND |
| 32 | PP_0168 | | putative surface adhesion protein | K12549 | putative surface adhesion protein | NOT FOUND |
| 33 | PP_4702 | AcsA-II | acetyl-CoA synthetase | K01895 | Pyruvate metabolism | |
| 34 | PP_0338 | AceF | AceF-S-acetyldihydrolipoate | K00627 | Pyruvate metabolism | |
| 35 | PP_0774 | Pta | Phosphate acetyltransferase | K13788 | Pyruvate metabolism | |
| 36 | PP_4737 | Dld2 | D-lactate dehydrogenase | K18930 | Pyruvate metabolism dld2 | NOT FOUND |
| 37 | PP_4894 | Hfq | RNA-binding protein Hfq | K03666 | Quorum sensing hfq | NOT FOUND |
| 38 | PP_0368 | | putative Acyl-CoA dehydrogenase | K20035 | Sulfur metabolism | NOT FOUND |
| 39 | PP_4194 | GltA | citrate synthase | K01647 | TCA cycle | |
| 40 | PP_0528 | IspA | farnesyl diphosphate synthase | K13789 | Terpenoid backbone biosynthesis | |
| 41 | PP_0548 | PadA1 | putative phenylacrylic acid decarboxylase | K03186 | Terpenoid backbone biosynthesis | |
| 42 | PP_3744 | GlcC | transcriptional dual regulator GlcC-Glycolate | K11474 | Transcriptional dual regulator GlcC-Glycolate | NOT FOUND |
| 43 | PP_1394 | | putative Acetolactate synthase, large subunit | K01652 | Valine, leucine and isoleucine biosynthesis | NOT FOUND |
| 44 | PP_3365 | | Acetolactate synthase | K01652 | Valine, leucine and isoleucine biosynthesis | NOT FOUND |
| 45 | PP_4678 | IlvC | Ketol-acid reductoisomerase | K00053 | Valine, leucine and isoleucine biosynthesis | |
| 46 | PP_1157 | | Acetolactate synthase | K01652 | Valine, leucine and isoleucine biosynthesis | |
| 47 | PP_4679 | IlvH | acetohydroxybutanoate synthase / acetolactate synthase | K01653 | Valine, leucine and isoleucine biosynthesis | |
| 48 | PP_3071 | Aacs | acetoacetyl-coenzyme A synthetase | K01907 | Valine, leucine and isoleucine degradation | NOT FOUND |
| 49 | PP_2136 | FadB | enoyl-CoA hydratase/3-hydroxyacyl-CoA dehydrogenase | K01825 | Valine, leucine and isoleucine degradation | NOT FOUND |
| 50 | PP_4667 | MmsA-II | methylmalonate-semialdehyde dehydrogenase | K00140 | Valine, leucine and isoleucine degradation | NOT FOUND |
| 51 | PP_0597 | MmsA-I | methylmalonate-semialdehyde dehydrogenase | PP_0597 | Valine, leucine and isoleucine degradation | NOT FOUND |
| 52 | PP_3122 | AtoA | acetoacetyl CoA-transferase (subunit A) | K01028 | Valine, leucine and isoleucine degradation | NOT FOUND |

Supplementary Table S2. List of CRISPRi FluxRETAP target genes

| No. | Target Gene | Enzyme | Enzyme description | KEGG Orthology | Metabolic pathways | iPath3 |
|---|---|---|---|---|---|---|
| 1 | PP_5186 | ArgE | acetylornithine deacetylase | K01438 | Arginine and proline metabolism | |
| 2 | PP_5203 | Fau | 5-formyltetrahydrofolate cyclo-ligase | K01934 | Cofactor and prosthetic group biosynthesis | |
| 3 | PP_0658 | | Homocysteine S-methyltransferase family protein | NOT AVAILABLE | Cysteine metabolism | NOT FOUND |
| 4 | PP_4174 | FabA | 3R-3-hydroxydecanoyl acyl carrier protein (ACP) dehydratase | K01716 | Fatty acid biosynthesis | |
| 5 | PP_4175 | FabB | 3-oxoacyl-[acyl-carrier-protein] synthase 1 | K00647 | Fatty acid biosynthesis | |
| 6 | PP_4635 | | trans-2-enoyl-CoA reductase (NAD(+)) | K00209 | Fatty acid biosynthesis | |
| 7 | PP_5046 | GlnA | glutamine synthetase | K01915 | Glutamate metabolism | |
| 8 | PP_0999 | ArcC | carbamate kinase | K00926 | Glutamate metabolism | |
| 9 | PP_0691 | ProB | Glutamate 5-kinase | K00931 | Glutamate metabolism | |
| 10 | PP_4862 | YdfG | 3-hydroxy acid dehydrogenase, NADP-dependent / malonic semialdehyde reductase | NOT AVAILABLE | Glycine, serine, threonine metabolism | NOT FOUND |
| 11 | PP_1023 | Pgl | 6-phosphogluconolactonase | K01057 | Glycolysis | |
| 12 | PP_4128 | NuoK | NADH-quinone oxidoreductase subunit K | K00340 | Oxydative phosphorylation | |
| 13 | PP_0815 | CyoD | cytochrome bo terminal oxidase subunit IV | K02300 | Oxydative phosphorylation | |
| 14 | PP_4121 | NuoC | NADH-quinone oxidoreductase subunit C/D | K13378 | Oxydative phosphorylation | |
| 15 | PP_4120 | NuoB | NADH-quinone oxidoreductase subunit B | K00331 | Oxydative phosphorylation | |
| 16 | PP_1317 | PetA | Ubiquinol-cytochrome c reductase iron-sulfur subunit | K00411 | Oxydative phosphorylation | |
| 17 | PP_1318 | PetB | Cytochrome b | K00412 | Oxydative phosphorylation | |
| 18 | PP_4129 | NuoL | NADH:ubiquinone oxidoreductase, membrane subunit L | K00341 | Oxydative phosphorylation | |
| 19 | PP_1319 | PetC | Ubiquinol--cytochrome c reductase, cytochrome c1 | K00413 | Oxydative phosphorylation | |
| 20 | PP_4119 | NuoA | NADH-quinone oxidoreductase subunit A | K00330 | Oxydative phosphorylation | |
| 21 | PP_4122 | NuoE | proton-pumping NADH:ubiquinone oxidoreductase, respiratory complex I, subunit E | K00334 | Oxydative phosphorylation | |
| 22 | PP_0812 | CyoA | cytochrome bo terminal oxidase subunit II | K02297 | Oxydative phosphorylation | |
| 23 | PP_0814 | CyoC | ytochrome bo terminal oxidase subunit III | K02299 | Oxydative phosphorylation | |
| 24 | PP_4651 | CioA | Ubiquinol oxidase subunit I, cyanide insensitive | K00425 | Oxydative phosphorylation | |
| 25 | PP_0103 | | cytochrome c oxidase subunit 2 | K02275 | Oxydative phosphorylation | |

| | | | | | | |
|---|---|---|---|---|---|---|
| 26 | PP_0106 | | cytochrome c oxidase subunit 3 | K02276 | Oxydative phosphorylation | |
| 27 | PP_4130 | NuoM | NADH:ubiquinone oxidoreductase, membrane subunit M | K00342 | Oxydative phosphorylation | |
| 28 | PP_0104 | CtaD | Cytochrome c oxidase subunit 1 | K02274 | Oxydative phosphorylation | |
| 29 | PP_0105 | | Cytochrome C oxidase assembly protein | K02258 | Oxydative phosphorylation | |
| 30 | PP_4650 | CioB | Ubiquinol oxidase subunit II, cyanide insensitive | K00426 | Oxydative phosphorylation | |
| 31 | PP_4123 | NuoF | NADH-quinone oxidoreductase subunit F | K00335 | Oxydative phosphorylation | |
| 32 | PP_0813 | CyoB | cytochrome bo terminal oxidase subunit I | K02298 | Oxydative phosphorylation | |
| 33 | PP_4043 | GntZ | utative 6-phosphogluconate dehydrogenase, decarboxylating | K00033 | Pentose phosphate pathway | |
| 34 | PP_1769 | PheA | chorismate mutase/Prephenate dehydratase | K14170 | Phenylalanine, tyrosine, tryptophan biosynthesis | |
| 35 | PP_1770 | | 3-phosphoshikimate 1-carboxyvinyltransferase | K24018 | Phenylalanine, tyrosine, tryptophan biosynthesis | |
| 36 | PP_0001 | ParB | probable chromosome-partitioning protein | K03497 | Probable-partitioning protein | NOT FOUND |
| 37 | PP_1664 | PurN | phosphoribosylglycinamide formyltransferase 1 | K11175 | Purine metabolism | |
| 38 | PP_1240 | PurC | Phosphoribosylaminoimidazole-succinocarboxamide synthase | K01923 | Purine metabolism | |
| 39 | PP_1506 | Adk | Adenylate kinase | K00939 | Purine metabolism | |
| 40 | PP_1457 | PurT | Phosphoribosylglycinamide formyltransferase 2 | K08289 | Purine metabolism | |
| 41 | PP_4266 | Apt | Adenine phosphoribosyltransferase | K00759 | Purine metabolism | |
| 42 | PP_1031 | GuaB | IMP dehydrogenase and single strand DNA binding factor | K00088 | Purine metabolism | |
| 43 | PP_1620 | SurE | 5'(3')-nucleotidase and polyphosphatase | K03787 | Pyrimidine metabolism | |
| 44 | PP_2095 | PyrD | Dihydroorotate dehydrogenase (quinone) | K00254 | Pyrimidine metabolism | |
| 45 | PP_1610 | PyrG | CTP synthase | K01937 | Pyrimidine metabolism | |
| 46 | PP_4194 | GltA | citrate synthase | K01647 | TCA cycle | |
| 47 | PP_0356 | GlcB | malate synthase G | K01638 | TCA cycle | |
| 48 | PP_4116 | AceA | isocitrate lyase | K01637 | TCA cycle | |
| 49 | PP_4189 | SucA | 2-oxoglutarate decarboxylase, thiamine-requiring E1 subunit | K00164 | TCA cycle | |
| 50 | PP_4187 | LdpG | dihydrolipoyl dehydrogenase | K00382 | TCA cycle | |
| 51 | PP_4188 | SucB | 2-oxoglutarate dehydrogenase dihydrolipoyltranssuccinylase subunit | K00658 | TCA cycle | |
| 52 | PP_4159 | KdpC | Potassium-transporting ATPase C chain | K01548 | Transport ABC system | NOT FOUND |
| 53 | PP_4161 | KdpA | potassium-transporting ATPase A chain | K01546 | Transport ABC system | NOT FOUND |

| | | | | | | |
|---|---|---|---|---|---|---|
| 54 | PP_0225 | SctC | Sulfur compound ABC transporter - ATP-binding subunit | K10010 | Transport ABC system | NOT FOUND |
| 55 | PP_4160 | KdpB | K+ transporting ATPase, KdpB subunit | K01547 | Transport ABC system | NOT FOUND |
| 56 | PP_0226 | SctS | sulfur compound ABC transporter - permease subunit | K10009 | Transport ABC system | NOT FOUND |
| 57 | PP_5128 | IlvD | Dihydroxy-acid dehydratase | K01687 | Valine, leucine, and isoleucine metabolism | |

Supplementary Table S3. List of downregulated genes from PP_4188 strain

| Protein Group | Protein Names | Protein | Protein Description | Fold Change | Log2(Fold Change) | P-Value (Equal Variance) | (-Log10(P-Value)) | Rank |
|---|---|---|---|---|---|---|---|---|
| Q88K38 | Q88K38_PSEPK | Rbsb | Ribose ABC transporter, periplasmic ribose-binding protein | 0.003286423 | -8.249266263 | 0.001195825 | 2.922332234 | 1 |
| Q88MY2 | PDXJ_PSEPK | Pdxj | Pyridoxine 5'-phosphate synthase | 0.018011482 | -5.794939331 | 1.30E-06 | 5.884647992 | 2 |
| Q88E40 | Q88E40_PSEPK | Mioc | MioC protein | 0.042452489 | -4.558007043 | 0.000162191 | 3.789974356 | 3 |
| Q88H07 | Q88H07_PSEPK | Pp_3558 | Amino acid transporter, periplasmic amino acid-binding protein, putative | 0.044852591 | -4.478664853 | 0.000898422 | 3.046519738 | 4 |
| Q88ES6 | Q88ES6_PSEPK | Pp_4377 | Flagellin FlaG, putative | 0.045179862 | -4.468176337 | 3.13E-05 | 4.504720107 | 5 |
| Q88RJ5 | Q88RJ5_PSEPK | Pp_0134 | Uncharacterized protein | 0.059121193 | -4.080180819 | 0.031998481 | 1.49487064 | 6 |
| Q88QI8 | Q88QI8_PSEPK | Oprg | Outer membrane protein OprG | 0.060034368 | -4.058067555 | 0.001073287 | 2.969283975 | 7 |
| Q88CB1 | DADA2_PSEPK | Dada2 | D-amino acid dehydrogenase 2 | 0.061015017 | -4.034691825 | 0.000254631 | 3.5940884 | 8 |
| Q88GW5 | Q88GW5_PSEPK | Pp_3602 | Ketoglutarate semialdehyde dehydrogenase | 0.066563747 | -3.909119533 | 0.017285891 | 1.762308238 | 9 |
| Q88QD4 | AROQ1_PSEPK | Aroq1 | 3-dehydroquinate dehydratase 1 | 0.069593149 | -3.844910898 | 0.002008452 | 2.697138494 | 10 |
| Q88MX8 | CMOB_PSEPK | Cmob | tRNA (mo5U34)-methyltransferase | 0.081272814 | -3.621083335 | 0.0044711 | 2.349585573 | 11 |
| Q88L83 | Q88L83_PSEPK | Pp_2052 | Hydrolase, haloacid dehalogenase-like family | 0.082300639 | -3.602952561 | 0.001109586 | 2.954838922 | 12 |
| Q88P65 | GCSP1_PSEPK | Gcvp1 | Glycine dehydrogenase (decarboxylating) 1 | 0.086920383 | -3.52416166 | 0.003748742 | 2.42611445 | 13 |
| Q88MP7 | Q88MP7_PSEPK | Pp_1523 | Uncharacterized protein | 0.106970312 | -3.224717638 | 0.000107187 | 3.969858356 | 14 |
| Q88K59 | Q88K59_PSEPK | Pp_2432 | Oxygen-insensitive NAD(P)H nitroreductase | 0.107208654 | -3.221506723 | 0.01053395 | 1.977408752 | 15 |
| Q88MF4 | Q88MF4_PSEPK | Pp_1617 | Esterase, putative | 0.112598663 | -3.150738401 | 0.003984412 | 2.399635791 | 16 |
| Q88HZ1 | Q88HZ1_PSEPK | Pp_3211 | ABC transporter, ATP-binding protein | 0.117200246 | -3.092952494 | 0.004371818 | 2.359337957 | 17 |
| Q88LR2 | Q88LR2_PSEPK | Pp_1867 | Uncharacterized protein | 0.121609268 | -3.039674916 | 0.032351585 | 1.49010444 | 18 |
| Q88C45 | Q88C45_PSEPK | Aspa | Aspartate ammonia-lyase | 0.122208367 | -3.032585027 | 0.001166225 | 2.933217699 | 19 |
| Q88KH2 | Q88KH2_PSEPK | Pp_2318 | Lipase, GDSL family | 0.12373818 | -3.01463738 | 0.006634097 | 2.178218162 | 20 |
| Q88NK1 | Q88NK1_PSEPK | Oprd | Porin D | 0.130819094 | -2.934354964 | 0.000381916 | 3.418031796 | 21 |
| Q88I97 | Q88I97_PSEPK | Pp_3102 | Uncharacterized protein | 0.134764917 | -2.891483123 | 0.014329483 | 1.843769465 | 22 |
| Q88CM3 | Q88CM3_PSEPK | Pp_5157 | Conserved domain protein | 0.137944148 | -2.857843839 | 0.004547514 | 2.342225918 | 23 |
| Q88CD2 | Q88CD2_PSEPK | Pp_5249 | Transcriptional regulator, LysR family | 0.146662159 | -2.769431414 | 0.005052473 | 2.296495984 | 24 |
| Q88ES8 | Q88ES8_PSEPK | Flis | Flagellar biosynthetis protein FliS | 0.14989376 | -2.737987769 | 0.00059732 | 3.223792858 | 25 |

| | | | | | | | | |
|---|---|---|---|---|---|---|---|---|
| Q88M32 | Q88M32_PSEPK | Pp_1743 | Sodium:solute symporter family protein | 0.152048629 | -2.717395283 | 0.001379257 | 2.860354877 | 26 |
| Q88K01 | Q88K01_PSEPK | Pp_2492 | Butanol dehydrogenase, NADH-dependent, putative | 0.160810119 | -2.636569901 | 0.017955209 | 1.745809533 | 27 |
| Q88MG9 | FABZ_PSEPK | Fabz | 3-hydroxyacyl-[acyl-carrier-protein] dehydratase FabZ | 0.161901 | -2.626816201 | 0.000439638 | 3.356904859 | 28 |
| Q88PD0 | Q88PD0_PSEPK | Xenb | Xenobiotic reductase B | 0.168886689 | -2.565872469 | 0.003313497 | 2.479713447 | 29 |
| Q88GK1 | Q88GK1_PSEPK | Pp_3720 | NAD(P)H quinone oxidoreductase, putative | 0.196711875 | -2.345844045 | 0.00188092 | 2.725629642 | 30 |
| Q88R88 | Q88R88_PSEPK | Pp_0245 | S1 RNA binding domain protein | 0.203003713 | -2.300421978 | 0.004499182 | 2.346866486 | 31 |
| P0A149 | Y002_PSEPK | Pp_0002 | Uncharacterized protein PP_0002 | 0.206393477 | -2.276530719 | 0.01652738 | 1.781795985 | 32 |
| Q88H64 | Q88H64_PSEPK | Pp_3497 | Peptidase, U32 family | 0.212307275 | -2.235774288 | 0.0429972 | 1.366559822 | 33 |
| Q88NX1 | Q88NX1_PSEPK | Bfr | Ferroxidase | 0.222272262 | -2.169600173 | 0.000557697 | 3.253601519 | 34 |
| Q88C38 | Q88C38_PSEPK | Pp_5345 | Acetyltransferase, GNAT family | 0.22535643 | -2.149719479 | 0.035951802 | 1.444279342 | 35 |
| Q88L46 | Q88L46_PSEPK | Oprf | Outer membrane protein OprF | 0.232724746 | -2.103303472 | 4.34E-05 | 4.36211881 | 36 |
| Q88RW7 | RECF_PSEPK | Recf | DNA replication and repair protein RecF | 0.235709625 | -2.084917421 | 0.034926755 | 1.456841767 | 37 |
| Q88RD1 | Q88RD1_PSEPK | Pp_0201 | Uncharacterized protein | 0.23595214 | -2.08343384 | 0.004495726 | 2.347200205 | 38 |
| Q88FA9 | Q88FA9_PSEPK | Kgda | 2-oxoglutarate dehydrogenase, E1 component | 0.237969085 | -2.071153935 | 1.08E-05 | 4.96716323 | 39 |
| Q88FB0 | Q88FB0_PSEPK | Kgdb | Dihydrolipoyllysine-residue succinyltransferase component of 2-oxoglutarate dehydrogenase complex | 0.238537433 | -2.067712413 | 2.79E-06 | 5.554287944 | 40 |
| Q88GL6 | Q88GL6_PSEPK | Pp_3705 | Uncharacterized protein | 0.240723236 | -2.054552692 | 0.019559565 | 1.708640819 | 41 |
| Q88P37 | Q88P37_PSEPK | Pp_1016 | Sugar ABC transporter, permease protein | 0.240899376 | -2.053497435 | 0.00064268 | 3.192005225 | 42 |
| Q88DT6 | SSRP_PSEPK | Smpb | SsrA-binding protein | 0.24122074 | -2.05157414 | 0.011839402 | 1.926670221 | 43 |
| Q88MD2 | TTCA_PSEPK | Ttca | tRNA 2-thiocytidine biosynthesis protein TtcA | 0.24350956 | -2.037949684 | 0.010392678 | 1.98327253 | 44 |
| Q88QC0 | Q88QC0_PSEPK | Pp_0574 | DNA-binding response regulator, LuxR family | 0.251661091 | -1.990445912 | 0.002451491 | 2.610569633 | 45 |
| Q88FI6 | Q88FI6_PSEPK | Pp_4109 | Uncharacterized protein | 0.2540052 | -1.977070061 | 0.045797526 | 1.339157978 | 46 |
| Q88DG7 | Q88DG7_PSEPK | Pp_4858 | Uncharacterized protein | 0.258666719 | -1.950833651 | 0.030807746 | 1.511340078 | 47 |
| Q88KZ2 | Q88KZ2_PSEPK | Pp_2147 | Uncharacterized protein | 0.261775288 | -1.933599185 | 0.035677859 | 1.447601219 | 48 |
| Q88RG2 | Q88RG2_PSEPK | Pp_0168 | Surface adhesion protein, putative | 0.267002722 | -1.905073645 | 0.033479617 | 1.475219518 | 49 |
| Q88NV5 | Q88NV5_PSEPK | Pp_1099 | Cold-shock domain family protein | 0.268844321 | -1.895157099 | 1.12E-05 | 4.95098374 | 50 |
| Q88EU1 | Q88EU1_PSEPK | Pp_4362 | Uncharacterized protein | 0.269374094 | -1.892316984 | 0.00895416 | 2.047975126 | 51 |
| Q88EQ1 | Q88EQ1_PSEPK | Bkda2 | 2-oxoisovalerate dehydrogenase, beta subunit | 0.272120125 | -1.877684436 | 0.017381464 | 1.75991364 | 52 |
| Q88EP9 | Q88EP9_PSEPK | Lpdv | Dihydrolipoyl dehydrogenase | 0.273134264 | -1.87231779 | 0.014634298 | 1.834628114 | 53 |

| | | | | | | | | |
|---|---|---|---|---|---|---|---|---|
| Q88D47 | Y4981_PSEPK | Pp_4981 | UPF0312 protein PP_4981 | 0.276899507 | -1.852565609 | 0.003053843 | 2.51515325 | 54 |
| Q88R51 | Q88R51_PSEPK | Pp_0283 | Amino acid ABC transporter, ATP-binding protein | 0.279074386 | -1.84127838 | 0.039519096 | 1.403192997 | 55 |
| Q88E78 | Q88E78_PSEPK | Pp_4588 | Nitroreductase family protein | 0.282036859 | -1.826044379 | 0.000635808 | 3.196673707 | 56 |
| Q88HZ0 | Q88HZ0_PSEPK | Pp_3212 | Rieske 2Fe-2S family protein | 0.286066521 | -1.805577429 | 0.001039326 | 2.983248142 | 57 |
| Q88QN1 | RS19_PSEPK | Rpss | 30S ribosomal protein S19 | 0.29117608 | -1.780036251 | 0.000158513 | 3.799936131 | 58 |
| Q88MV7 | Q88MV7_PSEPK | Ffh | Signal recognition particle protein | 0.293890044 | -1.766651607 | 0.000162273 | 3.789754236 | 59 |
| Q88DE5 | Q88DE5_PSEPK | Pp_4881 | Iron ABC transporter, periplasmic iron-binding protein, putative | 0.297106904 | -1.750945965 | 0.004253516 | 2.37125195 | 60 |
| Q88DL5 | NADD_PSEPK | Nadd | Probable nicotinate-nucleotide adenylyltransferase | 0.298577664 | -1.74382185 | 0.042200711 | 1.374680229 | 61 |
| Q88RA6 | Q88RA6_PSEPK | Pp_0227 | Cysteine ABC transporter, periplasmic cysteine-binding protein, putative | 0.308516466 | -1.696580605 | 0.002062668 | 2.685570774 | 62 |
| Q88FS9 | LOLA_PSEPK | Lola | Outer-membrane lipoprotein carrier protein | 0.30983192 | -1.690442313 | 0.001712224 | 2.766439306 | 63 |
| Q88N13 | Q88N13_PSEPK | Bglx | Periplasmic beta-glucosidase | 0.309997996 | -1.689669206 | 0.041478164 | 1.382180472 | 64 |
| Q88G02 | Q88G02_PSEPK | Pp_3924 | Uncharacterized protein | 0.314816082 | -1.667418853 | 0.044223305 | 1.354348804 | 65 |
| Q88N27 | OADC_PSEPK | Pp_1389 | Oxaloacetate decarboxylase | 0.315431481 | -1.66460144 | 0.001377552 | 2.860892029 | 66 |
| Q88BY6 | Q88BY6_PSEPK | Pp_5401 | Uncharacterized protein | 0.316459007 | -1.659909464 | 0.02156041 | 1.666342982 | 67 |
| Q88R95 | SSUD_PSEPK | Ssud | Alkanesulfonate monooxygenase | 0.32209366 | -1.634447829 | 0.005579055 | 2.253439352 | 68 |
| Q88HE3 | Q88HE3_PSEPK | Gntp | Gluconate transporter | 0.322746161 | -1.631528161 | 0.000249703 | 3.602576152 | 69 |
| Q88EQ0 | Q88EQ0_PSEPK | Bkdb | 2-oxoisovalerate dehydrogenase, lipoamide acyltransferase component | 0.324619757 | -1.623177289 | 0.001381926 | 2.859515239 | 70 |
| Q88N97 | RL13_PSEPK | Rplm | 50S ribosomal protein L13 | 0.330599103 | -1.596845285 | 0.000135347 | 3.868550576 | 71 |
| Q88DJ8 | Q88DJ8_PSEPK | Cobi | Precorrin-2 C20-methyltransferase | 0.332462126 | -1.588738096 | 0.042964424 | 1.36689101 | 72 |
| Q88F18 | Q88F18_PSEPK | Gad | Guanine aminohydrolase | 0.336100904 | -1.573033674 | 0.014589759 | 1.835951882 | 73 |
| Q88L82 | Q88L82_PSEPK | Pp_2053 | Uncharacterized protein | 0.338746572 | -1.561721748 | 0.028470712 | 1.545601667 | 74 |
| Q88LN7 | Q88LN7_PSEPK | Pp_1892 | Uncharacterized protein | 0.33984211 | -1.557063467 | 0.011297881 | 1.947002992 | 75 |
| Q88QM4 | RL24_PSEPK | Rplx | 50S ribosomal protein L24 | 0.340359472 | -1.554868835 | 0.000186204 | 3.730010654 | 76 |
| Q88FB1 | Q88FB1_PSEPK | Lpdg | Dihydrolipoyl dehydrogenase | 0.342862369 | -1.544298527 | 0.000668031 | 3.17520344 | 77 |
| Q88DM5 | LIPA_PSEPK | Lipa | Lipoyl synthase | 0.345510311 | -1.533199331 | 0.001938972 | 2.712428492 | 78 |
| Q88LL9 | RL32_PSEPK | Rpmf | 50S ribosomal protein L32 | 0.347462777 | -1.525069661 | 0.000342451 | 3.465401285 | 79 |
| Q88PG5 | Q88PG5_PSEPK | Pp_0885 | Dipeptide ABC transporter, periplasmic peptide-binding protein | 0.351935104 | -1.506618673 | 0.000439384 | 3.357156223 | 80 |
| Q88NC3 | Y1289_PSEPK | Pp_1289 | UPF0246 protein PP_1289 | 0.355941214 | -1.490289105 | 0.034385382 | 1.46362614 | 81 |

| | | | | | | | | |
|---|---|---|---|---|---|---|---|---|
| Q88FI1 | Q88FI1_PSEPK | Pp_4115 | Uncharacterized protein | 0.357329388 | -1.484673522 | 0.043409364 | 1.362416572 | 82 |
| Q88CF8 | Q88CF8_PSEPK | Rnk | Nucleoside diphosphate kinase regulator | 0.358522873 | -1.479862933 | 0.029304656 | 1.533063372 | 83 |
| Q88NZ7 | Q88NZ7_PSEPK | Pp_1056 | Iron-chelator utilization protein, putative | 0.360302338 | -1.472720082 | 0.003377298 | 2.471430594 | 84 |
| Q88HZ3 | Q88HZ3_PSEPK | Pp_3209 | Chlorohydrolase, putative | 0.36311332 | -1.461508243 | 0.003602211 | 2.443430844 | 85 |
| Q88IR3 | Q88IR3_PSEPK | Pp_2936 | ABC transporter, ATP-binding protein | 0.364698917 | -1.455222178 | 0.006109417 | 2.21400026 | 86 |
| Q88PM6 | Q88PM6_PSEPK | Pp_0824 | Phosphate ABC transporter, periplasmic phosphate-binding protein, putative | 0.365858095 | -1.450643915 | 0.021249332 | 1.672654726 | 87 |
| Q88R52 | Q88R52_PSEPK | Pp_0282 | Amino acid ABC transporter, periplasmic amino acid-binding protein | 0.365948303 | -1.45028824 | 0.004332277 | 2.363283791 | 88 |
| Q88D05 | Q88D05_PSEPK | Pp_5024 | Amino acid ABC transporter, periplasmic amino acid-binding protein | 0.381415765 | -1.39056362 | 8.24E-05 | 4.084285 | 89 |
| Q88QM0 | RL6_PSEPK | Rplf | 50S ribosomal protein L6 | 0.385411938 | -1.375526837 | 5.43E-06 | 5.265503029 | 90 |
| Q88GG2 | Q88GG2_PSEPK | Pp_3762 | Response regulator | 0.386533916 | -1.371333086 | 0.030490876 | 1.5158301 | 91 |
| Q88QN0 | RL22_PSEPK | Rplv | 50S ribosomal protein L22 | 0.386736999 | -1.370575302 | 0.005191818 | 2.284680572 | 92 |
| Q88FB3 | Q88FB3_PSEPK | Sucd | Succinyl-CoA ligase [ADP-forming] subunit alpha | 0.386827867 | -1.370236365 | 6.89E-06 | 5.161774108 | 93 |
| Q88DU7 | GREA_PSEPK | Grea | Transcription elongation factor GreA | 0.387175151 | -1.368941733 | 0.003565773 | 2.447846307 | 94 |
| Q88PR1 | Q88PR1_PSEPK | Nadc | Nicotinate-nucleotide pyrophosphorylase | 0.390061506 | -1.358226463 | 0.009806314 | 2.008494215 | 95 |
| Q88GB8 | Q88GB8_PSEPK | Pp_3806 | Hydrolase, isochorismatase family | 0.392865139 | -1.347893938 | 0.000169766 | 3.770150429 | 96 |
| Q88QL0 | RL17_PSEPK | Rplq | 50S ribosomal protein L17 | 0.396128114 | -1.335961 | 0.000148101 | 3.829442133 | 97 |
| Q88QM5 | RL14_PSEPK | Rpln | 50S ribosomal protein L14 | 0.396616707 | -1.334182645 | 0.000347482 | 3.459067668 | 98 |
| Q88ET9 | Q88ET9_PSEPK | Pp_4364 | Anti-sigma F factor antagonist, putative | 0.397472961 | -1.331071375 | 0.003964447 | 2.401817432 | 99 |
| Q88NJ8 | Q88NJ8_PSEPK | Pp_1209 | Cold-shock domain family protein | 0.399050014 | -1.32535852 | 0.01839863 | 1.735214513 | 100 |
| Q88Q10 | RL21_PSEPK | Rplu | 50S ribosomal protein L21 | 0.400876232 | -1.318771212 | 0.002263042 | 2.645307364 | 101 |
| Q88EH6 | ACSA1_PSEPK | Acsa1 | Acetyl-coenzyme A synthetase 1 | 0.404461879 | -1.30592436 | 0.00023267 | 3.633259794 | 102 |
| Q88K34 | Q88K34_PSEPK | Rbsk | Ribokinase | 0.404762268 | -1.304853286 | 0.041119513 | 1.38595204 | 103 |
| Q88QM3 | RL5_PSEPK | Rple | 50S ribosomal protein L5 | 0.40891015 | -1.290144219 | 6.52E-05 | 4.185716196 | 104 |
| Q88QK8 | Q88QK8_PSEPK | Pp_0482 | Ferroxidase | 0.410161561 | -1.285735803 | 0.006472645 | 2.188918215 | 105 |
| Q88P41 | Q88P41_PSEPK | Gltr-2 | DNA-binding response regulator GltR | 0.413037854 | -1.275654089 | 0.000290941 | 3.536195161 | 106 |
| Q88QP4 | RL1_PSEPK | Rpla | 50S ribosomal protein L1 | 0.422637727 | -1.24250654 | 7.10E-05 | 4.148981563 | 107 |
| P0A157 | RL7_PSEPK | Rpll | 50S ribosomal protein L7/L12 | 0.422858794 | -1.241752113 | 8.47E-05 | 4.071967195 | 108 |
| Q88CA0 | RL33_PSEPK | Rpmg | 50S ribosomal protein L33 | 0.423879793 | -1.238272903 | 0.005756454 | 2.239844976 | 109 |

| | | | | | | | | |
|---|---|---|---|---|---|---|---|---|
| Q88RV0 | Q88RV0_PSEPK | Czrr-1 | DNA-binding response regulator CzrR | 0.430553115 | -1.215736868 | 0.002591672 | 2.586420016 | 110 |
| Q88RK2 | Q88RK2_PSEPK | Pp_0127 | Thiol:disulfide interchange protein | 0.431526761 | -1.212478065 | 0.015300155 | 1.815304178 | 111 |
| Q88E83 | Q88E83_PSEPK | Pp_4583 | Peptidase, putative | 0.434235159 | -1.203451551 | 0.029041428 | 1.536982028 | 112 |
| Q88LE6 | Q88LE6_PSEPK | Pp_1987 | Methlytransferase, UbiE/COQ5 family | 0.43557268 | -1.199014626 | 0.00193554 | 2.713197958 | 113 |
| Q88C99 | RL28_PSEPK | Rpmb | 50S ribosomal protein L28 | 0.436420702 | -1.196208555 | 0.008630188 | 2.063979735 | 114 |
| Q88QN5 | RL3_PSEPK | Rplc | 50S ribosomal protein L3 | 0.437355711 | -1.193120961 | 0.000128346 | 3.891616565 | 115 |
| Q88KR7 | Q88KR7_PSEPK | Pp_2223 | Uncharacterized protein | 0.438341509 | -1.189872794 | 0.03227605 | 1.491119621 | 116 |
| Q88MV3 | RL19_PSEPK | Rpls | 50S ribosomal protein L19 | 0.440916841 | -1.181421514 | 0.000263074 | 3.579921699 | 117 |
| Q88QP5 | RL11_PSEPK | Rplk | 50S ribosomal protein L11 | 0.44310538 | -1.174278252 | 0.000286873 | 3.542310274 | 118 |
| Q88DI4 | Q88DI4_PSEPK | Pp_4841 | Branched-chain amino acid ABC transporter, periplasmic amino acid-binding protein, putative | 0.445269451 | -1.167249461 | 0.002780529 | 2.555872585 | 119 |
| Q88C70 | Q88C70_PSEPK | Hupa | DNA-binding protein HU-alpha | 0.446695866 | -1.16263519 | 0.005169892 | 2.286518565 | 120 |
| Q88PX7 | RL25_PSEPK | Rply | 50S ribosomal protein L25 | 0.448450683 | -1.156978756 | 0.000289175 | 3.538838738 | 121 |
| Q88QL7 | RL30_PSEPK | Rpmd | 50S ribosomal protein L30 | 0.448535556 | -1.156705741 | 0.001381602 | 2.859617155 | 122 |
| P0A136 | KAD_PSEPK | Adk | Adenylate kinase | 0.456394692 | -1.131646081 | 0.000789687 | 3.10254508 | 123 |
| Q88QN4 | RL4_PSEPK | Rpld | 50S ribosomal protein L4 | 0.456935106 | -1.129938808 | 0.000115621 | 3.936963105 | 124 |
| Q88QM8 | RL16_PSEPK | Rplp | 50S ribosomal protein L16 | 0.460263969 | -1.119466586 | 1.69E-05 | 4.772537901 | 125 |
| Q88Q09 | RL27_PSEPK | Rpma | 50S ribosomal protein L27 | 0.460382155 | -1.119096182 | 0.011103252 | 1.954549822 | 126 |
| Q88DA5 | THIC_PSEPK | Thic | Phosphomethylpyrimidine synthase | 0.46218385 | -1.113461247 | 0.001833906 | 2.736622978 | 127 |
| Q88N89 | Q88N89_PSEPK | Gmha | Phosphoheptose isomerase | 0.463566551 | -1.109151624 | 0.035993584 | 1.443774904 | 128 |
| Q88QL9 | RL18_PSEPK | Rplr | 50S ribosomal protein L18 | 0.464085674 | -1.107536932 | 3.42E-05 | 4.465620343 | 129 |
| Q88QN3 | RL23_PSEPK | Rplw | 50S ribosomal protein L23 | 0.465431006 | -1.103360773 | 0.000162789 | 3.788374798 | 130 |
| Q88FG5 | Q88FG5_PSEPK | Nuon | NADH-quinone oxidoreductase subunit N | 0.4657677 | -1.102317501 | 0.000983462 | 3.007242387 | 131 |
| Q88QN2 | RL2_PSEPK | Rplb | 50S ribosomal protein L2 | 0.467169772 | -1.097981165 | 2.18E-06 | 5.661485242 | 132 |
| Q88QM7 | RL29_PSEPK | Rpmc | 50S ribosomal protein L29 | 0.468348727 | -1.09434495 | 0.000549265 | 3.260218446 | 133 |
| Q88LV1 | Q88LV1_PSEPK | Pp_1826 | Hydrolase, isochorismatase family | 0.472993622 | -1.080107366 | 0.013800522 | 1.860104474 | 134 |
| Q88NR3 | Q88NR3_PSEPK | Pp_1142 | Uncharacterized protein | 0.474869458 | -1.074397123 | 0.012551037 | 1.901320397 | 135 |
| Q88NR8 | Q88NR8_PSEPK | Brag | Branched-chain amino acid ABC transporter, ATP-binding protein | 0.475250812 | -1.073239002 | 0.022743585 | 1.643141076 | 136 |
| Q88CB2 | Q88CB2_PSEPK | Dadx | Alanine racemase | 0.47665299 | -1.068988749 | 0.020531589 | 1.687577439 | 137 |

| | | | | | | | | |
|---|---|---|---|---|---|---|---|---|
| Q88NB6 | Q88NB6_PSEPK | Estb | Carboxylesterase | 0.480967737 | -1.055987973 | 0.047600307 | 1.322390243 | 138 |
| Q88KH9 | Q88KH9_PSEPK | Pp_2311 | Hydrolase, TatD family | 0.486802607 | -1.038591202 | 0.022043935 | 1.656710878 | 139 |
| Q88NR7 | Q88NR7_PSEPK | Braf | Branched-chain amino acid ABC transporter, ATP-binding protein | 0.487624504 | -1.036157468 | 0.00174108 | 2.759181341 | 140 |
| Q88FH1 | NUOH_PSEPK | Nuoh | NADH-quinone oxidoreductase subunit H | 0.487674003 | -1.036011029 | 0.028273428 | 1.54862153 | 141 |
| Q88QL6 | RL15_PSEPK | Rplo | 50S ribosomal protein L15 | 0.488334263 | -1.034059089 | 1.96E-05 | 4.707100958 | 142 |
| Q88BZ4 | Q88BZ4_PSEPK | Pp_5389 | Uncharacterized protein | 0.492151629 | -1.022825225 | 0.010176251 | 1.992412176 | 143 |
| Q88K24 | RL20_PSEPK | Rplt | 50S ribosomal protein L20 | 0.498465113 | -1.004435559 | 0.002665317 | 2.574251121 | 144 |
| Q88DF1 | RL9_PSEPK | Rpli | 50S ribosomal protein L9 | 0.498471787 | -1.004416243 | 0.000113018 | 3.946853441 | 145 |

Supplementary Table S4. List of upregulated proteins from PP_4188 strain

| Protein Group | Protein Names | Protein | Protein Description | Fold Change | Log2(Fold Change) | P-Value (Equal Variance) | (-Log10(P-Value)) | Rank |
|---|---|---|---|---|---|---|---|---|
| Q88JG3 | Q88JG3_PSEPK | Pp_2686 | Transglutaminase-like superfamily domain protein | 459.0034538 | 8.842361199 | 0.003048855 | 2.515863161 | 1 |
| Q88RJ9 | Q88RJ9_PSEPK | Pp_0130 | N-acetylmuramoyl-L-alanine amidase family protein | 88.20536801 | 6.462794553 | 0.003064274 | 2.513672352 | 2 |
| Q88CB4 | Q88CB4_PSEPK | Pp_5267 | Cytochrome c5 | 60.96447367 | 5.929896869 | 0.001960893 | 2.707546189 | 3 |
| Q88PY7 | Q88PY7_PSEPK | Pp_0711 | Hydrolase, isochorismatase family | 57.66882279 | 5.849719667 | 0.000501779 | 3.299487589 | 4 |
| Q88E52 | Q88E52_PSEPK | Pp_4616 | Uncharacterized protein | 38.90917988 | 5.282038667 | 0.017043091 | 1.768451639 | 5 |
| Q88IQ5 | Q88IQ5_PSEPK | Pp_2944 | Histidine kinase | 30.49895482 | 4.930687898 | 0.005404422 | 2.267250714 | 6 |
| Q88LN0 | Q88LN0_PSEPK | Pp_1899 | ExbD/TolR family protein | 30.31085388 | 4.92176259 | 0.012475951 | 1.903926356 | 7 |
| Q88HP1 | Q88HP1_PSEPK | Pp_3312 | Heat shock protein, putative | 25.19738 | 4.655201826 | 0.003227934 | 2.491075352 | 8 |
| Q88E49 | Q88E49_PSEPK | Pp_4619 | Maleylacetoacetate isomerase, putative | 19.56087481 | 4.289898988 | 0.011635203 | 1.934226047 | 9 |
| Q88D45 | Q88D45_PSEPK | Pp_4983 | Flavin-containing monamine oxidase family protein | 17.35932008 | 4.117638537 | 0.024520733 | 1.610466556 | 10 |
| Q88EE4 | Q88EE4_PSEPK | Pp_4521 | Aerotaxis receptor, putative | 16.9483255 | 4.083070836 | 0.006182897 | 2.208807957 | 11 |
| Q88DP4 | Q88DP4_PSEPK | Pp_4780 | Acyl-CoA dehydrogenase, putative | 15.61959621 | 3.965285253 | 0.003061331 | 2.514089711 | 12 |
| Q88NM0 | Q88NM0_PSEPK | Phoq | Histidine kinase | 14.5459937 | 3.862549952 | 0.00884763 | 2.053173054 | 13 |
| Q88M34 | Q88M34_PSEPK | Pp_1741 | Glycine betaine-binding protein, putative | 13.85113863 | 3.791932672 | 0.030945149 | 1.509407425 | 14 |
| Q88DJ5 | Q88DJ5_PSEPK | Cobl | Precorrin-6Y C5,15-methyltransferase (Decarboxylating) | 13.84523157 | 3.791317279 | 0.035451489 | 1.450365525 | 15 |
| Q88PB6 | Q88PB6_PSEPK | Mrec | Rod shape determining protein MreC | 13.60784043 | 3.766366223 | 0.013448826 | 1.871315628 | 16 |
| Q88JJ7 | Q88JJ7_PSEPK | Pp_2652 | Hydratase/decarboxylase, putative | 13.41915267 | 3.746221673 | 0.000799959 | 3.096932481 | 17 |
| Q88N53 | Q88N53_PSEPK | Pp_1363 | Uncharacterized protein | 12.92113841 | 3.691661278 | 0.022746793 | 1.643079828 | 18 |
| Q88DC8 | Q88DC8_PSEPK | Nnrd | ADP-dependent (S)-NAD(P)H-hydrate dehydratase | 12.76244475 | 3.673832811 | 0.032295579 | 1.490856927 | 19 |
| Q88C54 | Q88C54_PSEPK | Psts | Phosphate ABC transporter, periplasmic phosphate-binding protein | 12.14710001 | 3.602540022 | 0.000845345 | 3.072965984 | 20 |
| Q88DK4 | Q88DK4_PSEPK | Fis | DNA-binding protein Fis | 11.23072273 | 3.489378867 | 6.84E-05 | 4.165024197 | 21 |
| Q88KS3 | Q88KS3_PSEPK | Pp_2216 | Acyl-CoA dehydrogenase family protein | 10.24297441 | 3.356562809 | 0.005393366 | 2.268140139 | 22 |
| Q88QU5 | PLSY_PSEPK | Plsy | Glycerol-3-phosphate acyltransferase | 9.678320055 | 3.274756649 | 0.001698101 | 2.770036483 | 23 |
| Q88LC8 | Q88LC8_PSEPK | Pp_2007 | P-47-related protein | 9.667907059 | 3.273203604 | 0.010635955 | 1.973223496 | 24 |
| Q88M09 | MTNA_PSEPK | Mtna | Methylthioribose-1-phosphate isomerase | 9.414949756 | 3.234953396 | 0.002378285 | 2.623736177 | 25 |

| | | | | | | | | |
|---|---|---|---|---|---|---|---|---|
| Q88CC9 | Q88CC9_PSEPK | Pp_5252 | Uncharacterized protein | 8.723315723 | 3.124876606 | 0.009403538 | 2.026708721 | 26 |
| Q88MZ0 | Q88MZ0_PSEPK | Muca | Sigma factor algU negative regulatory protein MucA | 8.386117565 | 3.068003056 | 0.00354038 | 2.450950113 | 27 |
| Q88GT8 | Q88GT8_PSEPK | Pp_3629 | Uncharacterized protein | 8.211594286 | 3.037662349 | 0.007758249 | 2.110236277 | 28 |
| Q88RC9 | Q88RC9_PSEPK | Pp_0203 | Tabtoxinine-beta-lactam limiting dipeptidase, putative | 8.010420484 | 3.001877975 | 0.047758654 | 1.320947926 | 29 |
| Q88RN5 | Q88RN5_PSEPK | Pp_0094 | Hydrolase, haloacid dehalogenase-like family | 7.867687746 | 2.975939701 | 0.006940983 | 2.158579011 | 30 |
| Q88IV6 | Q88IV6_PSEPK | Pp_2893 | Uncharacterized protein | 7.452097915 | 2.89764663 | 0.00134017 | 2.872840005 | 31 |
| Q88LW9 | G6PI1_PSEPK | Pgi1 | Glucose-6-phosphate isomerase 1 | 7.312441416 | 2.870353161 | 0.023205529 | 1.634408534 | 32 |
| Q88NL6 | Q88NL6_PSEPK | Pp_1191 | S4 domain protein | 7.275087275 | 2.862964555 | 0.008887511 | 2.051219829 | 33 |
| Q88MQ2 | Q88MQ2_PSEPK | Pp_1518 | Uncharacterized protein | 7.162197178 | 2.840402237 | 0.006770031 | 2.169409355 | 34 |
| Q88DZ6 | Q88DZ6_PSEPK | Recd | RecBCD enzyme subunit RecD | 7.04178201 | 2.815940567 | 0.003876996 | 2.411504612 | 35 |
| Q88H61 | Q88H61_PSEPK | Pp_3504 | Uncharacterized protein | 6.988707546 | 2.805025676 | 0.007010937 | 2.15422392 | 36 |
| Q88CW0 | Q88CW0_PSEPK | Pp_5070 | ParA family protein | 6.933905454 | 2.793668165 | 0.004037365 | 2.39390204 | 37 |
| Q88F81 | Q88F81_PSEPK | Fpva | Outer membrane ferripyoverdine receptor | 6.296461942 | 2.654541389 | 0.043095727 | 1.365565784 | 38 |
| Q88M01 | Q88M01_PSEPK | Pp_1775 | Metallo-beta-lactamase family protein | 5.938918999 | 2.570200356 | 0.009203227 | 2.036059882 | 39 |
| Q88KC9 | Q88KC9_PSEPK | Csuc | Type 1 pili usher pathway chaperone CsuC | 5.881775474 | 2.556251713 | 0.011863986 | 1.925769358 | 40 |
| Q88MB9 | RLMD_PSEPK | Rlmd | 23S rRNA (uracil(1939)-C(5))-methyltransferase RlmD | 5.801422347 | 2.536406652 | 0.002810696 | 2.55118619 | 41 |
| Q88E48 | Q88E48_PSEPK | Pp_4620 | Fumarylacetoacetase | 5.778812954 | 2.530773174 | 0.000315647 | 3.500798538 | 42 |
| Q88CL6 | Q88CL6_PSEPK | Pp_5164 | Penicillin amidase family protein | 5.69835622 | 2.510545811 | 0.009369588 | 2.028279524 | 43 |
| Q88LC9 | Q88LC9_PSEPK | Pp_2006 | Uncharacterized protein | 5.510757355 | 2.462250605 | 0.000130075 | 3.885805077 | 44 |
| Q88HC7 | Q88HC7_PSEPK | Hpd | 4-hydroxyphenylpyruvate dioxygenase | 5.399937546 | 2.432942722 | 4.21E-05 | 4.37534556 | 45 |
| Q88L41 | Q88L41_PSEPK | Pp_2094 | Nitrate-binding protein NasS, putative | 5.334430754 | 2.415334327 | 0.000101229 | 3.994696689 | 46 |
| Q88IC8 | Q88IC8_PSEPK | Pp_3071 | Acetoacetyl-CoA synthetase, putative | 5.267841461 | 2.397211927 | 0.02260243 | 1.64584487 | 47 |
| Q88QC4 | Q88QC4_PSEPK | Pp_0570 | Extracellular protein, putative | 5.021260677 | 2.328049624 | 0.031803616 | 1.497523504 | 48 |
| Q88I56 | Q88I56_PSEPK | Pp_3145 | Uncharacterized protein | 5.013769438 | 2.325895654 | 0.000500445 | 3.300643488 | 49 |
| Q88M19 | Q88M19_PSEPK | Pp_1756 | Uncharacterized protein | 4.913362942 | 2.296710813 | 0.0003559 | 3.448672588 | 50 |
| Q88CK5 | Q88CK5_PSEPK | Pp_5175 | HlyD family secretion protein | 4.82346731 | 2.270070589 | 0.002030619 | 2.692371631 | 51 |
| Q88DC1 | Q88DC1_PSEPK | Pp_4906 | Uncharacterized protein | 4.801721719 | 2.263551795 | 0.002406801 | 2.618559857 | 52 |
| Q88RI5 | Q88RI5_PSEPK | Pp_0144 | Metalloprotease, insulinase family | 4.777452268 | 2.256241459 | 0.005184855 | 2.285263379 | 53 |

| | | | | | | | | |
|---|---|---|---|---|---|---|---|---|
| Q88K21 | Q88K21_PSEPK | Pp_2472 | Uncharacterized protein | 4.763535921 | 2.252032868 | 0.008221601 | 2.08504358 | 54 |
| Q88EC4 | Q88EC4_PSEPK | Pp_4542 | ABC transporter, ATP-binding protein/permease protein, putative | 4.693388688 | 2.230629944 | 0.040386512 | 1.393763656 | 55 |
| Q88JY8 | Q88JY8_PSEPK | Pp_2505 | GAF domain/GGDEF domain protein | 4.380823012 | 2.13120193 | 0.036713572 | 1.435173361 | 56 |
| Q88CE9 | Q88CE9_PSEPK | Pp_5232 | Uncharacterized protein | 4.321153139 | 2.11141636 | 0.000196802 | 3.705969906 | 57 |
| Q88P27 | Q88P27_PSEPK | Pp_1026 | Peptidase, M23/M37 family | 4.275299562 | 2.09602551 | 0.001991626 | 2.700792225 | 58 |
| Q88MQ0 | PLSB_PSEPK | Plsb | Glycerol-3-phosphate acyltransferase | 4.263055247 | 2.091887752 | 0.017684107 | 1.752416869 | 59 |
| Q88L37 | Q88L37_PSEPK | Dacb | D-alanyl-D-alanine carboxypeptidase | 4.259800692 | 2.090785931 | 0.009640887 | 2.015883029 | 60 |
| Q88DB9 | PSD_PSEPK | Psd | Phosphatidylserine decarboxylase proenzyme | 4.25246664 | 2.088299918 | 0.003100219 | 2.508607643 | 61 |
| Q88EY6 | Q88EY6_PSEPK | Pp_4314 | Uncharacterized protein | 4.227055279 | 2.079652979 | 0.01212581 | 1.916289255 | 62 |
| Q88ME0 | Q88ME0_PSEPK | Pp_1633 | Uncharacterized protein | 4.190802525 | 2.067226542 | 0.00593972 | 2.226234028 | 63 |
| P0A167 | RNPA_PSEPK | Rnpa | Ribonuclease P protein component | 4.18806912 | 2.066285253 | 0.027970865 | 1.553294102 | 64 |
| Q88CW3 | Q88CW3_PSEPK | Pp_5067 | Potassium efflux system protein KefA, putative | 4.076253575 | 2.027243801 | 0.005120972 | 2.290647592 | 65 |
| Q88PD1 | Q88PD1_PSEPK | Pp_0919 | Uncharacterized protein | 4.054301724 | 2.019453459 | 0.000409479 | 3.387768167 | 66 |
| Q88JX1 | Q88JX1_PSEPK | Pp_2526 | Uncharacterized protein | 4.022321737 | 2.008028484 | 0.022203353 | 1.653581434 | 67 |
| Q88DC4 | RSGA_PSEPK | Rsga | Putative ribosome biogenesis GTPase RsgA | 3.99270515 | 1.997366537 | 0.00131489 | 2.881110589 | 68 |
| Q88KW4 | Q88KW4_PSEPK | Pp_2175 | 3-oxoacyl-(Acyl-carrier-protein) reductase, putative | 3.987282998 | 1.995406004 | 0.001205249 | 2.918923325 | 69 |
| Q877U6 | Q877U6_PSEPK | Pp_1157 | Acetolactate synthase, catabolic, putative | 3.923114365 | 1.971999393 | 2.68E-05 | 4.571579264 | 70 |
| Q88KB0 | Q88KB0_PSEPK | Pp_2380 | Uncharacterized protein | 3.902701442 | 1.9644731 | 0.010408533 | 1.982610463 | 71 |
| O85207 | PHAG_PSEPK | Phag | (R)-3-hydroxydecanoyl-ACP:CoA transacylase | 3.89953905 | 1.963303598 | 0.003903401 | 2.408556839 | 72 |
| Q88CU6 | Q88CU6_PSEPK | Pp_5084 | Penicillin-binding protein | 3.882236479 | 1.956888 | 0.00093553 | 3.028942143 | 73 |
| P61113 | RL36_PSEPK | Rpmj | 50S ribosomal protein L36 | 3.790401957 | 1.922350848 | 0.0108888 | 1.963019959 | 74 |
| Q88D40 | Q88D40_PSEPK | Pp_4988 | Type IV pili sensor histidine kinase/response regulator, PilL/ChpA fusion protein | 3.772176898 | 1.915397333 | 0.016598082 | 1.779942099 | 75 |
| Q88NI7 | Q88NI7_PSEPK | Tolr | Biopolymer transport protein TolR | 3.756971785 | 1.909570281 | 0.015464358 | 1.810668118 | 76 |
| P0A130 | Y1224_PSEPK | Pp_1224 | Uncharacterized protein PP_1224 | 3.750815971 | 1.907204481 | 0.001569159 | 2.804333122 | 77 |
| Q88FW0 | Q88FW0_PSEPK | Pp_3968 | Histidine kinase | 3.715910227 | 1.893715648 | 0.036439428 | 1.438428446 | 78 |
| Q88RS1 | Q88RS1_PSEPK | Pp_0058 | 1-acyl-sn-glycerol-3-phosphate acyltransferase, putative | 3.689170044 | 1.883296288 | 0.008158845 | 2.088371328 | 79 |
| Q88MS6 | Q88MS6_PSEPK | Pp_1492 | Sensor histidine kinase/response regulator | 3.674718929 | 1.877633906 | 0.013384523 | 1.873397107 | 80 |

| | | | | | | | | |
|---|---|---|---|---|---|---|---|---|
| Q88N16 | Q88N16_PSEPK | Pp_1400 | Dicarboxylate MFS transporter | 3.666846913 | 1.874540036 | 0.010165201 | 1.992884018 | 81 |
| Q88DK6 | Q88DK6_PSEPK | Pp_4819 | Uncharacterized protein | 3.62382242 | 1.85751226 | 0.00212909 | 2.671806003 | 82 |
| Q88KT1 | PHNX_PSEPK | Phnx | Phosphonoacetaldehyde hydrolase | 3.604049982 | 1.849619019 | 0.000577762 | 3.238250797 | 83 |
| Q88HW7 | Q88HW7_PSEPK | Pp_3236 | Lipoprotein OprI, putative | 3.576865933 | 1.838696045 | 0.009884223 | 2.00505748 | 84 |
| Q88MC3 | Q88MC3_PSEPK | Gacs | Histidine kinase | 3.511881211 | 1.812244047 | 0.015934144 | 1.797671275 | 85 |
| Q88PV6 | Q88PV6_PSEPK | Pp_0742 | Uncharacterized protein | 3.459780322 | 1.790680437 | 0.003862459 | 2.41313609 | 86 |
| Q88C51 | Q88C51_PSEPK | Pp_5332 | Uncharacterized protein | 3.437296672 | 1.781274376 | 0.000108992 | 3.962606519 | 87 |
| Q88DI0 | Q88DI0_PSEPK | Pp_4845 | Branched-chain amino acid ABC transporter, ATP-binding protein, putative | 3.434541283 | 1.780117426 | 0.000158732 | 3.799334345 | 88 |
| Q88DW4 | Q88DW4_PSEPK | Pp_4704 | Uncharacterized protein | 3.35528551 | 1.746435534 | 7.20E-05 | 4.142933089 | 89 |
| Q88E51 | Q88E51_PSEPK | Pp_4617 | Leucine dehydrogenase | 3.332093816 | 1.736429021 | 0.006733916 | 2.171732274 | 90 |
| Q88EF7 | Q88EF7_PSEPK | Pp_4508 | Transcriptional regulator, AraC family | 3.32813709 | 1.734714861 | 0.026797174 | 1.571911004 | 91 |
| Q88HQ9 | Q88HQ9_PSEPK | Pp_3294 | Universal stress protein family | 3.294468224 | 1.720045612 | 0.022102358 | 1.655561387 | 92 |
| Q88NS3 | Q88NS3_PSEPK | Pp_1131 | Outer membrane lipoprotein, putative | 3.244875466 | 1.698163111 | 0.023669033 | 1.62581948 | 93 |
| Q88KA9 | Q88KA9_PSEPK | Pp_2381 | Conserved domain protein | 3.229224919 | 1.691187929 | 0.009371916 | 2.028171634 | 94 |
| Q88KQ8 | Q88KQ8_PSEPK | Pp_2232 | Transcriptional regulator, Cro/CI family | 3.226853638 | 1.690128142 | 0.015772765 | 1.802092176 | 95 |
| Q88PS1 | Q88PS1_PSEPK | Pp_0777 | Glutathione peroxidase | 3.214048995 | 1.684391921 | 0.036109513 | 1.442378371 | 96 |
| Q88EB2 | Q88EB2_PSEPK | Pp_4554 | Uncharacterized protein | 3.188541716 | 1.672896756 | 0.018055167 | 1.743398486 | 97 |
| Q88NF9 | MQO2_PSEPK | Mqo2 | Probable malate:quinone oxidoreductase 2 | 3.187756233 | 1.672541311 | 0.001128764 | 2.947396997 | 98 |
| Q88NM1 | Q88NM1_PSEPK | Phop | Transcriptional regulatory protein PhoP | 3.163337214 | 1.661447356 | 0.025301905 | 1.596846782 | 99 |
| Q88RN8 | Q88RN8_PSEPK | Pp_0091 | Lipoprotein, putative | 3.160825598 | 1.660301435 | 0.002992783 | 2.523924833 | 100 |
| Q88M79 | Q88M79_PSEPK | Pp_1695 | Sodium-solute symporter/sensory box histidine kinase/response regulator, putative | 3.149106846 | 1.654942707 | 0.00826405 | 2.082807047 | 101 |
| Q88FI4 | EFG2_PSEPK | Fusb | Elongation factor G 2 | 3.124693353 | 1.643714615 | 0.006877081 | 2.162595866 | 102 |
| Q88DW6 | ACSA2_PSEPK | Acsa2 | Acetyl-coenzyme A synthetase 2 | 3.107748959 | 1.635869969 | 0.011561097 | 1.937000941 | 103 |
| Q88KN3 | Q88KN3_PSEPK | Aer-1 | Aerotaxis receptor Aer-1 | 3.069718041 | 1.618106148 | 0.007263697 | 2.138842304 | 104 |
| Q88Q44 | MDH_PSEPK | Mdh | Probable malate dehydrogenase | 3.069691819 | 1.618093824 | 0.002794901 | 2.553633628 | 105 |
| Q88D80 | Q88D80_PSEPK | Puta | Bifunctional protein PutA | 3.067986063 | 1.617291929 | 0.001177184 | 2.929155672 | 106 |
| Q88K69 | Q88K69_PSEPK | Pp_2422 | Alkyl hydroperoxide reductase AhpD | 3.065244386 | 1.616002102 | 0.043252042 | 1.36399338 | 107 |

| | | | | | | | | |
|---|---|---|---|---|---|---|---|---|
| Q88EX1 | Q88EX1_PSEPK | Pp_4331 | Uncharacterized protein | 3.057273276 | 1.612245514 | 0.001648095 | 2.783017779 | 108 |
| Q88LF0 | Q88LF0_PSEPK | Pp_1983 | Sensory box protein | 3.018414545 | 1.593790957 | 0.00881262 | 2.054894974 | 109 |
| Q88KS4 | Q88KS4_PSEPK | Fadax | 3-ketoacyl-CoA thiolase | 3.018072185 | 1.593627312 | 0.019248151 | 1.715610983 | 110 |
| Q88RB8 | Q88RB8_PSEPK | Pp_0215 | Response regulator | 3.013963327 | 1.591661863 | 0.00069769 | 3.156337233 | 111 |
| Q88C90 | Q88C90_PSEPK | Pp_5293 | Uncharacterized protein | 2.985793253 | 1.578114272 | 0.00011201 | 3.950742572 | 112 |
| Q88QY0 | Q88QY0_PSEPK | Pp_0354 | CBS domain protein | 2.9361623 | 1.553931717 | 0.004594504 | 2.337761324 | 113 |
| Q88I67 | Q88I67_PSEPK | Pp_3134 | Conserved domain protein | 2.924345977 | 1.548114005 | 0.025319802 | 1.596539701 | 114 |
| Q88LM1 | Y1909_PSEPK | Maf-2 | Maf-like protein PP_1909 | 2.906335195 | 1.539201102 | 0.008306641 | 2.080574565 | 115 |
| Q88DQ3 | Q88DQ3_PSEPK | Pp_4770 | Uncharacterized protein | 2.900874863 | 1.536488062 | 0.022417803 | 1.64940695 | 116 |
| Q88HB9 | Q88HB9_PSEPK | Pp_3441 | Uncharacterized protein | 2.881795383 | 1.526967903 | 0.008367512 | 2.077403672 | 117 |
| Q88P96 | Q88P96_PSEPK | Pp_0956 | Phosphatase, YrbI family | 2.818286384 | 1.49481822 | 0.000186359 | 3.729648984 | 118 |
| Q88IA6 | Q88IA6_PSEPK | Pp_3093 | Uncharacterized protein | 2.81567727 | 1.493481983 | 0.001802545 | 2.74411384 | 119 |
| Q88CK6 | Q88CK6_PSEPK | Pp_5174 | Efflux membrane fusion protein, RND family | 2.811924307 | 1.49155776 | 0.044618399 | 1.350486018 | 120 |
| Q88CI9 | GCSP2_PSEPK | Gcvp2 | Glycine dehydrogenase (decarboxylating) 2 | 2.790557056 | 1.480553144 | 0.033753361 | 1.471682974 | 121 |
| Q88R38 | Q88R38_PSEPK | Pp_0296 | Glycine betaine/L-proline ABC transporter, periplasmic binding protein | 2.771795529 | 1.470820836 | 0.006992575 | 2.15536284 | 122 |
| Q88PU9 | Q88PU9_PSEPK | Pp_0749 | Uncharacterized protein | 2.766741007 | 1.4681876 | 0.030765053 | 1.511942329 | 123 |
| Q88QB4 | Q88QB4_PSEPK | Pp_0580 | MaoC domain protein | 2.73288934 | 1.450427044 | 0.043143403 | 1.365085601 | 124 |
| Q88KT0 | PHNW_PSEPK | Phnw | 2-aminoethylphosphonate--pyruvate transaminase | 2.702469451 | 1.43427831 | 0.001309448 | 2.882911821 | 125 |
| Q88HR2 | Q88HR2_PSEPK | Pp_3291 | Metallo-beta-lactamase family protein | 2.689278595 | 1.427219219 | 4.25E-05 | 4.372019498 | 126 |
| Q88HH4 | Q88HH4_PSEPK | Pp_3384 | Uncharacterized protein | 2.683043182 | 1.423870274 | 0.000121289 | 3.916176809 | 127 |
| Q88P97 | Q88P97_PSEPK | Lptc | Lipopolysaccharide export system protein LptC | 2.658021731 | 1.4103529 | 0.008449642 | 2.073161696 | 128 |
| Q88MR5 | Q88MR5_PSEPK | Pp_1504 | Uncharacterized protein | 2.657632538 | 1.410141642 | 0.001403926 | 2.852655714 | 129 |
| Q88KY4 | LOLD_PSEPK | Lold | Lipoprotein-releasing system ATP-binding protein LolD | 2.642728111 | 1.402028005 | 0.010006369 | 1.999723471 | 130 |
| Q88P86 | HIS8_PSEPK | Hisc | Histidinol-phosphate aminotransferase | 2.630237936 | 1.395193314 | 0.000107007 | 3.970586809 | 131 |
| Q88KS5 | Q88KS5_PSEPK | Fadb2x | 3-hydroxyacyl-CoA dehydrogenase FadB2x | 2.611083838 | 1.384648781 | 0.008078966 | 2.09264424 | 132 |
| Q88JS9 | Q88JS9_PSEPK | Pp_2568 | Ring-cleaving dioxygenase | 2.569676449 | 1.361586719 | 0.02532022 | 1.596532526 | 133 |
| Q88PS5 | Q88PS5_PSEPK | Pp_0773 | OmpA family protein | 2.56967228 | 1.361584379 | 0.010660016 | 1.972242139 | 134 |
| Q88EJ2 | Q88EJ2_PSEPK | Algz | Alginate biosynthesis transcriptional activator | 2.567579498 | 1.360408946 | 0.02921382 | 1.534411658 | 135 |

| | | | | | | | | |
|---|---|---|---|---|---|---|---|---|
| Q88PG3 | Q88PG3_PSEPK | Pp_0887 | Histidine kinase | 2.556124391 | 1.353958045 | 0.034414314 | 1.463260888 | 136 |
| Q88PQ1 | Q88PQ1_PSEPK | Pp_0797 | Uncharacterized protein | 2.552680551 | 1.352013006 | 0.00846459 | 2.072394093 | 137 |
| Q88EZ4 | Q88EZ4_PSEPK | Pp_4306 | Uncharacterized protein | 2.547939014 | 1.349330747 | 0.013592431 | 1.866702852 | 138 |
| Q88KY5 | Q88KY5_PSEPK | Lolc | Lipoprotein releasing system, permease protein | 2.544178813 | 1.347200071 | 0.015936052 | 1.797619253 | 139 |
| Q88MI2 | GLND_PSEPK | Glnd | Bifunctional uridylyltransferase/uridylyl-removing enzyme | 2.510491064 | 1.32796959 | 0.006033056 | 2.219462659 | 140 |
| Q88CC3 | Q88CC3_PSEPK | Pp_5258 | Aldehyde dehydrogenase family protein | 2.508718416 | 1.326950549 | 0.017464649 | 1.757840141 | 141 |
| Q88CE7 | Q88CE7_PSEPK | Pp_5234 | Nitrogen regulatory protein P-II | 2.505890303 | 1.325323261 | 0.000175077 | 3.756770139 | 142 |
| Q88D37 | Q88D37_PSEPK | Pilh | Type IV pili response regulator PilH | 2.496551613 | 1.319936733 | 0.039224525 | 1.406442304 | 143 |
| Q88QD0 | Q88QD0_PSEPK | Pp_0564 | Uncharacterized protein | 2.490462906 | 1.316413923 | 1.47E-05 | 4.833826113 | 144 |
| Q88EA3 | Q88EA3_PSEPK | Pp_4563 | Uncharacterized protein | 2.468693895 | 1.303747961 | 0.002925113 | 2.533857297 | 145 |
| Q88PF7 | Q88PF7_PSEPK | Pp_0893 | ThiJ/PfpI family protein | 2.456225015 | 1.296442732 | 0.003038115 | 2.517395852 | 146 |
| Q88DF5 | Q88DF5_PSEPK | Pp_4870 | Azurin | 2.441877278 | 1.287990696 | 0.018069656 | 1.743050109 | 147 |
| Q88E25 | Q88E25_PSEPK | Pp_4643 | Xanthine/uracil permease family protein | 2.441875058 | 1.287989385 | 0.04779814 | 1.320589006 | 148 |
| Q88LM8 | Y1901_PSEPK | Pp_1901 | UPF0434 protein PP_1901 | 2.390732948 | 1.257452986 | 0.04878949 | 1.311673726 | 149 |
| Q88I66 | Q88I66_PSEPK | Pp_3135 | Glycosyl transferase, putative | 2.369485452 | 1.244573803 | 0.004745717 | 2.32369817 | 150 |
| Q88BY7 | Q88BY7_PSEPK | Pp_5400 | Uncharacterized protein | 2.341196343 | 1.22724593 | 0.013975207 | 1.854641743 | 151 |
| Q88GG3 | Q88GG3_PSEPK | Pp_3761 | Sensor histidine kinase/response regulator | 2.328111519 | 1.219160167 | 0.000121563 | 3.915197219 | 152 |
| Q88KP1 | Q88KP1_PSEPK | Pp_2249 | Methyl-accepting chemotaxis transducer | 2.308074387 | 1.206689721 | 0.009791211 | 2.009163598 | 153 |
| Q88QB3 | Q88QB3_PSEPK | Pp_0581 | 3-oxoacyl-(Acyl-carrier-protein) reductase | 2.281581696 | 1.190034313 | 0.042072853 | 1.375998039 | 154 |
| Q88HH5 | Q88HH5_PSEPK | Pp_3383 | Gluconate dehydrogenase, putative | 2.269208032 | 1.182188876 | 0.001705768 | 2.768080072 | 155 |
| Q88R35 | Q88R35_PSEPK | Pp_0299 | Uncharacterized protein | 2.260510932 | 1.176648894 | 0.004557551 | 2.341268501 | 156 |
| Q88N18 | Q88N18_PSEPK | Pp_1398 | Uncharacterized protein | 2.254829974 | 1.173018651 | 0.024457517 | 1.611587641 | 157 |
| Q88EY5 | Q88EY5_PSEPK | Pp_4315 | Phenazine biosynthesis protein, PhzF family | 2.238506152 | 1.162536283 | 0.006342962 | 2.197707865 | 158 |
| Q88H65 | Q88H65_PSEPK | Pp_3496 | Uncharacterized protein | 2.233968516 | 1.159608853 | 0.000614859 | 3.211224123 | 159 |
| Q88LX2 | Q88LX2_PSEPK | Wbpm | Polysaccharide biosynthesis protein | 2.233368748 | 1.159221472 | 0.03815228 | 1.418479502 | 160 |
| Q88L40 | PYRD_PSEPK | Pyrd | Dihydroorotate dehydrogenase (quinone) | 2.212030201 | 1.145371083 | 0.005032308 | 2.298232778 | 161 |
| Q88CH9 | Q88CH9_PSEPK | Pp_5202 | Cell division protein ZapA | 2.211509234 | 1.145031266 | 0.039042592 | 1.408461362 | 162 |
| Q88RL3 | Q88RL3_PSEPK | Pp_0116 | Lipoprotein, putative | 2.203760213 | 1.139967255 | 0.006626309 | 2.178728306 | 163 |

| | | | | | | | | |
|---|---|---|---|---|---|---|---|---|
| Q88M18 | Q88M18_PSEPK | Bola | BolA protein | 2.203052263 | 1.13950372 | 0.014608467 | 1.835395347 | 164 |
| Q88MG0 | KDSA1_PSEPK | Kdsa1 | 2-dehydro-3-deoxyphosphooctonate aldolase 1 | 2.201647633 | 1.138583588 | 0.00251023 | 2.600286464 | 165 |
| Q88M88 | Q88M88_PSEPK | Pp_1686 | Glutathione peroxidase | 2.1986474 | 1.136616256 | 0.030383113 | 1.517367732 | 166 |
| Q88RB9 | DAVT_PSEPK | Davt | 5-aminovalerate aminotransferase DavT | 2.193267492 | 1.133081775 | 4.39E-05 | 4.357420525 | 167 |
| Q88EI5 | ASTB_PSEPK | Astb | N-succinylarginine dihydrolase | 2.18889543 | 1.130203035 | 0.006094209 | 2.215082641 | 168 |
| Q88E11 | Q88E11_PSEPK | Pp_4657 | Metallopeptidase, zinc binding | 2.178507595 | 1.123340143 | 0.004196184 | 2.377145505 | 169 |
| Q88HF7 | Q88HF7_PSEPK | Pp_3402 | Uncharacterized protein | 2.176800031 | 1.122208882 | 0.022682745 | 1.644304399 | 170 |
| Q88NX5 | Q88NX5_PSEPK | Pp_1078 | ABC transporter, ATP-binding protein, putative | 2.172767596 | 1.119533869 | 0.047566236 | 1.322701214 | 171 |
| Q88KI3 | Q88KI3_PSEPK | Pp_2307 | Uncharacterized protein | 2.171016474 | 1.118370674 | 0.003450882 | 2.462069852 | 172 |
| Q88NF0 | Q88NF0_PSEPK | Pp_1262 | Transcriptional regulator, LysR family | 2.166592421 | 1.115427779 | 0.012201395 | 1.913590509 | 173 |
| Q88QF1 | Q88QF1_PSEPK | Eutb | Ethanolamine ammonia-lyase, heavy subunit | 2.163416688 | 1.113311565 | 0.022155072 | 1.654526841 | 174 |
| Q88FT9 | Q88FT9_PSEPK | Dsre | DsrE protein | 2.157120547 | 1.109106801 | 0.017090658 | 1.767241217 | 175 |
| Q88NC0 | MOAC_PSEPK | Moac | Cyclic pyranopterin monophosphate synthase accessory protein | 2.154430582 | 1.107306614 | 0.014168982 | 1.848661363 | 176 |
| Q88E84 | Q88E84_PSEPK | Pp_4582 | Uncharacterized protein | 2.149955938 | 1.104307093 | 0.013890887 | 1.857270009 | 177 |
| Q88QI1 | Q88QI1_PSEPK | Pp_0512 | Uncharacterized protein | 2.139368104 | 1.097184737 | 0.006672425 | 2.175716304 | 178 |
| Q88LB1 | Q88LB1_PSEPK | Sbcc | Exonuclease SbcC | 2.135994039 | 1.094907621 | 0.006945749 | 2.158280929 | 179 |
| Q88N76 | MURG_PSEPK | Murg | UDP-N-acetylglucosamine--N-acetylmuramyl-(pentapeptide) pyrophosphoryl-undecaprenol N-acetylglucosamine transferase | 2.135549105 | 1.094607072 | 0.000210081 | 3.677613399 | 180 |
| Q88CY8 | Q88CY8_PSEPK | Glgp | Alpha-1,4 glucan phosphorylase | 2.130371549 | 1.091105067 | 0.047414967 | 1.324084552 | 181 |
| Q88CU3 | RL31_PSEPK | Rpme | 50S ribosomal protein L31 | 2.118607674 | 1.083116453 | 0.003588888 | 2.445040124 | 182 |
| Q88NK5 | Q88NK5_PSEPK | Pp_1202 | Membrane protein, putative | 2.113876085 | 1.079890809 | 0.016677518 | 1.777868587 | 183 |
| Q88RG4 | Q88RG4_PSEPK | Pp_0166 | HlyD family secretion protein | 2.090103566 | 1.063574431 | 0.007051303 | 2.151730621 | 184 |
| Q88R71 | Q88R71_PSEPK | Pp_0263 | Sigma-54 dependent response regulator | 2.088845461 | 1.062705761 | 0.042182054 | 1.374872276 | 185 |
| Q88PV4 | HEMH_PSEPK | Hemh | Ferrochelatase | 2.065835223 | 1.046725185 | 0.049805421 | 1.302723382 | 186 |
| Q88F46 | Q88F46_PSEPK | Ccop-1 | Cbb3-type cytochrome c oxidase subunit | 2.054698127 | 1.038926451 | 0.013849226 | 1.8585745 | 187 |
| Q88NW8 | RNT_PSEPK | Rnt | Ribonuclease T | 2.034385119 | 1.024592814 | 0.001866686 | 2.728928762 | 188 |
| Q88F41 | Q88F41_PSEPK | Ccop-2 | Cbb3-type cytochrome c oxidase subunit | 2.034178739 | 1.024446452 | 0.025556315 | 1.592501766 | 189 |
| Q88CN9 | TYSY_PSEPK | Thya | Thymidylate synthase | 2.029368624 | 1.021030947 | 0.001110105 | 2.954635977 | 190 |

| | | | | | | | | |
|---|---|---|---|---|---|---|---|---|
| Q88KI5 | Q88KI5_PSEPK | Pp_2305 | Uncharacterized protein | 2.01255394 | 1.00902745 | 0.04697223 | 1.328158817 | 191 |
| Q88MS0 | Q88MS0_PSEPK | Pp_1499 | Uncharacterized protein | 2.010831969 | 1.007792531 | 0.036949849 | 1.43238733 | 192 |
| Q88RV9 | Q88RV9_PSEPK | Pp_0020 | Uncharacterized protein | 2.00743509 | 1.005353339 | 0.002031027 | 2.692284393 | 193 |

Supplementary Table S5. List of plasmids used in this study*

| Plasmid No. | Plasmid description | JBEI Accession No. |
|---|---|---|
| pIY993 | pJET-J23119-sfGFP-dCas9 handle and terminator | JBx_249165 |
| pIY994 | pJET-Carb-RFP_NT1 | JBx_249167 |
| pIY995 | pJET-Carb-PP_4194_NT1 | JBx_249169 |
| pIY996 | pJET-Carb-PP_1010 | JBx_249171 |
| pIY997 | pJET-Carb-nontarget | JBx_249173 |
| pIY1021 | pJET-Carb-PP_3540 | JBx_249175 |
| pIY1022 | pJET-Carb-PP_1649 | JBx_249177 |
| pIY1023 | pJET-Carb-PP_4549 | JBx_249179 |
| pIY1024 | pJET-Carb-PP_4550 | JBx_249181 |
| pIY1025 | pJET-Carb-PP_0368 | JBx_249183 |
| pIY1026 | pJET-Carb-PP_0370 | JBx_249185 |
| pIY1027 | pJET-Carb-PP_2136 | JBx_249187 |
| pIY1028 | pJET-Carb-PP_4740_NT1 | JBx_249189 |
| pIY1029 | pJET-Carb-PP_4740_NT2 | JBx_249191 |
| pIY1030 | pJET-Carb-PP_4740_NT3 | JBx_249193 |
| pIY1031 | pJET-Carb-PP_0774 | JBx_249195 |
| pIY1032 | pJET-Carb-PP_0362 | JBx_249197 |
| pIY1033 | pJET-Carb-PP_0437 | JBx_249199 |
| pIY1034 | pJET-Carb-PP_2589 | JBx_249201 |
| pIY1035 | pJET-Carb-PP_3744 | JBx_249203 |
| pIY1036 | pJET-Carb-PP_4194_NT2 | JBx_249205 |
| pIY1037 | pJET-Carb-PP_4194_NT3 | JBx_249207 |
| pIY1038 | pJET-Carb-PP_1607_NT1 | JBx_249209 |
| pIY1039 | pJET-Carb-PP_1607_NT2 | JBx_249211 |
| pIY1040 | pJET-Carb-PP_1607_NT3 | JBx_249213 |
| pIY1041 | pJET-Carb-PP_1607_NT4 | JBx_249215 |
| pIY1042 | pJET-Carb-PP_1444 | JBx_249217 |
| pIY1043 | pJET-Carb-PP_0338 | JBx_249219 |
| pIY1044 | pJET-Carb-PP_3394 | JBx_249221 |
| pIY1045 | pJET-Carb-PP_3071 | JBx_249223 |
| pIY1046 | pJET-Carb-PP_0582 | JBx_249225 |
| pIY1047 | pJET-Carb-PP_3355 | JBx_249227 |
| pIY1048 | pJET-Carb-PP_3754 | JBx_249229 |
| pIY1049 | pJET-Carb-PP_3282 | JBx_249231 |
| pIY1050 | pJET-Carb-PP_3755 | JBx_249233 |
| pIY1051 | pJET-Carb-PP_1157 | JBx_249235 |
| pIY1052 | pJET-Carb-PP_1394 | JBx_249237 |
| pIY1053 | pJET-Carb-PP_3365 | JBx_249239 |
| pIY1054 | pJET-Carb-PP_4679 | JBx_249241 |
| pIY1055 | pJET-Carb-PP_4678 | JBx_249243 |
| pIY1056 | pJET-Carb-PP_1777 | JBx_249245 |

| | | |
|---|---|---|
| pIY1057 | pJET-Carb-PP_3578 | JBx_249247 |
| pIY1058 | pJET-Carb-PP_5288 | JBx_249249 |
| pIY1059 | pJET-Carb-PP_0597 | JBx_249251 |
| pIY1060 | pJET-Carb-PP_4667 | JBx_249253 |
| pIY1082 | pJET-Carb-PP_3122 | JBx_249255 |
| pIY1083 | pJET-Carb-PP_3123 | JBx_249257 |
| pIY1084 | pJET-Carb-PP_2137 | JBx_249259 |
| pIY1085 | pJET-Carb-PP_3280 | JBx_249261 |
| pIY1086 | pJET-Carb-PP_2471 | JBx_249263 |
| pIY1087 | pJET-Carb-PP_4737 | JBx_249265 |
| pIY1088 | pJET-Carb-PP_4894 | JBx_249267 |
| pIY1089 | pJET-Carb-PP_2213 | JBx_249269 |
| pIY1090 | pJET-Carb-PP_4487 | JBx_249271 |
| pIY1091 | pJET-Carb-PP_4702 | JBx_249273 |
| pIY1092 | pJET-Carb-PP_0528 | JBx_249275 |
| pIY1093 | pJET-Carb-PP_0548 | JBx_249277 |
| pIY1094 | pJET-Carb-PP_4636 | JBx_249279 |
| pIY1139 | pJET-Carb-PP_0527 | JBx_249281 |
| pIY1140 | pJET-Carb-PP_2533 | JBx_249283 |
| pIY1141 | pJET-Carb-PP_5155 | JBx_249285 |
| pIY1142 | pJET-Carb-PP_1444_NT2 | JBx_249287 |
| pIY1143 | pJET-Carb-PP_0339 | JBx_249289 |
| pIY1144 | pJET-Carb-PP_2051 | JBx_249291 |
| pIY1145 | pJET-Carb-PP_2215 | JBx_249293 |
| pIY1146 | pJET-Carb-PP_1866 | JBx_249295 |
| pIY1147 | pJET-Carb-PP_2324 | JBx_249297 |
| pIY1148 | pJET-Carb-PP_3080 | JBx_249299 |
| pIY1149 | pJET-Carb-PP_5351 | JBx_249301 |
| pIY1150 | pJET-Carb-PP_4736 | JBx_249303 |
| pIY1151 | pJET-Carb-PP_1773 | JBx_249305 |
| pIY1152 | pJET-Carb-PP_0356 | JBx_249307 |
| pIY1153 | pJET-Carb-PP_4379 | JBx_249309 |
| pIY1154 | pJET-Carb-PP_1319 | JBx_249311 |
| pIY1155 | pJET-Carb-PP_5258 | JBx_249313 |
| pIY1156 | pJET-Carb-PP_1616 | JBx_249315 |
| pIY1157 | pJET-Carb-PP_2803 | JBx_249317 |
| pIY1217 | pJET-Carb-PP_5420 | JBx_249319 |
| pIY1218 | pJET-Carb-PP_5419 | JBx_249321 |
| pIY1219 | pJET-Carb-PP_5417 | JBx_249323 |
| pIY1220 | pJET-Carb-PP_5416 | JBx_249325 |
| pIY1221 | pJET-Carb-PP_0806 | JBx_249327 |
| pIY1222 | pJET-Carb-PP_0168 | JBx_249329 |
| pIY1223 | pJET-Carb-PP_5335 | JBx_249331 |

| | | |
|---|---|---|
| pIY1224 | pJET-Carb-PP_4119 | JBx_249333 |
| pIY1225 | pJET-Carb-PP_4120 | JBx_249335 |
| pIY1226 | pJET-Carb-PP_4121 | JBx_249337 |
| pIY1227 | pJET-Carb-PP_4122 | JBx_249339 |
| pIY1228 | pJET-Carb-PP_4123 | JBx_249341 |
| pIY1229 | pJET-Carb-PP_4124 | JBx_249343 |
| pIY1230 | pJET-Carb-PP_4129 | JBx_249345 |
| pIY1231 | pJET-Carb-PP_4130 | JBx_249347 |
| pIY1232 | pJET-Carb-PP_4128 | JBx_249349 |
| pIY1233 | pJET-Carb-PP_0999 | JBx_249351 |
| pIY1234 | pJET-Carb-PP_1031 | JBx_249353 |
| pIY1235 | pJET-Carb-PP_1770 | JBx_249355 |
| pIY1236 | pJET-Carb-PP_1664 | JBx_249357 |
| pIY1237 | pJET-Carb-PP_4175 | JBx_249359 |
| pIY1238 | pJET-Carb-PP_4174 | JBx_249361 |
| pIY1239 | pJET-Carb-PP_4189 | JBx_249363 |
| pIY1240 | pJET-Carb-PP_4188 | JBx_249365 |
| pIY1241 | pJET-Carb-PP_4187 | JBx_249367 |
| pIY1242 | pJET-Carb-PP_1771 | JBx_249369 |
| pIY1243 | pJET-Carb-PP_0691 | JBx_249371 |
| pIY1244 | pJET-Carb-PP_0001 | JBx_249373 |
| pIY1245 | pJET-Carb-PP_4116 | JBx_249375 |
| pIY1246 | pJET-Carb-PP_4043 | JBx_249377 |
| pIY1247 | pJET-Carb-PP_4862 | JBx_249379 |
| pIY1248 | pJET-Carb-PP_0103 | JBx_249381 |
| pIY1249 | pJET-Carb-PP_0104 | JBx_249383 |
| pIY1250 | pJET-Carb-PP_0105 | JBx_249385 |
| pIY1251 | pJET-Carb-PP_0106 | JBx_249387 |
| pIY1252 | pJET-Carb-PP_1317 | JBx_249389 |
| pIY1253 | pJET-Carb-PP_1318 | JBx_249391 |
| pIY1406 | pJET-Carb-PP_1319 | JBx_249393 |
| pIY1407 | pJET-Carb-PP_0812 | JBx_249395 |
| pIY1408 | pJET-Carb-PP_0813 | JBx_249397 |
| pIY1409 | pJET-Carb-PP_0814 | JBx_249399 |
| pIY1410 | pJET-Carb-PP_0815 | JBx_249401 |
| pIY1411 | pJET-Carb-PP_1023 | JBx_249403 |
| pIY1412 | pJET-Carb-PP_1240 | JBx_249405 |
| pIY1413 | pJET-Carb-PP_5128 | JBx_249407 |
| pIY1414 | pJET-Carb-PP_5064 | JBx_249409 |
| pIY1415 | pJET-Carb-PP_2095 | JBx_249411 |
| pIY1416 | pJET-Carb-PP_1769 | JBx_249413 |
| pIY1417 | pJET-Carb-PP_4635 | JBx_249415 |
| pIY1418 | pJET-Carb-PP_1610 | JBx_249417 |

| | | |
|---|---|---|
| pIY1419 | pJET-Carb-PP_5186 | JBx_249419 |
| pIY1420 | pJET-Carb-PP_4161 | JBx_249421 |
| pIY1421 | pJET-Carb-PP_4160 | JBx_249423 |
| pIY1422 | pJET-Carb-PP_4159 | JBx_249425 |
| pIY1423 | pJET-Carb-PP_0658 | JBx_249427 |
| pIY1424 | pJET-Carb-PP_1506 | JBx_249429 |
| pIY1425 | pJET-Carb-PP_1620 | JBx_249431 |
| pIY1426 | pJET-Carb-PP_4266 | JBx_249433 |
| pIY1427 | pJET-Carb-PP_1457 | JBx_249435 |
| pIY1428 | pJET-Carb-PP_5203 | JBx_249437 |
| pIY1429 | pJET-Carb-PP_0227 | JBx_249439 |
| pIY1430 | pJET-Carb-PP_0226 | JBx_249441 |
| pIY1431 | pJET-Carb-PP_0225 | JBx_249443 |
| pIY1432 | pJET-Carb-PP_4651 | JBx_249445 |
| pIY1433 | pJET-Carb-PP_4650 | JBx_249447 |
| pIY1434 | pJET-Carb-PP_4193 | JBx_249449 |
| pIY1435 | pJET-Carb-PP_4192 | JBx_249451 |
| pIY1436 | pJET-Carb-PP_4191 | JBx_249453 |
| pIY1437 | pJET-Carb-PP_4190 | JBx_249455 |
| pIY1438 | pJET-Carb-PP_0897 | JBx_249457 |
| pIY1439 | pJET-Carb-PP_0944 | JBx_249459 |
| pIY1440 | pJET-Carb-PP_1755 | JBx_249461 |
| pIY1441 | pJET-Carb-PP_0654 | JBx_249463 |
| pIY1442 | pJET-Carb-PP_0751 | JBx_249465 |
| pIY1443 | pJET-Carb-PP_2925 | JBx_249467 |
| pIY1444 | pJET-Carb-PP_1251 | JBx_249469 |
| pIY1445 | pJET-Carb-PP_2112 | JBx_249471 |
| pIY1446 | pJET-Carb-PP_2339 | JBx_249473 |
| pIY1447 | pJET-Carb-PP_4011 | JBx_249475 |
| pIY1448 | pJET-Carb-PP_4012 | JBx_249477 |
| pIY1449 | pJET-Carb-PP_4185 | JBx_249479 |
| pIY1450 | pJET-Carb-PP_4186 | JBx_249481 |
| pIY1572 | pJET-Carb-BFP | JBx_249483 |
| pIY1573 | pJET-Carb-PP_4960 | JBx_249485 |
| pIY1574 | pJET-Carb-PP_1791 | JBx_249487 |
| pIY1575 | pJET-Carb-PP_2871 | JBx_249489 |
| pIY1576 | pJET-Carb-PP_3224 | JBx_249491 |
| pIY1577 | pJET-Carb-PP_4715 | JBx_249493 |
| pIY1590 | pJET-Carb-PP_0297 | JBx_249495 |
| pIY1591 | pJET-Carb-PP_0987 | JBx_249497 |
| pIY1592 | pJET-Carb-PP_2930 | JBx_249499 |
| pIY1593 | pJET-Carb-PP_3144 | JBx_249501 |
| pIY1594 | pJET-Carb-PP_3191 | JBx_249503 |

| | | |
|---|---|---|
| pIY1595 | pJET-Carb-PP_3446 | JBx_249505 |
| pIY1596 | pJET-Carb-PP_5149 | JBx_249507 |
| pIY1597 | pJET-Carb-PP_0731 | JBx_249509 |
| pIY1598 | pJET-Carb-PP_3664 | JBx_249511 |
| pIY1599 | pJET-Carb-PP_4677 | JBx_249513 |
| pIY1600 | pJET-Carb-PP_0082 | JBx_249515 |
| pIY1601 | pJET-Carb-PP_0083 | JBx_249517 |
| pIY1602 | pJET-Carb-PP_0988 | JBx_249519 |
| pIY1603 | pJET-Carb-PP_5192 | JBx_249521 |
| pIY1636 | pJET-Carb-PP_0321 | JBx_249523 |
| pIY1637 | pJET-Carb-PP_1808 | JBx_249525 |
| pIY1638 | pJET-Carb-PP_4701 | JBx_249527 |
| pIY1639 | pJET-Carb-PP_1011 | JBx_249529 |
| pIY1640 | pJET-Carb-PP_1156 | JBx_249531 |
| pIY1641 | pJET-Carb-PP_2149 | JBx_249533 |
| pIY1642 | pJET-Carb-PP_4963 | JBx_249535 |
| pIY1643 | pJET-Carb-PP_5056 | JBx_249537 |
| pIY1644 | pJET-Carb-PP_1612 | JBx_249539 |
| pIY1645 | pJET-Carb-PP_1362 | JBx_249541 |
| pIY1754 | pJET-Carb-PP_5075 | JBx_249543 |
| pIY1755 | pJET-Carb-PP_5076 | JBx_249545 |
| pIY1756 | pJET-Carb-PP_0675 | JBx_249547 |
| pIY1757 | pJET-Carb-PP_2080 | JBx_249549 |
| pIY1758 | pJET-Carb-PP_5046_NT2 | JBx_249551 |
| pIY1759 | pJET-Carb-PP_3591 | JBx_249553 |
| pIY1760 | pJET-Carb-PP_2336 | JBx_249555 |
| pIY1761 | pJET-Carb-PP_2652 | JBx_249557 |
| pIY1762 | pJET-Carb-PP_3662 | JBx_249559 |
| pIY1763 | pJET-Carb-PP_4012_NT2 | JBx_249561 |
| pIY1764 | pJET-Carb-PP_0356 | JBx_249563 |
| pIY989 | pRSF1010-Gm-NagR-pNagAa-dCas9-thrLABC-sfGFP | JBx_249567 |
| pIY1003 | pRSF1010-Gm-NagR-pNagAa-dCas9-thrLABC-RFP_NT1 | JBx_249569 |
| pIY1004 | pRSF1010-Gm-NagR-pNagAa-dCas9-thrLABC-PP_4194_NT1 | JBx_249571 |
| pIY1005 | pRSF1010-Gm-NagR-pNagAa-dCas9-thrLABC-PP_1010 | JBx_249573 |
| pIY1006 | pRSF1010-Gm-NagR-pNagAa-dCas9-thrLABC-nontarget | JBx_249575 |
| pIY1061 | pRSF1010-Gm-NagR-pNagAa-dCas9-thrLABC-PP_3540 | JBx_249577 |
| pIY1062 | pRSF1010-Gm-NagR-pNagAa-dCas9-thrLABC-PP_1649 | JBx_249579 |
| pIY1063 | pRSF1010-Gm-NagR-pNagAa-dCas9-thrLABC-PP_4549 | JBx_249581 |
| pIY1064 | pRSF1010-Gm-NagR-pNagAa-dCas9-thrLABC-PP_4550 | JBx_249583 |
| pIY1065 | pRSF1010-Gm-NagR-pNagAa-dCas9-thrLABC-PP_0368 | JBx_249585 |
| pIY1066 | pRSF1010-Gm-NagR-pNagAa-dCas9-thrLABC-PP_0370 | JBx_249587 |
| pIY1067 | pRSF1010-Gm-NagR-pNagAa-dCas9-thrLABC-PP_2136 | JBx_249589 |
| pIY1068 | pRSF1010-Gm-NagR-pNagAa-dCas9-thrLABC-PP_4740_NT1 | JBx_249591 |

| | | |
|---|---|---|
| pIY1069 | pRSF1010-Gm-NagR-pNagAa-dCas9-thrLABC-PP_4740_NT2 | JBx_249593 |
| pIY1070 | pRSF1010-Gm-NagR-pNagAa-dCas9-thrLABC-PP_4740_NT3 | JBx_249595 |
| pIY1071 | pRSF1010-Gm-NagR-pNagAa-dCas9-thrLABC-PP_0774 | JBx_249597 |
| pIY1072 | pRSF1010-Gm-NagR-pNagAa-dCas9-thrLABC-PP_0362 | JBx_249599 |
| pIY1073 | pRSF1010-Gm-NagR-pNagAa-dCas9-thrLABC-PP_0437 | JBx_249601 |
| pIY1074 | pRSF1010-Gm-NagR-pNagAa-dCas9-thrLABC-PP_2589 | JBx_249603 |
| pIY1075 | pRSF1010-Gm-NagR-pNagAa-dCas9-thrLABC-PP_3744 | JBx_249605 |
| pIY1076 | pRSF1010-Gm-NagR-pNagAa-dCas9-thrLABC-PP_4194_NT2 | JBx_249607 |
| pIY1077 | pRSF1010-Gm-NagR-pNagAa-dCas9-thrLABC-PP_4194_NT3 | JBx_249609 |
| pIY1078 | pRSF1010-Gm-NagR-pNagAa-dCas9-thrLABC-PP_1607_NT1 | JBx_249611 |
| pIY1079 | pRSF1010-Gm-NagR-pNagAa-dCas9-thrLABC-PP_1607_NT2 | JBx_249613 |
| pIY1080 | pRSF1010-Gm-NagR-pNagAa-dCas9-thrLABC-PP_1607_NT3 | JBx_249615 |
| pIY1081 | pRSF1010-Gm-NagR-pNagAa-dCas9-thrLABC-PP_1607_NT4 | JBx_249617 |
| pIY1099 | pRSF1010-Gm-NagR-pNagAa-dCas9-thrLABC-PP_1444 | JBx_249619 |
| pIY1100 | pRSF1010-Gm-NagR-pNagAa-dCas9-thrLABC-PP_0338 | JBx_249621 |
| pIY1101 | pRSF1010-Gm-NagR-pNagAa-dCas9-thrLABC-PP_3394 | JBx_249623 |
| pIY1102 | pRSF1010-Gm-NagR-pNagAa-dCas9-thrLABC-PP_3071 | JBx_249625 |
| pIY1103 | pRSF1010-Gm-NagR-pNagAa-dCas9-thrLABC-PP_0582 | JBx_249627 |
| pIY1104 | pRSF1010-Gm-NagR-pNagAa-dCas9-thrLABC-PP_3355 | JBx_249629 |
| pIY1105 | pRSF1010-Gm-NagR-pNagAa-dCas9-thrLABC-PP_3754 | JBx_249631 |
| pIY1106 | pRSF1010-Gm-NagR-pNagAa-dCas9-thrLABC-PP_3282 | JBx_249633 |
| pIY1107 | pRSF1010-Gm-NagR-pNagAa-dCas9-thrLABC-PP_3755 | JBx_249635 |
| pIY1108 | pRSF1010-Gm-NagR-pNagAa-dCas9-thrLABC-PP_1157 | JBx_249637 |
| pIY1109 | pRSF1010-Gm-NagR-pNagAa-dCas9-thrLABC-PP_1394 | JBx_249639 |
| pIY1110 | pRSF1010-Gm-NagR-pNagAa-dCas9-thrLABC-PP_3365 | JBx_249641 |
| pIY1111 | pRSF1010-Gm-NagR-pNagAa-dCas9-thrLABC-PP_4679 | JBx_249643 |
| pIY1112 | pRSF1010-Gm-NagR-pNagAa-dCas9-thrLABC-PP_4678 | JBx_249645 |
| pIY1113 | pRSF1010-Gm-NagR-pNagAa-dCas9-thrLABC-PP_1777 | JBx_249647 |
| pIY1114 | pRSF1010-Gm-NagR-pNagAa-dCas9-thrLABC-PP_3578 | JBx_249649 |
| pIY1115 | pRSF1010-Gm-NagR-pNagAa-dCas9-thrLABC-PP_5288 | JBx_249651 |
| pIY1116 | pRSF1010-Gm-NagR-pNagAa-dCas9-thrLABC-PP_0597 | JBx_249653 |
| pIY1117 | pRSF1010-Gm-NagR-pNagAa-dCas9-thrLABC-PP_4667 | JBx_249655 |
| pIY1124 | pRSF1010-Gm-NagR-pNagAa-dCas9-thrLABC-PP_3122 | JBx_249657 |
| pIY1125 | pRSF1010-Gm-NagR-pNagAa-dCas9-thrLABC-PP_3123 | JBx_249659 |
| pIY1126 | pRSF1010-Gm-NagR-pNagAa-dCas9-thrLABC-PP_2137 | JBx_249661 |
| pIY1127 | pRSF1010-Gm-NagR-pNagAa-dCas9-thrLABC-PP_3280 | JBx_249663 |
| pIY1128 | pRSF1010-Gm-NagR-pNagAa-dCas9-thrLABC-PP_2471 | JBx_249665 |
| pIY1129 | pRSF1010-Gm-NagR-pNagAa-dCas9-thrLABC-PP_4737 | JBx_249667 |
| pIY1130 | pRSF1010-Gm-NagR-pNagAa-dCas9-thrLABC-PP_4894 | JBx_249669 |
| pIY1131 | pRSF1010-Gm-NagR-pNagAa-dCas9-thrLABC-PP_2213 | JBx_249671 |
| pIY1132 | pRSF1010-Gm-NagR-pNagAa-dCas9-thrLABC-PP_4487 | JBx_249673 |
| pIY1133 | pRSF1010-Gm-NagR-pNagAa-dCas9-thrLABC-PP_4702 | JBx_249675 |
| pIY1134 | pRSF1010-Gm-NagR-pNagAa-dCas9-thrLABC-PP_0528 | JBx_249677 |

| | | |
|---|---|---|
| pIY1135 | pRSF1010-Gm-NagR-pNagAa-dCas9-thrLABC-PP_0548 | JBx_249679 |
| pIY1136 | pRSF1010-Gm-NagR-pNagAa-dCas9-thrLABC-PP_4636 | JBx_249681 |
| pIY1194 | pRSF1010-Gm-NagR-pNagAa-dCas9-thrLABC-PP_0527 | JBx_249683 |
| pIY1195 | pRSF1010-Gm-NagR-pNagAa-dCas9-thrLABC-PP_2533 | JBx_249685 |
| pIY1196 | pRSF1010-Gm-NagR-pNagAa-dCas9-thrLABC-PP_5155 | JBx_249687 |
| pIY1197 | pRSF1010-Gm-NagR-pNagAa-dCas9-thrLABC-PP_1444_NT2 | JBx_249689 |
| pIY1198 | pRSF1010-Gm-NagR-pNagAa-dCas9-thrLABC-PP_0339 | JBx_249691 |
| pIY1199 | pRSF1010-Gm-NagR-pNagAa-dCas9-thrLABC-PP_2051 | JBx_249693 |
| pIY1200 | pRSF1010-Gm-NagR-pNagAa-dCas9-thrLABC-PP_2215 | JBx_249695 |
| pIY1201 | pRSF1010-Gm-NagR-pNagAa-dCas9-thrLABC-PP_1866 | JBx_249697 |
| pIY1202 | pRSF1010-Gm-NagR-pNagAa-dCas9-thrLABC-PP_2324 | JBx_249699 |
| pIY1203 | pRSF1010-Gm-NagR-pNagAa-dCas9-thrLABC-PP_3080 | JBx_249701 |
| pIY1204 | pRSF1010-Gm-NagR-pNagAa-dCas9-thrLABC-PP_5351 | JBx_249703 |
| pIY1205 | pRSF1010-Gm-NagR-pNagAa-dCas9-thrLABC-PP_4736 | JBx_249705 |
| pIY1206 | pRSF1010-Gm-NagR-pNagAa-dCas9-thrLABC-PP_1773 | JBx_249707 |
| pIY1207 | pRSF1010-Gm-NagR-pNagAa-dCas9-thrLABC-PP_0356 | JBx_249709 |
| pIY1208 | pRSF1010-Gm-NagR-pNagAa-dCas9-thrLABC-PP_4379 | JBx_249711 |
| pIY1209 | pRSF1010-Gm-NagR-pNagAa-dCas9-thrLABC-PP_1319 | JBx_249713 |
| pIY1210 | pRSF1010-Gm-NagR-pNagAa-dCas9-thrLABC-PP_5258 | JBx_249715 |
| pIY1211 | pRSF1010-Gm-NagR-pNagAa-dCas9-thrLABC-PP_1616 | JBx_249717 |
| pIY1212 | pRSF1010-Gm-NagR-pNagAa-dCas9-thrLABC-PP_2803 | JBx_249719 |
| pIY1266 | pRSF1010-Gm-NagR-pNagAa-dCas9-thrLABC-PP_5420 | JBx_249721 |
| pIY1267 | pRSF1010-Gm-NagR-pNagAa-dCas9-thrLABC-PP_5419 | JBx_249723 |
| pIY1268 | pRSF1010-Gm-NagR-pNagAa-dCas9-thrLABC-PP_5417 | JBx_249725 |
| pIY1269 | pRSF1010-Gm-NagR-pNagAa-dCas9-thrLABC-PP_5416 | JBx_249727 |
| pIY1270 | pRSF1010-Gm-NagR-pNagAa-dCas9-thrLABC-PP_0806 | JBx_249729 |
| pIY1271 | pRSF1010-Gm-NagR-pNagAa-dCas9-thrLABC-PP_0168 | JBx_249731 |
| pIY1272 | pRSF1010-Gm-NagR-pNagAa-dCas9-thrLABC-PP_5335 | JBx_249733 |
| pIY1273 | pRSF1010-Gm-NagR-pNagAa-dCas9-thrLABC-PP_4119 | JBx_249735 |
| pIY1274 | pRSF1010-Gm-NagR-pNagAa-dCas9-thrLABC-PP_4120 | JBx_249737 |
| pIY1275 | pRSF1010-Gm-NagR-pNagAa-dCas9-thrLABC-PP_4121 | JBx_249739 |
| pIY1276 | pRSF1010-Gm-NagR-pNagAa-dCas9-thrLABC-PP_4122 | JBx_249741 |
| pIY1277 | pRSF1010-Gm-NagR-pNagAa-dCas9-thrLABC-PP_4123 | JBx_249743 |
| pIY1278 | pRSF1010-Gm-NagR-pNagAa-dCas9-thrLABC-PP_4124 | JBx_249745 |
| pIY1279 | pRSF1010-Gm-NagR-pNagAa-dCas9-thrLABC-PP_4129 | JBx_249747 |
| pIY1280 | pRSF1010-Gm-NagR-pNagAa-dCas9-thrLABC-PP_4130 | JBx_249749 |
| pIY1281 | pRSF1010-Gm-NagR-pNagAa-dCas9-thrLABC-PP_4128 | JBx_249751 |
| pIY1282 | pRSF1010-Gm-NagR-pNagAa-dCas9-thrLABC-PP_0999 | JBx_249753 |
| pIY1283 | pRSF1010-Gm-NagR-pNagAa-dCas9-thrLABC-PP_1031 | JBx_249755 |
| pIY1284 | pRSF1010-Gm-NagR-pNagAa-dCas9-thrLABC-PP_1770 | JBx_249757 |
| pIY1285 | pRSF1010-Gm-NagR-pNagAa-dCas9-thrLABC-PP_1664 | JBx_249759 |
| pIY1286 | pRSF1010-Gm-NagR-pNagAa-dCas9-thrLABC-PP_4175 | JBx_249761 |
| pIY1287 | pRSF1010-Gm-NagR-pNagAa-dCas9-thrLABC-PP_4174 | JBx_249763 |

| | | |
|---|---|---|
| pIY1288 | pRSF1010-Gm-NagR-pNagAa-dCas9-thrLABC-PP_4189 | JBx_249765 |
| pIY1289 | pRSF1010-Gm-NagR-pNagAa-dCas9-thrLABC-PP_4188 | JBx_249767 |
| pIY1290 | pRSF1010-Gm-NagR-pNagAa-dCas9-thrLABC-PP_4187 | JBx_249769 |
| pIY1291 | pRSF1010-Gm-NagR-pNagAa-dCas9-thrLABC-PP_1771 | JBx_249771 |
| pIY1292 | pRSF1010-Gm-NagR-pNagAa-dCas9-thrLABC-PP_0691 | JBx_249773 |
| pIY1293 | pRSF1010-Gm-NagR-pNagAa-dCas9-thrLABC-PP_0001 | JBx_249775 |
| pIY1294 | pRSF1010-Gm-NagR-pNagAa-dCas9-thrLABC-PP_4116 | JBx_249777 |
| pIY1295 | pRSF1010-Gm-NagR-pNagAa-dCas9-thrLABC-PP_4043 | JBx_249779 |
| pIY1296 | pRSF1010-Gm-NagR-pNagAa-dCas9-thrLABC-PP_4862 | JBx_249781 |
| pIY1297 | pRSF1010-Gm-NagR-pNagAa-dCas9-thrLABC-PP_0103 | JBx_249783 |
| pIY1298 | pRSF1010-Gm-NagR-pNagAa-dCas9-thrLABC-PP_0104 | JBx_249785 |
| pIY1299 | pRSF1010-Gm-NagR-pNagAa-dCas9-thrLABC-PP_0105 | JBx_249787 |
| pIY1300 | pRSF1010-Gm-NagR-pNagAa-dCas9-thrLABC-PP_0106 | JBx_249789 |
| pIY1301 | pRSF1010-Gm-NagR-pNagAa-dCas9-thrLABC-PP_1317 | JBx_249791 |
| pIY1302 | pRSF1010-Gm-NagR-pNagAa-dCas9-thrLABC-PP_1318 | JBx_249793 |
| pIY1451 | pRSF1010-Gm-NagR-pNagAa-dCas9-thrLABC-PP_1319 | JBx_249795 |
| pIY1452 | pRSF1010-Gm-NagR-pNagAa-dCas9-thrLABC-PP_0812 | JBx_249797 |
| pIY1453 | pRSF1010-Gm-NagR-pNagAa-dCas9-thrLABC-PP_0813 | JBx_249799 |
| pIY1454 | pRSF1010-Gm-NagR-pNagAa-dCas9-thrLABC-PP_0814 | JBx_249801 |
| pIY1455 | pRSF1010-Gm-NagR-pNagAa-dCas9-thrLABC-PP_0815 | JBx_249803 |
| pIY1456 | pRSF1010-Gm-NagR-pNagAa-dCas9-thrLABC-PP_1023 | JBx_249805 |
| pIY1457 | pRSF1010-Gm-NagR-pNagAa-dCas9-thrLABC-PP_1240 | JBx_249807 |
| pIY1458 | pRSF1010-Gm-NagR-pNagAa-dCas9-thrLABC-PP_5128 | JBx_249809 |
| pIY1459 | pRSF1010-Gm-NagR-pNagAa-dCas9-thrLABC-PP_5064 | JBx_249811 |
| pIY1460 | pRSF1010-Gm-NagR-pNagAa-dCas9-thrLABC-PP_2095 | JBx_249813 |
| pIY1461 | pRSF1010-Gm-NagR-pNagAa-dCas9-thrLABC-PP_1769 | JBx_249815 |
| pIY1462 | pRSF1010-Gm-NagR-pNagAa-dCas9-thrLABC-PP_4635 | JBx_249817 |
| pIY1463 | pRSF1010-Gm-NagR-pNagAa-dCas9-thrLABC-PP_1610 | JBx_249819 |
| pIY1464 | pRSF1010-Gm-NagR-pNagAa-dCas9-thrLABC-PP_5186 | JBx_249821 |
| pIY1465 | pRSF1010-Gm-NagR-pNagAa-dCas9-thrLABC-PP_4161 | JBx_249823 |
| pIY1466 | pRSF1010-Gm-NagR-pNagAa-dCas9-thrLABC-PP_4160 | JBx_249825 |
| pIY1467 | pRSF1010-Gm-NagR-pNagAa-dCas9-thrLABC-PP_4159 | JBx_249827 |
| pIY1468 | pRSF1010-Gm-NagR-pNagAa-dCas9-thrLABC-PP_0658 | JBx_249829 |
| pIY1469 | pRSF1010-Gm-NagR-pNagAa-dCas9-thrLABC-PP_1506 | JBx_249831 |
| pIY1470 | pRSF1010-Gm-NagR-pNagAa-dCas9-thrLABC-PP_1620 | JBx_249833 |
| pIY1471 | pRSF1010-Gm-NagR-pNagAa-dCas9-thrLABC-PP_4266 | JBx_249835 |
| pIY1472 | pRSF1010-Gm-NagR-pNagAa-dCas9-thrLABC-PP_1457 | JBx_249837 |
| pIY1473 | pRSF1010-Gm-NagR-pNagAa-dCas9-thrLABC-PP_5203 | JBx_249839 |
| pIY1474 | pRSF1010-Gm-NagR-pNagAa-dCas9-thrLABC-PP_0227 | JBx_249841 |
| pIY1475 | pRSF1010-Gm-NagR-pNagAa-dCas9-thrLABC-PP_0226 | JBx_249843 |
| pIY1476 | pRSF1010-Gm-NagR-pNagAa-dCas9-thrLABC-PP_0225 | JBx_249845 |
| pIY1477 | pRSF1010-Gm-NagR-pNagAa-dCas9-thrLABC-PP_4651 | JBx_249847 |
| pIY1478 | pRSF1010-Gm-NagR-pNagAa-dCas9-thrLABC-PP_4650 | JBx_249849 |

| | | |
|---|---|---|
| pIY1479 | pRSF1010-Gm-NagR-pNagAa-dCas9-thrLABC-PP_4193 | JBx_249851 |
| pIY1480 | pRSF1010-Gm-NagR-pNagAa-dCas9-thrLABC-PP_4192 | JBx_249853 |
| pIY1481 | pRSF1010-Gm-NagR-pNagAa-dCas9-thrLABC-PP_4191 | JBx_249855 |
| pIY1482 | pRSF1010-Gm-NagR-pNagAa-dCas9-thrLABC-PP_4190 | JBx_249857 |
| pIY1483 | pRSF1010-Gm-NagR-pNagAa-dCas9-thrLABC-PP_0897 | JBx_249859 |
| pIY1484 | pRSF1010-Gm-NagR-pNagAa-dCas9-thrLABC-PP_0944 | JBx_249861 |
| pIY1485 | pRSF1010-Gm-NagR-pNagAa-dCas9-thrLABC-PP_1755 | JBx_249863 |
| pIY1486 | pRSF1010-Gm-NagR-pNagAa-dCas9-thrLABC-PP_0654 | JBx_249865 |
| pIY1487 | pRSF1010-Gm-NagR-pNagAa-dCas9-thrLABC-PP_0751 | JBx_249867 |
| pIY1488 | pRSF1010-Gm-NagR-pNagAa-dCas9-thrLABC-PP_2925 | JBx_249869 |
| pIY1489 | pRSF1010-Gm-NagR-pNagAa-dCas9-thrLABC-PP_1251 | JBx_249871 |
| pIY1490 | pRSF1010-Gm-NagR-pNagAa-dCas9-thrLABC-PP_2112 | JBx_249873 |
| pIY1491 | pRSF1010-Gm-NagR-pNagAa-dCas9-thrLABC-PP_2339 | JBx_249875 |
| pIY1492 | pRSF1010-Gm-NagR-pNagAa-dCas9-thrLABC-PP_4011 | JBx_249877 |
| pIY1493 | pRSF1010-Gm-NagR-pNagAa-dCas9-thrLABC-PP_4012 | JBx_249879 |
| pIY1494 | pRSF1010-Gm-NagR-pNagAa-dCas9-thrLABC-PP_4185 | JBx_249881 |
| pIY1495 | pRSF1010-Gm-NagR-pNagAa-dCas9-thrLABC-PP_4186 | JBx_249883 |
| **pIY1581** | pRSF1010-Gm-NagR-pNagAa-dCas9-thrLABC-BFP | JBx_249885 |
| pIY1582 | pRSF1010-Gm-NagR-pNagAa-dCas9-thrLABC-PP_4960 | JBx_249887 |
| pIY1583 | pRSF1010-Gm-NagR-pNagAa-dCas9-thrLABC-PP_2871 | JBx_249889 |
| pIY1604 | pRSF1010-Gm-NagR-pNagAa-dCas9-thrLABC-PP_0297 | JBx_249891 |
| pIY1605 | pRSF1010-Gm-NagR-pNagAa-dCas9-thrLABC-PP_0987 | JBx_249893 |
| pIY1606 | pRSF1010-Gm-NagR-pNagAa-dCas9-thrLABC-PP_2930 | JBx_249895 |
| pIY1607 | pRSF1010-Gm-NagR-pNagAa-dCas9-thrLABC-PP_3144 | JBx_249897 |
| pIY1608 | pRSF1010-Gm-NagR-pNagAa-dCas9-thrLABC-PP_3191 | JBx_249899 |
| pIY1609 | pRSF1010-Gm-NagR-pNagAa-dCas9-thrLABC-PP_3446 | JBx_249901 |
| pIY1610 | pRSF1010-Gm-NagR-pNagAa-dCas9-thrLABC-PP_5149 | JBx_249903 |
| pIY1611 | pRSF1010-Gm-NagR-pNagAa-dCas9-thrLABC-PP_0731 | JBx_249905 |
| pIY1612 | pRSF1010-Gm-NagR-pNagAa-dCas9-thrLABC-PP_3664 | JBx_249907 |
| pIY1613 | pRSF1010-Gm-NagR-pNagAa-dCas9-thrLABC-PP_4677 | JBx_249909 |
| pIY1614 | pRSF1010-Gm-NagR-pNagAa-dCas9-thrLABC-PP_0082 | JBx_249911 |
| pIY1615 | pRSF1010-Gm-NagR-pNagAa-dCas9-thrLABC-PP_0083 | JBx_249913 |
| pIY1616 | pRSF1010-Gm-NagR-pNagAa-dCas9-thrLABC-PP_0988 | JBx_249915 |
| pIY1617 | pRSF1010-Gm-NagR-pNagAa-dCas9-thrLABC-PP_5192 | JBx_249917 |
| pIY1659 | pRSF1010-Gm-NagR-pNagAa-dCas9-thrLABC-PP_0321 | JBx_249919 |
| pIY1660 | pRSF1010-Gm-NagR-pNagAa-dCas9-thrLABC-PP_1808 | JBx_249921 |
| pIY1661 | pRSF1010-Gm-NagR-pNagAa-dCas9-thrLABC-PP_4701 | JBx_249923 |
| pIY1662 | pRSF1010-Gm-NagR-pNagAa-dCas9-thrLABC-PP_1011 | JBx_249925 |
| pIY1663 | pRSF1010-Gm-NagR-pNagAa-dCas9-thrLABC-PP_1156 | JBx_249927 |
| pIY1664 | pRSF1010-Gm-NagR-pNagAa-dCas9-thrLABC-PP_2149 | JBx_249929 |
| pIY1665 | pRSF1010-Gm-NagR-pNagAa-dCas9-thrLABC-PP_4963 | JBx_249931 |
| pIY1666 | pRSF1010-Gm-NagR-pNagAa-dCas9-thrLABC-PP_5056 | JBx_249933 |
| pIY1667 | pRSF1010-Gm-NagR-pNagAa-dCas9-thrLABC-PP_1612 | JBx_249935 |

| | | |
|---|---|---|
| pIY1668 | pRSF1010-Gm-NagR-pNagAa-dCas9-thrLABC-PP_1362 | JBx_249937 |
| pIY1701 | pRSF1010-Gm-NagR-pNagAa-dCas9-thrLABC-PP_4188_0528 | JBx_249939 |
| pIY1702 | pRSF1010-Gm-NagR-pNagAa-dCas9-thrLABC-PP_4188_0812_4160_0168 | JBx_249941 |
| pIY1703 | pRSF1010-Gm-NagR-pNagAa-dCas9-thrLABC-PP_4188_0812_4160_0168_0528 | JBx_249943 |
| pIY1704 | pRSF1010-Gm-NagR-pNagAa-dCas9-thrLABC-PP_4188_0812_4160_0528 | JBx_249945 |
| pIY1705 | pRSF1010-Gm-NagR-pNagAa-dCas9-thrLABC-PP_4188_0812_4160_0528_4174 | JBx_249947 |
| pIY1706 | pRSF1010-Gm-NagR-pNagAa-dCas9-thrLABC-PP_4188_0812_4160_3280 | JBx_249949 |
| pIY1707 | pRSF1010-Gm-NagR-pNagAa-dCas9-thrLABC-PP_4188_0812_4160_4043 | JBx_249951 |
| pIY1708 | pRSF1010-Gm-NagR-pNagAa-dCas9-thrLABC-PP_4188_0813 | JBx_249953 |
| pIY1709 | pRSF1010-Gm-NagR-pNagAa-dCas9-thrLABC-PP_4188_0815 | JBx_249955 |
| pIY1710 | pRSF1010-Gm-NagR-pNagAa-dCas9-thrLABC-PP_4188_4160 | JBx_249957 |
| pIY1711 | pRSF1010-Gm-NagR-pNagAa-dCas9-thrLABC-PP_4188_4174 | JBx_249959 |
| pIY1712 | pRSF1010-Gm-NagR-pNagAa-dCas9-thrLABC-PP_4188_4266 | JBx_249961 |
| pIY1713 | pRSF1010-Gm-NagR-pNagAa-dCas9-thrLABC-PP_4192_0812 | JBx_249963 |
| pIY1714 | pRSF1010-Gm-NagR-pNagAa-dCas9-thrLABC-PP_4192_0812_4160_0168 | JBx_249965 |
| pIY1715 | pRSF1010-Gm-NagR-pNagAa-dCas9-thrLABC-PP_4192_4160 | JBx_249967 |
| pIY1716 | pRSF1010-Gm-NagR-pNagAa-dCas9-thrLABC-PP_4188_0528_4174 | JBx_249969 |
| pIY1717 | pRSF1010-Gm-NagR-pNagAa-dCas9-thrLABC-PP_4188_0812_4160_4174 | JBx_249971 |
| pIY1724 | pRSF1010-Gm-NagR-pNagAa-dCas9-thrLABC-PP_4188_0812 | JBx_249973 |
| pIY1725 | pRSF1010-Gm-NagR-pNagAa-dCas9-thrLABC-PP_4188_0812_4160 | JBx_249975 |
| pIY1726 | pRSF1010-Gm-NagR-pNagAa-dCas9-thrLABC-PP_4188_0528_4160 | JBx_249977 |
| pIY1727 | pRSF1010-Gm-NagR-pNagAa-dCas9-thrLABC-PP_4188_0812_4174 | JBx_249979 |
| pIY1728 | pRSF1010-Gm-NagR-pNagAa-dCas9-thrLABC-PP_4188_0528_4160_0168 | JBx_249981 |
| pIY1729 | pRSF1010-Gm-NagR-pNagAa-dCas9-thrLABC-PP_4188_0528_4160_3280 | JBx_249983 |
| pIY1730 | pRSF1010-Gm-NagR-pNagAa-dCas9-thrLABC-PP_4188_0528_4160_4043 | JBx_249985 |
| pIY1833 | pRSF1010-Gm-NagR-pNagAa-dCas9-thrLABC-PP_4011_4012 | JBx_249987 |
| pIY1834 | pRSF1010-Gm-NagR-pNagAa-dCas9-thrLABC-PP_4188_4116 | JBx_249989 |
| pIY1835 | pRSF1010-Gm-NagR-pNagAa-dCas9-thrLABC-PP_4188_0356 | JBx_249991 |
| pIY1836 | pRSF1010-Gm-NagR-pNagAa-dCas9-thrLABC-PP_4188_4116_0356 | JBx_249993 |
| pIY1837 | pRSF1010-Gm-NagR-pNagAa-dCas9-thrLABC-PP_4116_0356 | JBx_249995 |
| pIY1838 | pRSF1010-Gm-NagR-pNagAa-dCas9-thrLABC-PP_4188_0356_4160 | JBx_249997 |
| pIY1839 | pRSF1010-Gm-NagR-pNagAa-dCas9-thrLABC-PP_4188_1457 | JBx_249999 |
| pIY1840 | pRSF1010-Gm-NagR-pNagAa-dCas9-thrLABC-PP_4188_4635 | JBx_250001 |
| pIY1841 | pRSF1010-Gm-NagR-pNagAa-dCas9-thrLABC-PP_4188_1457_4635 | JBx_250003 |
| pIY1842 | pRSF1010-Gm-NagR-pNagAa-dCas9-thrLABC-PP_4188_1467_4160_0168 | JBx_250005 |
| pIY1843 | pRSF1010-Gm-NagR-pNagAa-dCas9-thrLABC-PP_4192_1457 | JBx_250007 |
| pIY1844 | pRSF1010-Gm-NagR-pNagAa-dCas9-thrLABC-PP_4192_4635 | JBx_250009 |
| pIY1845 | pRSF1010-Gm-NagR-pNagAa-dCas9-thrLABC-PP_4192_1457_4635 | JBx_250011 |
| pIY1846 | pRSF1010-Gm-NagR-pNagAa-dCas9-thrLABC-PP_4192_1457_4160_0168 | JBx_250013 |
| pIY1847 | pRSF1010-Gm-NagR-pNagAa-dCas9-thrLABC-PP_4011_4012_4116 | JBx_250015 |
| pIY1848 | pRSF1010-Gm-NagR-pNagAa-dCas9-thrLABC-PP_5075 | JBx_250017 |
| pIY1849 | pRSF1010-Gm-NagR-pNagAa-dCas9-thrLABC-PP_5076 | JBx_250019 |
| pIY1850 | pRSF1010-Gm-NagR-pNagAa-dCas9-thrLABC-PP_0675 | JBx_250021 |

| | | |
|---|---|---|
| pIY1851 | pRSF1010-Gm-NagR-pNagAa-dCas9-thrLABC-PP_2080 | JBx_250023 |
| pIY1852 | pRSF1010-Gm-NagR-pNagAa-dCas9-thrLABC-PP_5046_NT2 | JBx_250025 |
| pIY1853 | pRSF1010-Gm-NagR-pNagAa-dCas9-thrLABC-PP_3591 | JBx_250027 |
| pIY1854 | pRSF1010-Gm-NagR-pNagAa-dCas9-thrLABC-PP_2336 | JBx_250029 |
| pIY1855 | pRSF1010-Gm-NagR-pNagAa-dCas9-thrLABC-PP_2652 | JBx_250031 |
| pIY1856 | pRSF1010-Gm-NagR-pNagAa-dCas9-thrLABC-PP_3662 | JBx_250033 |
| pIY1857 | pRSF1010-Gm-NagR-pNagAa-dCas9-thrLABC-PP_4012_NT2 | JBx_250035 |
| pIY1858 | pRSF1010-Gm-NagR-pNagAa-dCas9-thrLABC-PP_0356 | JBx_250037 |
| pIY1501 | pRSF1010-Gm-NagR-pNagAa-dCas9-thrLABC-PP_1610_4188 | JBx_250612 |
| pIY1502 | pRSF1010-Gm-NagR-pNagAa-dCas9-thrLABC-PP_0654_4266 | JBx_250614 |
| pIY1503 | pRSF1010-Gm-NagR-pNagAa-dCas9-thrLABC-PP_4365_4679 | JBx_250616 |
| pIY1504 | pRSF1010-Gm-NagR-pNagAa-dCas9-thrLABC-PP_4185_4679 | JBx_250618 |
| pIY1505 | pRSF1010-Gm-NagR-pNagAa-dCas9-thrLABC-PP_0944_4185 | JBx_250620 |
| pIY1506 | pRSF1010-Gm-NagR-pNagAa-dCas9-thrLABC-PP_0104_0813 | JBx_250622 |
| pIY1507 | pRSF1010-Gm-NagR-pNagAa-dCas9-thrLABC-PP_0944_1755 | JBx_250624 |
| pIY1509 | pRSF1010-Gm-NagR-pNagAa-dCas9-thrLABC-PP_1755_4189 | JBx_250626 |
| pIY1510 | pRSF1010-Gm-NagR-pNagAa-dCas9-thrLABC-PP_1023_5419 | JBx_250628 |
| pIY1511 | pRSF1010-Gm-NagR-pNagAa-dCas9-thrLABC-PP_4189_5186 | JBx_250630 |
| pIY1512 | pRSF1010-Gm-NagR-pNagAa-dCas9-thrLABC-PP_0105_0813 | JBx_250632 |
| pIY1513 | pRSF1010-Gm-NagR-pNagAa-dCas9-thrLABC-PP_0814_4189 | JBx_250634 |
| pIY1514 | pRSF1010-Gm-NagR-pNagAa-dCas9-thrLABC-PP_4188_4679 | JBx_250636 |
| pIY1515 | pRSF1010-Gm-NagR-pNagAa-dCas9-thrLABC-PP_4043_4188 | JBx_250638 |
| pIY1517 | pRSF1010-Gm-NagR-pNagAa-dCas9-thrLABC-PP_0944_1610_1755 | JBx_250640 |
| pIY1518 | pRSF1010-Gm-NagR-pNagAa-dCas9-thrLABC-PP_0813_0814_5186 | JBx_250642 |
| pIY1520 | pRSF1010-Gm-NagR-pNagAa-dCas9-thrLABC-PP_1755_4189_4266 | JBx_250644 |
| pIY1522 | pRSF1010-Gm-NagR-pNagAa-dCas9-thrLABC-PP_0944_1610_4189 | JBx_250646 |
| pIY1523 | pRSF1010-Gm-NagR-pNagAa-dCas9-thrLABC-PP_0225_0813_4192 | JBx_250648 |
| pIY1524 | pRSF1010-Gm-NagR-pNagAa-dCas9-thrLABC-PP_0105_0277_0813 | JBx_250650 |
| pIY1525 | pRSF1010-Gm-NagR-pNagAa-dCas9-thrLABC-PP_0813_0814_1610 | JBx_250652 |
| pIY1526 | pRSF1010-Gm-NagR-pNagAa-dCas9-thrLABC-PP_4189_4266 | JBx_250654 |
| pIY1527 | pRSF1010-Gm-NagR-pNagAa-dCas9-thrLABC-PP_0104_4679 | JBx_250656 |
| pIY1528 | pRSF1010-Gm-NagR-pNagAa-dCas9-thrLABC-PP_0105_4189 | JBx_250658 |
| pIY1529 | pRSF1010-Gm-NagR-pNagAa-dCas9-thrLABC-PP_0944_5186 | JBx_250660 |
| pIY1530 | pRSF1010-Gm-NagR-pNagAa-dCas9-thrLABC-PP_0105_4185 | JBx_250662 |
| pIY1531 | pRSF1010-Gm-NagR-pNagAa-dCas9-thrLABC-PP_4189_5186 | JBx_250664 |
| pIY1532 | pRSF1010-Gm-NagR-pNagAa-dCas9-thrLABC-PP_0105_4188 | JBx_250666 |
| pIY1533 | pRSF1010-Gm-NagR-pNagAa-dCas9-thrLABC-PP_1023_4188 | JBx_250668 |
| pIY1534 | pRSF1010-Gm-NagR-pNagAa-dCas9-thrLABC-PP_0104_4188 | JBx_250670 |
| pIY1535 | pRSF1010-Gm-NagR-pNagAa-dCas9-thrLABC-PP_0225_4185 | JBx_250672 |
| pIY1536 | pRSF1010-Gm-NagR-pNagAa-dCas9-thrLABC-PP_0104_4185_4188 | JBx_250674 |
| pIY1537 | pRSF1010-Gm-NagR-pNagAa-dCas9-thrLABC-PP_0105_0813_4188 | JBx_250676 |
| pIY1538 | pRSF1010-Gm-NagR-pNagAa-dCas9-thrLABC-PP_0654_4185_5186 | JBx_250678 |
| pIY1544 | pRSF1010-Gm-NagR-pNagAa-dCas9-thrLABC-PP_0225_0944_4189 | JBx_250680 |

| | | |
|---|---|---|
| pIY1545 | pRSF1010-Gm-NagR-pNagAa-dCas9-thrLABC-PP_0654_1023_4189 | JBx_250682 |
| pIY1546 | pRSF1010-Gm-NagR-pNagAa-dCas9-thrLABC-PP_1610_4188_4188 | JBx_250684 |
| pIY1547 | pRSF1010-Gm-NagR-pNagAa-dCas9-thrLABC-PP_0654_4185_4188 | JBx_250686 |
| pIY1548 | pRSF1010-Gm-NagR-pNagAa-dCas9-thrLABC-PP_4188_4635_5419 | JBx_250688 |

*Link to the plasmid maps of some constructs are provided.

### Supplementary Table S6. List of oligos used in this study

| Oligo name | Sequence (5' to 3') |
|---|---|
| IY677_RFP_NT1_sgRNA_F | TCTGGGTCTCTTAGCatttcgaactcgtgaccgttaaGTTTGGAGACCATCG |
| IY678_RFP_NT1_sgRNA_R | CGATGGTCTCCAAACttaacggtcacgagttcgaaatGCTAAGAGACCCAGA |
| IY679_gltA_NT1_sgRNA_F | TCTGGGTCTCTTAGCGTTAAAATGGGCAGCTCGACGGGTTTGGAGACCATCG |
| IY680_gltA_NT1_sgRNA_R | CGATGGTCTCCAAACCCGTCGAGCTGCCCATTTTAACGCTAAGAGACCCAGA |
| IY681_edd_NT1_sgRNA_F | TCTGGGTCTCTTAGCgctgggtgacctcaaggatgcgGTTTGGAGACCATCG |
| IY682_edd_NT1_sgRNA_R | CGATGGTCTCCAAACcgcatccttgaggtcacccagcGCTAAGAGACCCAGA |
| IY683_nontarget_NT1_sgRNA_F | TCTGGGTCTCTTAGCtcctgcacatagccttccggcaGTTTGGAGACCATCG |
| IY684_nontarget_NT1_sgRNA_R | CGATGGTCTCCAAACtgccggaaggctatgtgcaggaGCTAAGAGACCCAGA |
| IY695_PP4549_NT1_sgRNA_F | TCTGGGTCTCTTAGCcaggcgcttgagtggtggcagcGTTTGGAGACCATCG |
| IY696_PP4549_NT1_sgRNA_R | CGATGGTCTCCAAACgctgccaccactcaagcgcctgGCTAAGAGACCCAGA |
| IY697_PP4550_NT1_sgRNA_F | TCTGGGTCTCTTAGCCCGCCTGGGGCAACCGGTAGGCGTTTGGAGACCATCG |
| IY698_PP4550_NT1_sgRNA_R | CGATGGTCTCCAAACGCCTACCGGTTGCCCCAGGCGGGCTAAGAGACCCAGA |
| IY699_PP0368_NT1_sgRNA_F | TCTGGGTCTCTTAGCTCACCGGCCGAAATGAAGATCTGTTTGGAGACCATCG |
| IY700_PP0368_NT1_sgRNA_R | CGATGGTCTCCAAACAGATCTTCATTTCGGCCGGTGAGCTAAGAGACCCAGA |
| IY701_PP0370_NT1_sgRNA_F | TCTGGGTCTCTTAGCAACGCCGGCATACATGTTCGGAGTTTGGAGACCATCG |
| IY702_PP0370_NT1_sgRNA_R | CGATGGTCTCCAAACTCCGAACATGTATGCCGGCGTTGCTAAGAGACCCAGA |
| IY703_PP2136_NT1_sgRNA_F | TCTGGGTCTCTTAGCaggctctcggcgacagaggtctGTTTGGAGACCATCG |
| IY704_PP2136_NT1_sgRNA_R | CGATGGTCTCCAAACagacctctgtcgccgagagcctGCTAAGAGACCCAGA |
| IY705_hsdR_NT1_sgRNA_F | TCTGGGTCTCTTAGCcctaggactgagctagccgtcaGTTTGGAGACCATCG |
| IY706_hsdR_NT1_sgRNA_R | CGATGGTCTCCAAACtgacggctagctcagtcctaggGCTAAGAGACCCAGA |
| IY707_hsdR_NT2_sgRNA_F | TCTGGGTCTCTTAGCGCCGTAGAGGATGAAGTCTGCAGTTTGGAGACCATCG |
| IY708_hsdR_NT2_sgRNA_R | CGATGGTCTCCAAACTGCAGACTTCATCCTCTACGGCGCTAAGAGACCCAGA |
| IY709_hsdR_NT3_sgRNA_F | TCTGGGTCTCTTAGCATGTCCCACCCAGCGTGCTGGAGTTTGGAGACCATCG |
| IY710_hsdR_NT3_sgRNA_R | CGATGGTCTCCAAACTCCAGCACGCTGGGTGGGACATGCTAAGAGACCCAGA |
| IY711_pta_NT1_sgRNA_F | TCTGGGTCTCTTAGCggtggcttgatgccgtgagtgcGTTTGGAGACCATCG |
| IY712_pta_NT1_sgRNA_R | CGATGGTCTCCAAACgcactcacggcatcaagccaccGCTAAGAGACCCAGA |
| IY713_bioB_NT1_sgRNA_F | TCTGGGTCTCTTAGCCCTGGAACAGCAAGTCATTGAAGTTTGGAGACCATCG |
| IY714_bioB_NT1_sgRNA_R | CGATGGTCTCCAAACTTCAATGACTTGCTGTTCCAGGGCTAAGAGACCCAGA |
| IY715_birA_NT1_sgRNA_F | TCTGGGTCTCTTAGCactgcgctgcgactcaccccctaGTTTGGAGACCATCG |
| IY716_birA_NT1_sgRNA_R | CGATGGTCTCCAAACtaggggtgagtcgcagcgcagtGCTAAGAGACCCAGA |
| IY717_PP2589_NT2_sgRNA_F | TCTGGGTCTCTTAGCgccgaaaacggtgattgctcggGTTTGGAGACCATCG |
| IY718_PP2589_NT2_sgRNA_R | CGATGGTCTCCAAACccgagcaatcaccgttttcggcGCTAAGAGACCCAGA |
| IY719_glgC_NT1_sgRNA_F | TCTGGGTCTCTTAGCgcgaactgaacaggtgcatcaaGTTTGGAGACCATCG |
| IY720_glgC_NT1_sgRNA_R | CGATGGTCTCCAAACttgatgcacctgttcagttcgcGCTAAGAGACCCAGA |
| IY721_gltA_NT2_sgRNA_F | TCTGGGTCTCTTAGCTGTAAACCATCGCAGCCAGGGTGTTTGGAGACCATCG |
| IY722_gltA_NT2_sgRNA_R | CGATGGTCTCCAAACACCCTGGCTGCGATGGTTTACAGCTAAGAGACCCAGA |
| IY723_gltA_NT3_sgRNA_F | TCTGGGTCTCTTAGCGTACGTGCCAGGGCGAAGATCAGTTTGGAGACCATCG |
| IY724_gltA_NT3_sgRNA_R | CGATGGTCTCCAAACTGATCTTCGCCCTGGCACGTACGCTAAGAGACCCAGA |
| IY725_accA_NT1_sgRNA_F | TCTGGGTCTCTTAGCggtgttgctcttgtcttgcagaGTTTGGAGACCATCG |
| IY726_accA_NT1_sgRNA_R | CGATGGTCTCCAAACtctgcaagacaagagcaacaccGCTAAGAGACCCAGA |

| | |
|---|---|
| IY727_accA_NT2_sgRNA_F | TCTGGGTCTCTTAGCgttccaggcgatggcttcactcGTTTGGAGACCATCG |
| IY728_accA_NT2_sgRNA_R | CGATGGTCTCCAAACgagtgaagccatcgcctggaacGCTAAGAGACCCAGA |
| IY729_accA_NT3_sgRNA_F | TCTGGGTCTCTTAGCacagaatcgaagcacagccttcGTTTGGAGACCATCG |
| IY730_accA_NT3_sgRNA_R | CGATGGTCTCCAAACgaaggctgtgcttcgattctgtGCTAAGAGACCCAGA |
| IY731_accA_NT4_sgRNA_F | TCTGGGTCTCTTAGCaccagatcggcacggatgctttGTTTGGAGACCATCG |
| IY732_accA_NT4_sgRNA_R | CGATGGTCTCCAAACaaagcatccgtgccgatctggtGCTAAGAGACCCAGA |
| IY735_PP1444_NT1_sgRNA_F | TCTGGGTCTCTTAGCGTAGTACAGCGATCCGCCCAGCGTTTGGAGACCATCG |
| IY736_PP1444_NT1_sgRNA_R | CGATGGTCTCCAAACGCTGGGCGGATCGCTGTACTACGCTAAGAGACCCAGA |
| IY737_PP0339_NT1_sgRNA_F | TCTGGGTCTCTTAGCgtggtgatcgcatatggcagctGTTTGGAGACCATCG |
| IY738_PP0339_NT1_sgRNA_R | CGATGGTCTCCAAACagctgccatatgcgatcaccacGCTAAGAGACCCAGA |
| IY739_PP0338_NT1_sgRNA_F | TCTGGGTCTCTTAGCtgatcacgccagccttgggcgcGTTTGGAGACCATCG |
| IY740_PP0338_NT1_sgRNA_R | CGATGGTCTCCAAACgcgcccaaggctggcgtgatcaGCTAAGAGACCCAGA |
| IY747_PP3394_NT1_sgRNA_F | TCTGGGTCTCTTAGCCCAGCACGGTTACTTGCAGGTCGTTTGGAGACCATCG |
| IY748_PP3394_NT1_sgRNA_R | CGATGGTCTCCAAACGACCTGCAAGTAACCGTGCTGGGCTAAGAGACCCAGA |
| IY749_PP3122_NT1_sgRNA_F | TCTGGGTCTCTTAGCGCGCCGCTTGATTTCGCTGATAGTTTGGAGACCATCG |
| IY750_PP3122_NT1_sgRNA_R | CGATGGTCTCCAAACTATCAGCGAAATCAAGCGGCGCGCTAAGAGACCCAGA |
| IY751_PP3123_NT1_sgRNA_F | TCTGGGTCTCTTAGCGCGGGTACATAGTTGGCCACCAGTTTGGAGACCATCG |
| IY752_PP3123_NT1_sgRNA_R | CGATGGTCTCCAAACTGGTGGCCAACTATGTACCCGCGCTAAGAGACCCAGA |
| IY753_PP3071_NT1_sgRNA_F | TCTGGGTCTCTTAGCGCATTGGCGAACCACTGTGCATGTTTGGAGACCATCG |
| IY754_PP3071_NT1_sgRNA_R | CGATGGTCTCCAAACATGCACAGTGGTTCGCCAATGCGCTAAGAGACCCAGA |
| IY755_PP0582_NT1_sgRNA_F | TCTGGGTCTCTTAGCacccaagacgcactcgcgggtcGTTTGGAGACCATCG |
| IY756_PP0582_NT1_sgRNA_R | CGATGGTCTCCAAACgacccgcgagtgcgtcttgggtGCTAAGAGACCCAGA |
| IY757_PP2051_NT1_sgRNA_F | TCTGGGTCTCTTAGCtcgatgccagcacgttctactgGTTTGGAGACCATCG |
| IY758_PP2051_NT1_sgRNA_R | CGATGGTCTCCAAACcagtagaacgtgctggcatcgaGCTAAGAGACCCAGA |
| IY759_PP2215_NT1_sgRNA_F | TCTGGGTCTCTTAGCctgacgataacgatggggtcatGTTTGGAGACCATCG |
| IY760_PP2215_NT1_sgRNA_R | CGATGGTCTCCAAACatgaccccatcgttatcgtcagGCTAAGAGACCCAGA |
| IY761_PP3355_NT1_sgRNA_F | TCTGGGTCTCTTAGCacactgtctaccgcctgaggggGTTTGGAGACCATCG |
| IY762_PP3355_NT1_sgRNA_R | CGATGGTCTCCAAACcccctcaggcggtagacagtgtGCTAAGAGACCCAGA |
| IY763_PP3754_NT1_sgRNA_F | TCTGGGTCTCTTAGCgattaccgtgcccatcaccatgGTTTGGAGACCATCG |
| IY764_PP3754_NT1_sgRNA_R | CGATGGTCTCCAAACcatggtgatgggcacggtaatcGCTAAGAGACCCAGA |
| IY765_PP4636_NT1_sgRNA_F | TCTGGGTCTCTTAGCccgaggtcaaccgcaggcactgGTTTGGAGACCATCG |
| IY766_PP4636_NT1_sgRNA_R | CGATGGTCTCCAAACcagtgcctgcggttgacctcggGCTAAGAGACCCAGA |
| IY767_PP3282_NT1_sgRNA_F | TCTGGGTCTCTTAGCgggcggttatcgtagagcttgaGTTTGGAGACCATCG |
| IY768_PP3282_NT1_sgRNA_R | CGATGGTCTCCAAACtcaagctctacgataaccgcccGCTAAGAGACCCAGA |
| IY769_PP3755_NT1_sgRNA_F | TCTGGGTCTCTTAGCctgggtgtagtcggtgctggtaGTTTGGAGACCATCG |
| IY770_PP3755_NT1_sgRNA_R | CGATGGTCTCCAAACtaccagcaccgactacacccagGCTAAGAGACCCAGA |
| IY771_PP1157_NT1_sgRNA_F | TCTGGGTCTCTTAGCgaaacctgcggactgctcgtgaGTTTGGAGACCATCG |
| IY772_PP1157_NT1_sgRNA_R | CGATGGTCTCCAAACtcacgagcagtccgcaggtttcGCTAAGAGACCCAGA |
| IY773_PP1394_NT1_sgRNA_F | TCTGGGTCTCTTAGCtgacgcggggtgatgtggcgaaGTTTGGAGACCATCG |
| IY774_PP1394_NT1_sgRNA_R | CGATGGTCTCCAAACttcgccacatcaccccgcgtcaGCTAAGAGACCCAGA |
| IY775_PP3365_NT1_sgRNA_F | TCTGGGTCTCTTAGCgaaacctgcggactgctcgtgaGTTTGGAGACCATCG |
| IY776_PP3365_NT1_sgRNA_R | CGATGGTCTCCAAACtcacgagcagtccgcaggtttcGCTAAGAGACCCAGA |
| IY777_PP4679_NT1_sgRNA_F | TCTGGGTCTCTTAGCTCAGCGTCAGACGCGACAGGGTGTTTGGAGACCATCG |

| | |
|---|---|
| IY779_PP4679_NT1_sgRNA_R | CGATGGTCTCCAAACACCCTGTCGCGTCTGACGCTGAGCTAAGAGACCCAGA |
| IY780_PP4680_NT1_sgRNA_F | TCTGGGTCTCTTAGCAGACCGACGGTAACGTCTACACGTTTGGAGACCATCG |
| IY781_PP4680_NT1_sgRNA_R | CGATGGTCTCCAAACGTGTAGACGTTACCGTCGGTCTGCTAAGAGACCCAGA |
| IY782_PP1777_NT1_sgRNA_F | TCTGGGTCTCTTAGCagtaaggcccctcatcaatgctGTTTGGAGACCATCG |
| IY783_PP1777_NT1_sgRNA_R | CGATGGTCTCCAAACagcattgatgaggggccttactGCTAAGAGACCCAGA |
| IY784_PP3578_NT1_sgRNA_F | TCTGGGTCTCTTAGCgcgtctgggcggcccgtgtaatGTTTGGAGACCATCG |
| IY785_PP3578_NT1_sgRNA_R | CGATGGTCTCCAAACattacacgggccgcccagacgcGCTAAGAGACCCAGA |
| IY786_PP5288_NT1_sgRNA_F | TCTGGGTCTCTTAGCcacgccgcggatgtcatatgcgGTTTGGAGACCATCG |
| IY787_PP5288_NT1_sgRNA_R | CGATGGTCTCCAAACcgcatatgacatccgcggcgtgGCTAAGAGACCCAGA |
| IY788_PP0597_NT1_sgRNA_F | TCTGGGTCTCTTAGCccagtggacgggttgaagacatGTTTGGAGACCATCG |
| IY789_PP0597_NT1_sgRNA_R | CGATGGTCTCCAAACatgtcttcaacccgtccactggGCTAAGAGACCCAGA |
| IY790_PP4667_NT1_sgRNA_F | TCTGGGTCTCTTAGCttgacgatgtcacgccattcggGTTTGGAGACCATCG |
| IY791_PP4667_NT1_sgRNA_R | CGATGGTCTCCAAACccgaatggcgtgacatcgtcaaGCTAAGAGACCCAGA |
| IY792_PP2137_NT1_sgRNA_F | TCTGGGTCTCTTAGCgcggtgcatgccacccttggagGTTTGGAGACCATCG |
| IY793_PP2137_NT1_sgRNA_R | CGATGGTCTCCAAACctccaagggtggcatgcaccgcGCTAAGAGACCCAGA |
| IY794_PP3280_NT1_sgRNA_F | TCTGGGTCTCTTAGCgggccccggcatagcggccaatGTTTGGAGACCATCG |
| IY795_PP3280_NT1_sgRNA_R | CGATGGTCTCCAAACattggccgctatgccggggcccGCTAAGAGACCCAGA |
| IY796_PP2471_NT1_sgRNA_F | TCTGGGTCTCTTAGCcgaaggtcgaagttgccgaaacGTTTGGAGACCATCG |
| IY797_PP2471_NT1_sgRNA_R | CGATGGTCTCCAAACgtttcggcaacttcgaccttcgGCTAAGAGACCCAGA |
| IY798_PP4737_NT1_sgRNA_F | TCTGGGTCTCTTAGCcggggccgatcttgcgcccgaaGTTTGGAGACCATCG |
| IY799_PP4737_NT1_sgRNA_R | CGATGGTCTCCAAACttcgggcgcaagatcggccccgGCTAAGAGACCCAGA |
| IY800_PP4894_NT1_sgRNA_F | TCTGGGTCTCTTAGCcgttgaccaggtaaatggatacGTTTGGAGACCATCG |
| IY801_PP4894_NT1_sgRNA_R | CGATGGTCTCCAAACgtatccatttacctggtcaacgGCTAAGAGACCCAGA |
| IY802_PP2213_NT1_sgRNA_F | TCTGGGTCTCTTAGCcctcggcgtaatcgcgcatcatGTTTGGAGACCATCG |
| IY803_PP2213_NT1_sgRNA_R | CGATGGTCTCCAAACatgatgcgcgattacgccgaggGCTAAGAGACCCAGA |
| IY804_PP4487_NT1_sgRNA_F | TCTGGGTCTCTTAGCcctcgggacgaacgggatacagGTTTGGAGACCATCG |
| IY805_PP4487_NT1_sgRNA_R | CGATGGTCTCCAAACctgtatcccgttcgtcccgaggGCTAAGAGACCCAGA |
| IY806_PP4702_NT1_sgRNA_F | TCTGGGTCTCTTAGCttcgaccgactggcggtagaggGTTTGGAGACCATCG |
| IY807_PP4702_NT1_sgRNA_R | CGATGGTCTCCAAACcctctaccgccagtcggtcgaaGCTAAGAGACCCAGA |
| IY808_PP0154_NT1_sgRNA_F | TCTGGGTCTCTTAGCtggcacgttcggccaatgcatgGTTTGGAGACCATCG |
| IY809_PP0154_NT1_sgRNA_R | CGATGGTCTCCAAACcatgcattggccgaacgtgccaGCTAAGAGACCCAGA |
| IY816_PP0528_NT1_sgRNA_F | TCTGGGTCTCTTAGCGCTGTAGCGCATGGCGGCGTAAGTTTGGAGACCATCG |
| IY817_PP0528_NT1_sgRNA_R | CGATGGTCTCCAAACTTACGCCGCCATGCGCTACAGCGCTAAGAGACCCAGA |
| IY818_PP0548_NT1_sgRNA_F | TCTGGGTCTCTTAGCtcgagcaggcgaaggccatactGTTTGGAGACCATCG |
| IY819_PP0548_NT1_sgRNA_R | CGATGGTCTCCAAACagtatggccttcgcctgctcgaGCTAAGAGACCCAGA |
| IY820_PP0527_NT1_sgRNA_F | TCTGGGTCTCTTAGCggcgcccggtgaggatcttgtgGTTTGGAGACCATCG |
| IY821_PP0527_NT1_sgRNA_R | CGATGGTCTCCAAACcacaagatcctcaccgggcgccGCTAAGAGACCCAGA |
| IY822_PP2533_NT1_sgRNA_F | TCTGGGTCTCTTAGCcaagttgttccagcgtggcaggGTTTGGAGACCATCG |
| IY823_PP2533_NT1_sgRNA_R | CGATGGTCTCCAAACcctgccacgctggaacaacttgGCTAAGAGACCCAGA |
| IY824_PP5155_NT1_sgRNA_F | TCTGGGTCTCTTAGCttctcagccagtaccgaaagctGTTTGGAGACCATCG |
| IY825_PP5155_NT1_sgRNA_R | CGATGGTCTCCAAACagctttcggtactggctgagaaGCTAAGAGACCCAGA |
| IY826_PP1444_NT2_sgRNA_F | TCTGGGTCTCTTAGCTGGCGAACCAGATAGCCAGCCGGTTTGGAGACCATCG |
| IY827_PP1444_NT2_sgRNA_R | CGATGGTCTCCAAACCGGCTGGCTATCTGGTTCGCCAGCTAAGAGACCCAGA |

| | |
|---|---|
| IY828_PP0339_NT2_sgRNA_F | TCTGGGTCTCTTAGCcgtgcttcgtgggtgacagggaGTTTGGAGACCATCG |
| IY829_PP0339_NT2_sgRNA_R | CGATGGTCTCCAAACtccctgtcacccacgaagcacgGCTAAGAGACCCAGA |
| IY830_PP2051_NT2_sgRNA_F | TCTGGGTCTCTTAGCtgctcgctcatgcgcaataccgGTTTGGAGACCATCG |
| IY831_PP2051_NT2_sgRNA_R | CGATGGTCTCCAAACcggtattgcgcatgagcgagcaGCTAAGAGACCCAGA |
| IY832_PP2215_NT2_sgRNA_F | TCTGGGTCTCTTAGCccttgccctcggtgacttccacGTTTGGAGACCATCG |
| IY833_PP2215_NT2_sgRNA_R | CGATGGTCTCCAAACgtggaagtcaccgagggcaaggGCTAAGAGACCCAGA |
| IY834_PP1866_NT1_sgRNA_F | TCTGGGTCTCTTAGCccatacgccccactttcaccacGTTTGGAGACCATCG |
| IY835_PP1866_NT1_sgRNA_R | CGATGGTCTCCAAACgtggtgaaagtggggcgtatggGCTAAGAGACCCAGA |
| IY836_PP2324_NT1_sgRNA_F | TCTGGGTCTCTTAGCccttgatgtcgtggatggagcaGTTTGGAGACCATCG |
| IY837_PP2324_NT1_sgRNA_R | CGATGGTCTCCAAACtgctccatccacgacatcaaggGCTAAGAGACCCAGA |
| IY838_PP3080_NT1_sgRNA_F | TCTGGGTCTCTTAGCcggcgtattccagggccgagcgGTTTGGAGACCATCG |
| IY839_PP3080_NT1_sgRNA_R | CGATGGTCTCCAAACcgctcggccctggaatacgccgGCTAAGAGACCCAGA |
| IY840_PP5351_NT1_sgRNA_F | TCTGGGTCTCTTAGCatcgggaagtagtcgacccgtgGTTTGGAGACCATCG |
| IY841_PP5351_NT1_sgRNA_R | CGATGGTCTCCAAACcacgggtcgactacttcccgatGCTAAGAGACCCAGA |
| IY842_PP4736_NT1_sgRNA_F | TCTGGGTCTCTTAGCcgacacagtcgacatggtgaacGTTTGGAGACCATCG |
| IY843_PP4736_NT1_sgRNA_R | CGATGGTCTCCAAACgttcaccatgtcgactgtgtcgGCTAAGAGACCCAGA |
| IY844_PP1773_NT1_sgRNA_F | TCTGGGTCTCTTAGCttcgggttacggcctacgcgagGTTTGGAGACCATCG |
| IY845_PP1773_NT1_sgRNA_R | CGATGGTCTCCAAACctcgcgtaggccgtaacccgaaGCTAAGAGACCCAGA |
| IY846_PP0356_NT1_sgRNA_F | TCTGGGTCTCTTAGCtcttcgtccacgttctgggtggGTTTGGAGACCATCG |
| IY847_PP0356_NT1_sgRNA_R | CGATGGTCTCCAAACccacccagaacgtggacgaagaGCTAAGAGACCCAGA |
| IY848_PP4379_NT1_sgRNA_F | TCTGGGTCTCTTAGCacgatcttcgaatacgggtcggGTTTGGAGACCATCG |
| IY849_PP4379_NT1_sgRNA_R | CGATGGTCTCCAAACccgacccgtattcgaagatcgtGCTAAGAGACCCAGA |
| IY850_PP1319_NT1_sgRNA_F | TCTGGGTCTCTTAGCcccaggctgtgaccggaaacgaGTTTGGAGACCATCG |
| IY851_PP1319_NT1_sgRNA_R | CGATGGTCTCCAAACtcgtttccggtcacagcctgggGCTAAGAGACCCAGA |
| IY852_PP5258_NT1_sgRNA_F | TCTGGGTCTCTTAGCgtgccaggtttcacgcatgtggGTTTGGAGACCATCG |
| IY853_PP5258_NT1_sgRNA_R | CGATGGTCTCCAAACccacatgcgtgaaacctggcacGCTAAGAGACCCAGA |
| IY854_PP1616_NT1_sgRNA_F | TCTGGGTCTCTTAGCtggcccttgtacgagaagcgcgGTTTGGAGACCATCG |
| IY855_PP1616_NT1_sgRNA_R | CGATGGTCTCCAAACcgcgcttctcgtacaagggccaGCTAAGAGACCCAGA |
| IY856_PP2803_NT1_sgRNA_F | TCTGGGTCTCTTAGCcccgtgccccgatagcgctgggGTTTGGAGACCATCG |
| IY857_PP2803_NT1_sgRNA_R | CGATGGTCTCCAAACcccagcgctatcggggcacgggGCTAAGAGACCCAGA |
| IY877_PP5420_sgRNA_NT1_F | TCTGGGTCTCTTAGCCCTTTCCACTGCCACAACACCAGTTTGGAGACCATCG |
| IY878_PP5420_sgRNA_NT1_R | CGATGGTCTCCAAACTGGTGTTGTGGCAGTGGAAAGGGCTAAGAGACCCAGA |
| IY879_PP5419_sgRNA_NT1_F | TCTGGGTCTCTTAGCGCCTTGGCTTCTGCAGCCGAATGTTTGGAGACCATCG |
| IY880_PP5419_sgRNA_NT1_R | CGATGGTCTCCAAACATTCGGCTGCAGAAGCCAAGGCGCTAAGAGACCCAGA |
| IY881_PP5417_sgRNA_NT1_F | TCTGGGTCTCTTAGCAGTACAAAAATCAGGAAAGCAAGTTTGGAGACCATCG |
| IY882_PP5417_sgRNA_NT1_R | CGATGGTCTCCAAACTTGCTTTCCTGATTTTTGTACTGCTAAGAGACCCAGA |
| IY883_PP5416_sgRNA_NT1_F | TCTGGGTCTCTTAGCCCGTGGATCCGCACGATACCGTGTTTGGAGACCATCG |
| IY884_PP5416_sgRNA_NT1_R | CGATGGTCTCCAAACACGGTATCGTGCGGATCCACGGGCTAAGAGACCCAGA |
| IY885_PP5413_sgRNA_NT1_F | TCTGGGTCTCTTAGCGCCTTACCAACTGGAACGGAAAGTTTGGAGACCATCG |
| IY886_PP5413_sgRNA_NT1_R | CGATGGTCTCCAAACTTTCCGTTCCAGTTGGTAAGGCGCTAAGAGACCCAGA |
| IY887_PP0806_sgRNA_NT1_F | TCTGGGTCTCTTAGCGGTGAACGCTGCCGCATCGTACGTTTGGAGACCATCG |
| IY888_PP0806_sgRNA_NT1_R | CGATGGTCTCCAAACGTACGATGCGGCAGCGTTCACCGCTAAGAGACCCAGA |
| IY889_PP0168_sgRNA_NT1_F | TCTGGGTCTCTTAGCTATCGGTCACTTGGGTAACCGCGTTTGGAGACCATCG |

| | |
|---|---|
| IY890_PP0168_sgRNA_NT1_R | CGATGGTCTCCAAACGCGGTTACCCAAGTGACCGATAGCTAAGAGACCCAGA |
| IY891_PP5336_sgRNA_NT1_F | TCTGGGTCTCTTAGCCTGGAACAGCAGATCCGGGGTGGTTTGGAGACCATCG |
| IY892_PP5336_sgRNA_NT1_R | CGATGGTCTCCAAACCACCCCGGATCTGCTGTTCCAGGCTAAGAGACCCAGA |
| IY893_PP5335_sgRNA_NT1_F | TCTGGGTCTCTTAGCTGCGAGAGGAAGGCCACGGTCTGTTTGGAGACCATCG |
| IY894_PP5335_sgRNA_NT1_R | CGATGGTCTCCAAACAGACCGTGGCCTTCCTCTCGCAGCTAAGAGACCCAGA |
| IY895_PP4016_sgRNA_NT1_F | TCTGGGTCTCTTAGCtcaggccgaattcgctgaaaatGTTTGGAGACCATCG |
| IY896_PP4016_sgRNA_NT1_R | CGATGGTCTCCAAACattttcagcgaattcggcctgaGCTAAGAGACCCAGA |
| IY897_PP4119_sgRNA_NT1_F | TCTGGGTCTCTTAGCCGGGCAGCATGCCGGATTCGAAGTTTGGAGACCATCG |
| IY898_PP4119_sgRNA_NT1_R | CGATGGTCTCCAAACTTCGAATCCGGCATGCTGCCCGGCTAAGAGACCCAGA |
| IY899_PP4120_sgRNA_NT1_F | TCTGGGTCTCTTAGCTGACCATGAAGTCGGCCTGACGGTTTGGAGACCATCG |
| IY900_PP4120_sgRNA_NT1_R | CGATGGTCTCCAAACCGTCAGGCCGACTTCATGGTCAGCTAAGAGACCCAGA |
| IY901_PP4121_sgRNA_NT1_F | TCTGGGTCTCTTAGCGAAGCTGAGCACCTCTTTGAGCGTTTGGAGACCATCG |
| IY902_PP4121_sgRNA_NT1_R | CGATGGTCTCCAAACGCTCAAAGAGGTGCTCAGCTTCGCTAAGAGACCCAGA |
| IY903_PP4122_sgRNA_NT1_F | TCTGGGTCTCTTAGCGGGCTTCGATGGACGCCGCGCGGTTTGGAGACCATCG |
| IY904_PP4122_sgRNA_NT1_R | CGATGGTCTCCAAACCGCGCGGCGTCCATCGAAGCCCGCTAAGAGACCCAGA |
| IY905_PP4123_sgRNA_NT1_F | TCTGGGTCTCTTAGCATCAGGCCCCACTTCACGCCAGGTTTGGAGACCATCG |
| IY906_PP4123_sgRNA_NT1_R | CGATGGTCTCCAAACCTGGCGTGAAGTGGGGCCTGATGCTAAGAGACCCAGA |
| IY907_PP4124_sgRNA_NT1_F | TCTGGGTCTCTTAGCTCTTCATCGTCGATGGAAATCCGTTTGGAGACCATCG |
| IY908_PP4124_sgRNA_NT1_R | CGATGGTCTCCAAACGGATTTCCATCGACGATGAAGAGCTAAGAGACCCAGA |
| IY909_PP4129_sgRNA_NT1_F | TCTGGGTCTCTTAGCGGCTGTACGCGCCGCCCTCAGGGTTTGGAGACCATCG |
| IY910_PP4129_sgRNA_NT1_R | CGATGGTCTCCAAACCCTGAGGGCGGCGCGTACAGCCGCTAAGAGACCCAGA |
| IY911_PP4130_sgRNA_NT1_F | TCTGGGTCTCTTAGCGCGGGAGCAAGGGTGTAGTCGCGTTTGGAGACCATCG |
| IY912_PP4130_sgRNA_NT1_R | CGATGGTCTCCAAACGCGACTACACCCTTGCTCCCGCGCTAAGAGACCCAGA |
| IY913_PP4128_sgRNA_NT1_F | TCTGGGTCTCTTAGCTGACCGTCGGGCTGGACCCAACGTTTGGAGACCATCG |
| IY914_PP4128_sgRNA_NT1_R | CGATGGTCTCCAAACGTTGGGTCCAGCCCGACGGTCAGCTAAGAGACCCAGA |
| IY915_PP0999_sgRNA_NT1_F | TCTGGGTCTCTTAGCggcctcgtctggcttgtacgacGTTTGGAGACCATCG |
| IY916_PP0999_sgRNA_NT1_R | CGATGGTCTCCAAACgtcgtacaagccagacgaggccGCTAAGAGACCCAGA |
| IY917_PP1031_sgRNA_NT1_F | TCTGGGTCTCTTAGCtcggtcacggtatccatggcagGTTTGGAGACCATCG |
| IY918_PP1031_sgRNA_NT1_R | CGATGGTCTCCAAACctgccatggataccgtgaccgaGCTAAGAGACCCAGA |
| IY919_PP1770_sgRNA_NT1_F | TCTGGGTCTCTTAGCatgacatcggcgcccacacaagGTTTGGAGACCATCG |
| IY920_PP1770_sgRNA_NT1_R | CGATGGTCTCCAAACcttgtgtgggcgccgatgtcatGCTAAGAGACCCAGA |
| IY921_PP1664_sgRNA_NT1_F | TCTGGGTCTCTTAGCgttgagcaggcggccctgatagGTTTGGAGACCATCG |
| IY922_PP1664_sgRNA_NT1_R | CGATGGTCTCCAAACctatcagggccgcctgctcaacGCTAAGAGACCCAGA |
| IY923_PP1914_sgRNA_NT1_F | TCTGGGTCTCTTAGCgtaatgccgtgttctttcagggGTTTGGAGACCATCG |
| IY924_PP1914_sgRNA_NT1_R | CGATGGTCTCCAAACccctgaaagaacacggcattacGCTAAGAGACCCAGA |
| IY925_PP4175_sgRNA_NT1_F | TCTGGGTCTCTTAGCgttggaaacctgctcttcggtcGTTTGGAGACCATCG |
| IY926_PP4175_sgRNA_NT1_R | CGATGGTCTCCAAACgaccgaagagcaggtttccaacGCTAAGAGACCCAGA |
| IY927_PP4174_sgRNA_NT1_F | TCTGGGTCTCTTAGCctcaccgcgactgcagcgcagcGTTTGGAGACCATCG |
| IY928_PP4174_sgRNA_NT1_R | CGATGGTCTCCAAACgctgcgctgcagtcgcggtgagGCTAAGAGACCCAGA |
| IY929_PP4189_sgRNA_NT1_F | TCTGGGTCTCTTAGCTGTTCACTGCTCACGCTCCCGGGTTTGGAGACCATCG |
| IY930_PP4189_sgRNA_NT1_R | CGATGGTCTCCAAACCCGGGAGCGTGAGCAGTGAACAGCTAAGAGACCCAGA |
| IY931_PP4188_sgRNA_NT1_F | TCTGGGTCTCTTAGCAGGTCCTTGTACTTCGAACGCAGTTTGGAGACCATCG |
| IY932_PP4188_sgRNA_NT1_R | CGATGGTCTCCAAACTGCGTTCGAAGTACAAGGACCTGCTAAGAGACCCAGA |

| | |
|---|---|
| IY933_PP4187_sgRNA_NT1_F | TCTGGGTCTCTTAGCCAGGCAGTCTTCAGACCAAGTTGTTTGGAGACCATCG |
| IY934_PP4187_sgRNA_NT1_R | CGATGGTCTCCAAACAACTTGGTCTGAAGACTGCCTGGCTAAGAGACCCAGA |
| IY935_PP1771_sgRNA_NT1_F | TCTGGGTCTCTTAGCggatgatctgttgtagcttaccGTTTGGAGACCATCG |
| IY936_PP1771_sgRNA_NT1_R | CGATGGTCTCCAAACggtaagctacaacagatcatccGCTAAGAGACCCAGA |
| IY937_PP0691_sgRNA_NT1_F | TCTGGGTCTCTTAGCcggtcgcttggtccagcccagcGTTTGGAGACCATCG |
| IY938_PP0691_sgRNA_NT1_R | CGATGGTCTCCAAACgctgggctggaccaagcgaccgGCTAAGAGACCCAGA |
| IY939_PP0001_sgRNA_NT1_F | TCTGGGTCTCTTAGCcgccaatcgggcgtaccacgatGTTTGGAGACCATCG |
| IY940_PP0001_sgRNA_NT1_R | CGATGGTCTCCAAACatcgtggtacgcccgattggcgGCTAAGAGACCCAGA |
| IY941_PP4116_sgRNA_NT1_F | TCTGGGTCTCTTAGCgcaaaggtgtgctcaggttgcaGTTTGGAGACCATCG |
| IY942_PP4116_sgRNA_NT1_R | CGATGGTCTCCAAACtgcaacctgagcacacctttgcGCTAAGAGACCCAGA |
| IY943_PP4043_sgRNA_NT1_F | TCTGGGTCTCTTAGCctgaattgcgacagctgagggtGTTTGGAGACCATCG |
| IY944_PP4043_sgRNA_NT1_R | CGATGGTCTCCAAACaccctcagctgtcgcaattcagGCTAAGAGACCCAGA |
| IY945_PP4862_sgRNA_NT1_F | TCTGGGTCTCTTAGCgacgttcaggatcgacgcgccgGTTTGGAGACCATCG |
| IY946_PP4862_sgRNA_NT1_R | CGATGGTCTCCAAACcggcgcgtcgatcctgaacgtcGCTAAGAGACCCAGA |
| IY946_PP0103_sgRNA_NT1_F | TCTGGGTCTCTTAGCGATCCAGAAGATGGTCATGTGCGTTTGGAGACCATCG |
| IY947_PP0103_sgRNA_NT1_R | CGATGGTCTCCAAACGCACATGACCATCTTCTGGATCGCTAAGAGACCCAGA |
| IY948_PP0104_sgRNA_NT1_F | TCTGGGTCTCTTAGCGAACATGATGAAACTGAACCACGTTTGGAGACCATCG |
| IY949_PP0104_sgRNA_NT1_R | CGATGGTCTCCAAACGTGGTTCAGTTTCATCATGTTCGCTAAGAGACCCAGA |
| IY950_PP0105_sgRNA_NT1_F | TCTGGGTCTCTTAGCACTGCACCCGCACAGCACGCGTGTTTGGAGACCATCG |
| IY951_PP0105_sgRNA_NT1_R | CGATGGTCTCCAAACACGCGTGCTGTGCGGGTGCAGTGCTAAGAGACCCAGA |
| IY952_PP0106_sgRNA_NT1_F | TCTGGGTCTCTTAGCAAATCAGCGGCCCGTGGGAATCGTTTGGAGACCATCG |
| IY953_PP0106_sgRNA_NT1_R | CGATGGTCTCCAAACGATTCCCACGGGCCGCTGATTTGCTAAGAGACCCAGA |
| IY954_PP1317_sgRNA_NT1_F | TCTGGGTCTCTTAGCCCGTTCGCCGCACGATGAATACGTTTGGAGACCATCG |
| IY955_PP1317_sgRNA_NT1_R | CGATGGTCTCCAAACGTATTCATCGTGCGGCGAACGGGCTAAGAGACCCAGA |
| IY956_PP1318_sgRNA_NT1_F | TCTGGGTCTCTTAGCAGGCGAACGCCTCTTCCGCCGAGTTTGGAGACCATCG |
| IY957_PP1318_sgRNA_NT1_R | CGATGGTCTCCAAACTCGGCGGAAGAGGCGTTCGCCTGCTAAGAGACCCAGA |
| IY958_PP1319_sgRNA_NT1_F | TCTGGGTCTCTTAGCCATCAGTTCATGCGGGATACCCGTTTGGAGACCATCG |
| IY959_PP1319_sgRNA_NT1_R | CGATGGTCTCCAAACGGGTATCCCGCATGAACTGATGGCTAAGAGACCCAGA |
| IY960_PP0812_sgRNA_NT1_F | TCTGGGTCTCTTAGCGGTGGCTGCCTTGTTGGAAGCAGTTTGGAGACCATCG |
| IY961_PP0812_sgRNA_NT1_R | CGATGGTCTCCAAACTGCTTCCAACAAGGCAGCCACCGCTAAGAGACCCAGA |
| IY962_PP0813_sgRNA_NT1_F | TCTGGGTCTCTTAGCAGATCTGGTCATAGTGTTCAGGGTTTGGAGACCATCG |
| IY963_PP0813_sgRNA_NT1_R | CGATGGTCTCCAAACCCTGAACACTATGACCAGATCTGCTAAGAGACCCAGA |
| IY964_PP0814_sgRNA_NT1_F | TCTGGGTCTCTTAGCGGTGGCGAAGAGCGACGCAAACGTTTGGAGACCATCG |
| IY965_PP0814_sgRNA_NT1_R | CGATGGTCTCCAAACGTTTGCGTCGCTCTTCGCCACCGCTAAGAGACCCAGA |
| IY966_PP0815_sgRNA_NT1_F | TCTGGGTCTCTTAGCCTCTTTCGAGCGGTCCATGTGCGTTTGGAGACCATCG |
| IY967_PP0815_sgRNA_NT1_R | CGATGGTCTCCAAACGCACATGGACCGCTCGAAAGAGGCTAAGAGACCCAGA |
| IY968_PP1023_sgRNA_NT1_F | TCTGGGTCTCTTAGCcgctggccagtttctccaggaaGTTTGGAGACCATCG |
| IY969_PP1023_sgRNA_NT1_R | CGATGGTCTCCAAACttcctggagaaactggccagcgGCTAAGAGACCCAGA |
| IY970_PP1240_sgRNA_NT1_F | TCTGGGTCTCTTAGCgttacggaacagcaggatcaagGTTTGGAGACCATCG |
| IY971_PP1240_sgRNA_NT1_R | CGATGGTCTCCAAACcttgatcctgctgttccgtaacGCTAAGAGACCCAGA |
| IY972_PP5128_sgRNA_NT1_F | TCTGGGTCTCTTAGCttggagatacagacgatggcgtGTTTGGAGACCATCG |
| IY973_PP5128_sgRNA_NT1_R | CGATGGTCTCCAAACacgccatcgtctgtatctccaaGCTAAGAGACCCAGA |
| IY974_PP5046_sgRNA_NT1_F | TCTGGGTCTCTTAGCgcgaatacggtgtcaccgatgcGTTTGGAGACCATCG |

| | |
|---|---|
| IY975_PP5046_sgRNA_NT1_R | CGATGGTCTCCAAACgcatcggtgacaccgtattcgcGCTAAGAGACCCAGA |
| IY976_PP2095_sgRNA_NT1_F | TCTGGGTCTCTTAGCcttccggcaggcggaacaaccgGTTTGGAGACCATCG |
| IY977_PP2095_sgRNA_NT1_R | CGATGGTCTCCAAACcggttgttccgcctgccggaagGCTAAGAGACCCAGA |
| IY978_PP1769_sgRNA_NT1_F | TCTGGGTCTCTTAGCcacgctcgggacggtagaacacGTTTGGAGACCATCG |
| IY979_PP1769_sgRNA_NT1_R | CGATGGTCTCCAAACgtgttctaccgtcccgagcgtgGCTAAGAGACCCAGA |
| IY980_PP4635_sgRNA_NT1_F | TCTGGGTCTCTTAGCtcgcgtacgcccagcttgcgggGTTTGGAGACCATCG |
| IY981_PP4635_sgRNA_NT1_R | CGATGGTCTCCAAACcccgcaagctgggcgtacgcgaGCTAAGAGACCCAGA |
| IY982_PP1610_sgRNA_NT1_F | TCTGGGTCTCTTAGCcaaaaacttcaccgtgctggaaGTTTGGAGACCATCG |
| IY983_PP1610_sgRNA_NT1_R | CGATGGTCTCCAAACttccagcacggtgaagtttttgGCTAAGAGACCCAGA |
| IY984_PP5186_sgRNA_NT1_F | TCTGGGTCTCTTAGCatggagctttcttcgtcgcaggGTTTGGAGACCATCG |
| IY985_PP5186_sgRNA_NT1_R | CGATGGTCTCCAAACcctgcgacgaagaaagctccatGCTAAGAGACCCAGA |
| IY986_PP4161_sgRNA_NT1_F | TCTGGGTCTCTTAGCGTGTTGGTAACGAAACTGACGGGTTTGGAGACCATCG |
| IY987_PP4161_sgRNA_NT1_R | CGATGGTCTCCAAACCCGTCAGTTTCGTTACCAACACGCTAAGAGACCCAGA |
| IY988_PP4160_sgRNA_NT1_F | TCTGGGTCTCTTAGCCCTGATTCGCGTATTACCGGTGGTTTGGAGACCATCG |
| IY989_PP4160_sgRNA_NT1_R | CGATGGTCTCCAAACCACCGGTAATACGCGAATCAGGGCTAAGAGACCCAGA |
| IY990_PP4159_sgRNA_NT1_F | TCTGGGTCTCTTAGCCCAAGGCCTCCGGTGGCAAGTGGTTTGGAGACCATCG |
| IY991_PP4159_sgRNA_NT1_R | CGATGGTCTCCAAACCACTTGCCACCGGAGGCCTTGGGCTAAGAGACCCAGA |
| IY992_PP0658_sgRNA_NT1_F | TCTGGGTCTCTTAGCgccggtaggacccgaacagcggGTTTGGAGACCATCG |
| IY993_PP0658_sgRNA_NT1_R | CGATGGTCTCCAAACccgctgttcgggtcctaccggcGCTAAGAGACCCAGA |
| IY994_PP1506_sgRNA_NT1_F | TCTGGGTCTCTTAGCggtgcgtgggaagccgtcgaacGTTTGGAGACCATCG |
| IY995_PP1506_sgRNA_NT1_R | CGATGGTCTCCAAACgttcgacggcttcccacgcaccGCTAAGAGACCCAGA |
| IY996_PP1620_sgRNA_NT1_F | TCTGGGTCTCTTAGCgccccaggtgcacgcagtcggtGTTTGGAGACCATCG |
| IY997_PP1620_sgRNA_NT1_R | CGATGGTCTCCAAACaccgactgcgtgcacctggggcGCTAAGAGACCCAGA |
| IY998_PP4266_sgRNA_NT1_F | TCTGGGTCTCTTAGCctggaataacggggtaatgtcgGTTTGGAGACCATCG |
| IY999_PP4266_sgRNA_NT1_R | CGATGGTCTCCAAACcgacattaccccgttattccagGCTAAGAGACCCAGA |
| IY1000_PP1457_sgRNA_NT1_F | TCTGGGTCTCTTAGCgaagtggtacggcgaggttggcGTTTGGAGACCATCG |
| IY1001_PP1457_sgRNA_NT1_R | CGATGGTCTCCAAACgccaacctcgccgtaccacttcGCTAAGAGACCCAGA |
| IY1002_PP5203_sgRNA_NT1_F | TCTGGGTCTCTTAGCaatcggctctggaatacggaagGTTTGGAGACCATCG |
| IY1003_PP5203_sgRNA_NT1_R | CGATGGTCTCCAAACcttccgtattccagagccgattGCTAAGAGACCCAGA |
| IY1004_PP0227_sgRNA_NT1_F | TCTGGGTCTCTTAGCTTTTTCTTGCGCTCTTCGGAAAGTTTGGAGACCATCG |
| IY1005_PP0227_sgRNA_NT1_R | CGATGGTCTCCAAACTTTCCGAAGAGCGCAAGAAAAAGCTAAGAGACCCAGA |
| IY1006_PP0226_sgRNA_NT1_F | TCTGGGTCTCTTAGCAGATGAAGCTGTTGCCCAGCGGGTTTGGAGACCATCG |
| IY1007_PP0226_sgRNA_NT1_R | CGATGGTCTCCAAACCCGCTGGGCAACAGCTTCATCTGCTAAGAGACCCAGA |
| IY1008_PP0225_sgRNA_NT1_F | TCTGGGTCTCTTAGCCGCACTCTGCTGCCCGCCCAAAGTTTGGAGACCATCG |
| IY1009_PP0225_sgRNA_NT1_R | CGATGGTCTCCAAACTTTGGGCGGGCAGCAGAGTGCGGCTAAGAGACCCAGA |
| IY1010_PP4651_sgRNA_NT1_F | TCTGGGTCTCTTAGCTGACCAGAAGTGATAGAGGTCAGTTTGGAGACCATCG |
| IY1011_PP4651_sgRNA_NT1_R | CGATGGTCTCCAAACTGACCTCTATCACTTCTGGTCAGCTAAGAGACCCAGA |
| IY1012_PP4650_sgRNA_NT1_F | TCTGGGTCTCTTAGCAGGTTTCGTTGCCGTCCCATAGTTTGGAGACCATCG |
| IY1013_PP4650_sgRNA_NT1_R | CGATGGTCTCCAAACTATGGGACGGCAACGAAACCTGCTAAGAGACCCAGA |
| IY1035_PP4193_F | TCTGGGTCTCTTAGCGGATGGACGTGTACGCAGTGACGTTTGGAGACCATCG |
| IY1036_PP4193_R | CGATGGTCTCCAAACGTCACTGCGTACACGTCCATCCGCTAAGAGACCCAGA |
| IY1037_PP4192_F | TCTGGGTCTCTTAGCGGCGTCAGGTAGTCGGTTGCAAGTTTGGAGACCATCG |
| IY1038_PP4192_R | CGATGGTCTCCAAACTTGCAACCGACTACCTGACGCCGCTAAGAGACCCAGA |

| | |
|---|---|
| IY1039_PP4191_F | TCTGGGTCTCTTAGCCCGATGTAGTCGGAACCCTTGAGTTTGGAGACCATCG |
| IY1040_PP4191_R | CGATGGTCTCCAAACTCAAGGGTTCCGACTACATCGGGCTAAGAGACCCAGA |
| IY1041_PP4090_F | TCTGGGTCTCTTAGCTGCCAATACATCCAGCACCATCGTTTGGAGACCATCG |
| IY1042_PP4090_R | CGATGGTCTCCAAACGATGGTGCTGGATGTATTGGCAGCTAAGAGACCCAGA |
| IY1043_PP0897_F | TCTGGGTCTCTTAGCGGTGTCCTGGCAGATTGGACGAGTTTGGAGACCATCG |
| IY1044_PP0897_R | CGATGGTCTCCAAACTCGTCCAATCTGCCAGGACACCGCTAAGAGACCCAGA |
| IY1045_PP0944_F | TCTGGGTCTCTTAGCagcctgtctggaacacatccacGTTTGGAGACCATCG |
| IY1046_PP0944_R | CGATGGTCTCCAAACgtggatgtgttccagacaggctGCTAAGAGACCCAGA |
| IY1047_PP1755_F | TCTGGGTCTCTTAGCtgcccgtctgccagacgacgagGTTTGGAGACCATCG |
| IY1048_PP1755_R | CGATGGTCTCCAAACctcgtcgtctggcagacgggcaGCTAAGAGACCCAGA |
| IY1049_PP0654_F | TCTGGGTCTCTTAGCgccccgccgtgaacatgggtatGTTTGGAGACCATCG |
| IY1050_PP0654_R | CGATGGTCTCCAAACatacccatgttcacggcggggcGCTAAGAGACCCAGA |
| IY1051_PP0751_F | TCTGGGTCTCTTAGCcttgtcaccctcgacatagctcGTTTGGAGACCATCG |
| IY1052_PP0751_R | CGATGGTCTCCAAACgagctatgtcgagggtgacaagGCTAAGAGACCCAGA |
| IY1053_PP2925_F | TCTGGGTCTCTTAGCctcgtagcgcttcttcaggaagGTTTGGAGACCATCG |
| IY1054_PP2925_R | CGATGGTCTCCAAACcttcctgaagaagcgctacgagGCTAAGAGACCCAGA |
| IY1055_PP1251_F | TCTGGGTCTCTTAGCcacggtcgggttctcggtcaccGTTTGGAGACCATCG |
| IY1056_PP1251_R | CGATGGTCTCCAAACggtgaccgagaacccgaccgtgGCTAAGAGACCCAGA |
| IY1057_PP2112_F | TCTGGGTCTCTTAGCggagcgccgttcgccaagccatGTTTGGAGACCATCG |
| IY1058_PP2112_R | CGATGGTCTCCAAACatggcttggcgaacggcgctccGCTAAGAGACCCAGA |
| IY1059_PP2339_F | TCTGGGTCTCTTAGCtggaaggcatcgaacatcagtaGTTTGGAGACCATCG |
| IY1060_PP2339_R | CGATGGTCTCCAAACtactgatgttcgatgccttccaGCTAAGAGACCCAGA |
| IY1061_PP4011_F | TCTGGGTCTCTTAGCcgatcccgtcgccttcgatgtaGTTTGGAGACCATCG |
| IY1062_PP4011_R | CGATGGTCTCCAAACtacatcgaaggcgacgggatcgGCTAAGAGACCCAGA |
| IY1063_PP4012_F | TCTGGGTCTCTTAGCgttcacggcggagcccttgatgGTTTGGAGACCATCG |
| IY1064_PP4012_R | CGATGGTCTCCAAACcatcaagggctccgccgtgaacGCTAAGAGACCCAGA |
| IY1065_PP4185_F | TCTGGGTCTCTTAGCccaggatcgagtctttgcagaaGTTTGGAGACCATCG |
| IY1066_PP4185_R | CGATGGTCTCCAAACttctgcaaagactcgatcctggGCTAAGAGACCCAGA |
| IY1067_PP4186_F | TCTGGGTCTCTTAGCatcttggagacgggttgaccgtGTTTGGAGACCATCG |
| IY1068_PP4186_R | CGATGGTCTCCAAACacggtcaacccgtctccaagatGCTAAGAGACCCAGA |
| IY1077_BFP_sgRNA_F | TCTGGGTCTCTTAGCgggaaggactgcttgaagaagtGTTTGGAGACCATCG |
| IY1078_BFP_sgRNA_R | CGATGGTCTCCAAACacttcttcaagcagtccttcccGCTAAGAGACCCAGA |
| IY1079_PP_4960_sgRNA_F | TCTGGGTCTCTTAGCggtgcatgcacaccgggatgtgGTTTGGAGACCATCG |
| IY1080_PP_4960_sgRNA_R | CGATGGTCTCCAAACcacatcccggtgtgcatgcaccGCTAAGAGACCCAGA |
| IY1081_PP_1791_sgRNA_F | TCTGGGTCTCTTAGCaggtcaaccggcgattcactcgGTTTGGAGACCATCG |
| IY1082_PP_1791_sgRNA_R | CGATGGTCTCCAAACcgagtgaatcgccggttgacctGCTAAGAGACCCAGA |
| IY1083_PP_2871_sgRNA_F | TCTGGGTCTCTTAGCgccgtaaaactccatggacatcGTTTGGAGACCATCG |
| IY1084_PP_2871_sgRNA_R | CGATGGTCTCCAAACgatgtccatggagttttacggcGCTAAGAGACCCAGA |
| IY1085_PP_3224_sgRNA_F | TCTGGGTCTCTTAGCtgtgcagcacagccttgagatcGTTTGGAGACCATCG |
| IY1086_PP_3224_sgRNA_R | CGATGGTCTCCAAACgatctcaaggctgtgctgcacaGCTAAGAGACCCAGA |
| IY1087_PP_4715_sgRNA_F | TCTGGGTCTCTTAGCaataatctggcgacgctccgaaGTTTGGAGACCATCG |
| IY1088_PP_4715_sgRNA_R | CGATGGTCTCCAAACttcggagcgtcgccagattattGCTAAGAGACCCAGA |
| IY1089_PP0297_sgRNA_F | TCTGGGTCTCTTAGCccagacaaactcgatgggcaagGTTTGGAGACCATCG |

| | |
|---|---|
| IY1090_PP0297_sgRNA_R | CGATGGTCTCCAAACcttgcccatcgagtttgtctggGCTAAGAGACCCAGA |
| IY1091_PP0987_sgRNA_F | TCTGGGTCTCTTAGCtgccgttggggtggtagtccaaGTTTGGAGACCATCG |
| IY1092_PP0987_sgRNA_R | CGATGGTCTCCAAACttggactaccaccccaacggcaGCTAAGAGACCCAGA |
| IY1093_PP2930_sgRNA_F | TCTGGGTCTCTTAGCtttcgggtatcaccacggtcacGTTTGGAGACCATCG |
| IY1094_PP2930_sgRNA_R | CGATGGTCTCCAAACgtgaccgtggtgatacccgaaaGCTAAGAGACCCAGA |
| IY1095_PP3144_sgRNA_F | TCTGGGTCTCTTAGCcgatgcgcgattcgatcgtggaGTTTGGAGACCATCG |
| IY1096_PP3144_sgRNA_R | CGATGGTCTCCAAACtccacgatcgaatcgcgcatcgGCTAAGAGACCCAGA |
| IY1097_PP3144_sgRNA_F | TCTGGGTCTCTTAGCcgatgcgcgattcgatcgtggaGTTTGGAGACCATCG |
| IY1098_PP3144_sgRNA_R | CGATGGTCTCCAAACtccacgatcgaatcgcgcatcgGCTAAGAGACCCAGA |
| IY1099_PP3191_sgRNA_F | TCTGGGTCTCTTAGCtgcccatggttaccggatgaggGTTTGGAGACCATCG |
| IY1100_PP3191_sgRNA_R | CGATGGTCTCCAAACcctcatccggtaaccatgggcaGCTAAGAGACCCAGA |
| IY1101_PP3446_sgRNA_F | TCTGGGTCTCTTAGCcacatcggtcagtcgcgacaggGTTTGGAGACCATCG |
| IY1102_PP3446_sgRNA_R | CGATGGTCTCCAAACcctgtcgcgactgaccgatgtgGCTAAGAGACCCAGA |
| IY1103_PP5149_sgRNA_F | TCTGGGTCTCTTAGCccttcgaccttgatctccggagGTTTGGAGACCATCG |
| IY1104_PP5149_sgRNA_R | CGATGGTCTCCAAACctccggagatcaaggtcgaaggGCTAAGAGACCCAGA |
| IY1105_PP0731_sgRNA_F | TCTGGGTCTCTTAGCgcacccgggaaataacacttgaGTTTGGAGACCATCG |
| IY1106_PP0731_sgRNA_R | CGATGGTCTCCAAACtcaagtgttatttcccgggtgcGCTAAGAGACCCAGA |
| IY1107_PP3664_sgRNA_F | TCTGGGTCTCTTAGCgtggtagcgatccaggcgatagGTTTGGAGACCATCG |
| IY1108_PP3664_sgRNA_R | CGATGGTCTCCAAACctatcgcctggatcgctaccacGCTAAGAGACCCAGA |
| IY1109_PP4677_sgRNA_F | TCTGGGTCTCTTAGCagcgagtcatactcggcaccgaGTTTGGAGACCATCG |
| IY1110_PP4677_sgRNA_R | CGATGGTCTCCAAACtcggtgccgagtatgactcgctGCTAAGAGACCCAGA |
| IY1111_PP0082_sgRNA_F | TCTGGGTCTCTTAGCcggcgtggtctggttatccacaGTTTGGAGACCATCG |
| IY1112_PP0082_sgRNA_R | CGATGGTCTCCAAACtgtggataaccagaccacgccgGCTAAGAGACCCAGA |
| IY1113_PP0083_sgRNA_F | TCTGGGTCTCTTAGCtggcacccatgtagatcacgcaGTTTGGAGACCATCG |
| IY1114_PP0083_sgRNA_R | CGATGGTCTCCAAACtgcgtgatctacatgggtgccaGCTAAGAGACCCAGA |
| IY1115_PP0988_sgRNA_F | TCTGGGTCTCTTAGCgcagcagtggcttcgtcgagcaGTTTGGAGACCATCG |
| IY1116_PP0988_sgRNA_R | CGATGGTCTCCAAACtgctcgacgaagccactgctgcGCTAAGAGACCCAGA |
| IY1117_PP5192_sgRNA_F | TCTGGGTCTCTTAGCtgttcggcgtaaccggcgagctGTTTGGAGACCATCG |
| IY1118_PP5192_sgRNA_R | CGATGGTCTCCAAACagctcgccggttacgccgaacaGCTAAGAGACCCAGA |
| IY1139_PP0321_sgRNA_F | TCTGGGTCTCTTAGCcggaagtactccgatgcacgctGTTTGGAGACCATCG |
| IY1140_PP0321_sgRNA_R | CGATGGTCTCCAAACagcgtgcatcggagtacttccgGCTAAGAGACCCAGA |
| IY1141_PP1808_sgRNA_F | TCTGGGTCTCTTAGCgccgcaagcgctaagggaaaacGTTTGGAGACCATCG |
| IY1142_PP1808_sgRNA_R | CGATGGTCTCCAAACgttttcccttagcgcttgcggcGCTAAGAGACCCAGA |
| IY1143_PP4701_sgRNA_F | TCTGGGTCTCTTAGCgcgtcctgcaggcccacttcatGTTTGGAGACCATCG |
| IY1144_PP4701_sgRNA_R | CGATGGTCTCCAAACatgaagtgggcctgcaggacgcGCTAAGAGACCCAGA |
| IY1145_PP1011_sgRNA_F | TCTGGGTCTCTTAGCgatcaccagtgccggcctcgagGTTTGGAGACCATCG |
| IY1146_PP1011_sgRNA_R | CGATGGTCTCCAAACctcgaggccggcactggtgatcGCTAAGAGACCCAGA |
| IY1147_PP1156_sgRNA_F | TCTGGGTCTCTTAGCcgtggcgtaccactttcaagtgGTTTGGAGACCATCG |
| IY1148_PP1156_sgRNA_R | CGATGGTCTCCAAACcacttgaaagtggtacgccacgGCTAAGAGACCCAGA |
| IY1149_PP2149_sgRNA_F | TCTGGGTCTCTTAGCtggacggacagttcggtttcgtGTTTGGAGACCATCG |
| IY1150_PP2149_sgRNA_R | CGATGGTCTCCAAACacgaaaccgaactgtccgtccaGCTAAGAGACCCAGA |
| IY1151_PP4963_sgRNA_F | TCTGGGTCTCTTAGCcggcagagaattcgccttcggtGTTTGGAGACCATCG |
| IY1152_PP4963_sgRNA_R | CGATGGTCTCCAAACaccgaaggcgaattctctgccgGCTAAGAGACCCAGA |

| | |
|---|---|
| IY1153_PP5056_sgRNA_F | TCTGGGTCTCTTAGCccgaggttcatatgaccgacttGTTTGGAGACCATCG |
| IY1154_PP5056_sgRNA_R | CGATGGTCTCCAAACaagtcggtcatatgaacctcggGCTAAGAGACCCAGA |
| IY1155_PP1612_sgRNA_F | TCTGGGTCTCTTAGCtcacggatcgggccgttgatgtGTTTGGAGACCATCG |
| IY1156_PP1612_sgRNA_R | CGATGGTCTCCAAACacatcaacggcccgatccgtgaGCTAAGAGACCCAGA |
| IY1157_PP1362_sgRNA_F | TCTGGGTCTCTTAGCgatgcggatcttcggaccctgcGTTTGGAGACCATCG |
| IY1158_PP1362_sgRNA_R | CGATGGTCTCCAAACgcagggtccgaagatccgcatcGCTAAGAGACCCAGA |
| IY1211_PP5075_F | TCTGGGTCTCTTAGCACACGGCCGCATACTTCCGGCAGTTTGGAGACCATCG |
| IY1212_PP5075_R | CGATGGTCTCCAAACTGCCGGAAGTATGCGGCCGTGTGCTAAGAGACCCAGA |
| IY1213_PP5076_F | TCTGGGTCTCTTAGCCGAAGCGTGCTCCGGGGACTCCGTTTGGAGACCATCG |
| IY1214_PP5076_R | CGATGGTCTCCAAACGGAGTCCCCGGAGCACGCTTCGGCTAAGAGACCCAGA |
| IY1215_PP0675_F | TCTGGGTCTCTTAGCacgcggaaaagtacggcgcgctGTTTGGAGACCATCG |
| IY1216_PP0675_R | CGATGGTCTCCAAACagcgcgccgtacttttccgcgtGCTAAGAGACCCAGA |
| IY1217_PP2080_F | TCTGGGTCTCTTAGCggcagtagcagaagcggccataGTTTGGAGACCATCG |
| IY1218_PP2080_R | CGATGGTCTCCAAACtatggccgcttctgctactgccGCTAAGAGACCCAGA |
| IY1219_PP5046_NT2_F | TCTGGGTCTCTTAGCgccgagtaggccagcattaccgGTTTGGAGACCATCG |
| IY1220_PP5046_NT2_R | CGATGGTCTCCAAACcggtaatgctggcctactcggcGCTAAGAGACCCAGA |
| IY1221_PP3591_F | TCTGGGTCTCTTAGCgcaaggcgcagcgaaagcgatgGTTTGGAGACCATCG |
| IY1222_PP3591_R | CGATGGTCTCCAAACcatcgctttcgctgcgccttgcGCTAAGAGACCCAGA |
| IY1223_PP2336_F | TCTGGGTCTCTTAGCcggcagacgcatccaggaggcgGTTTGGAGACCATCG |
| IY1224_PP2336_R | CGATGGTCTCCAAACcgcctcctggatgcgtctgccgGCTAAGAGACCCAGA |
| IY1225_PP2652_F | TCTGGGTCTCTTAGCatggcaacgccccgcttccagtGTTTGGAGACCATCG |
| IY1226_PP2652_R | CGATGGTCTCCAAACactggaagcggggcgttgccatGCTAAGAGACCCAGA |
| IY1227_PP3662_F | TCTGGGTCTCTTAGCgatagtcttggcattgtcgacaGTTTGGAGACCATCG |
| IY1228_PP3662_R | CGATGGTCTCCAAACtgtcgacaatgccaagactatcGCTAAGAGACCCAGA |
| IY1229_PP4012_F | TCTGGGTCTCTTAGCccagcttgacccaatcctggatGTTTGGAGACCATCG |
| IY1230_PP4012_R | CGATGGTCTCCAAACatccaggattgggtcaagctggGCTAAGAGACCCAGA |
| IY1231_PP0356_F | TCTGGGTCTCTTAGCgaccagttgtgggccggccatgGTTTGGAGACCATCG |
| IY1232_PP0356_R | CGATGGTCTCCAAACcatggccggcccacaactggtcGCTAAGAGACCCAGA |
| IY77_dCas9_F | cgaatcttggagctcccgctg |
| IY169_termB15_R | cagtctttcgactgagcctttcg |
| IY425_RSF1010_R | ggttcgaccatgcgcaagc |

**Supplementary Note 1**

Pearson correlation analysis between gene expression and isoprenol production from all strains can be found here: https://benchling.com/s/etr-mta4DgBAatF0hFYzy275?m=slm-s52wQQYHQWxFJGHGC79z. Data used as an input for Pearson correlation analysis can be found here: https://benchling.com/s/etr-l20YX8nWcCM66vIvFUZf?m=slm-b4jTtjqlVo4ElPg5bjn3. Python script to generate the Pearson correlation analysis is detailed below.

```
import pandas as pd
from scipy.stats import pearsonr

# Load your data (adjust the filename or data import method as needed)
data = pd.read_csv('/path/to/your/data.csv')

# Initialize lists to store correlation coefficients and p-values
correlations = []
p_values = []

# Iterate over each protein column (excluding strain and isoprenol_production)
for protein in data.columns[2:]:  # Assuming 1st column is 'Strain', 2nd is 'isoprenol_production'
    protein_expression = data[protein]
    isoprenol_production = data['isoprenol_production']

    # Check if there is enough variability in the protein expression data
    if protein_expression.nunique() > 1:
        corr, p_val = pearsonr(protein_expression, isoprenol_production)
    else:
        # If no variability in protein expression, correlation and p-value are not applicable
        corr, p_val = float('nan'), float('nan')

    # Append results to the lists
    correlations.append(corr)
    p_values.append(p_val)

# Create a DataFrame with the results
results = pd.DataFrame({
    'Protein': data.columns[2:],  # Protein names
    'Correlation_with_isoprenol_production': correlations,
    'p_value': p_values
})

# Filter out non-significant correlations (p-value >= 0.05)
significant_results = results[results['p_value'] < 0.05]

# Save the filtered results to a new CSV file (optional)
significant_results.to_csv('significant_protein_isoprenol_correlation.csv', index=False)

# Display the top significant correlations by absolute value (strongest correlations)
top_significant_results = significant_results.dropna().sort_values(by='Correlation_with_isoprenol_production',
key=abs, ascending=False)
print(top_significant_results.head(10))
```

### Supplementary Note 2

The construction of multiplexed CRISPRi plasmids involves two main steps: (1) the creation of sgRNA storage plasmids and (2) the assembly of CRISPRi plasmids. The detailed protocol for constructing CRISPRi plasmids containing multiple sgRNAs is provided below.

**Step I: Creation of sgRNA storage plasmids**

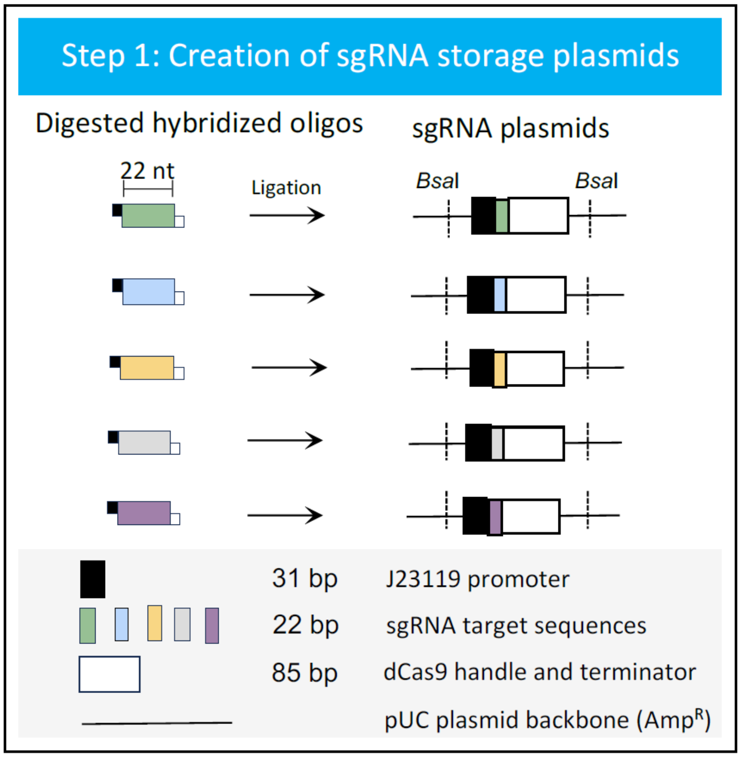


**A. Hybridization of oligos**

An sgRNA consists of a J23119 promoter (31 bp), an sgRNA target sequence (22 bp), and a dCas9 handle with a terminator (85 bp). The sgRNA target sequence is synthesized as a pair of oligos (see Supplementary Table S6).

To prepare the oligos for hybridization, dissolve the lyophilized oligos in 10 mM TRIS-HCl buffer (pH 7.9) or $ddH_2O$ to a final concentration of 100 µM as stock solutions. Mix the forward and reverse oligos (refer to the table below), then hybridize them by heating at 95°C for 10 minutes in a heating block. Allow the mixture to cool to room temperature on the heating block.
The hybridized oligos can be stored at -20°C for several months.

Annealing buffer: 10mM TRIS-HCl buffer pH7.9, 100mM NaCl, 10mM $MgCl_2$

| | Volume to add | stock concentration | final concentration |
|---|---|---|---|
| Annealing buffer | 49 µl | | |
| Forward oligo | 0.5 µl | 100 µM | 1 µM |

| | | | |
|---|---|---|---|
| Reverse oligo | 0.5 µl | 100 µM | 1 µM |
| | TOTAL Vol 50 µl | | |

**B. Digestion of hybridized oligos and cloning to plasmid pIY993 (pJET-J23119-sfGFP-dCas9 handle and terminator)**

1. Digestion of hybridized oligos by *Bsa*I-HFv2

Hybridized oligos = 5 µL
rCutSmart™ Buffer (NEB #B6004) = 3 µL
*Bsa*I-HFv2 (NEB #R3733S) = 0.5 µL
Nuclease-free water (NEB #B1500S) = 21.5 µL

Incubate at 37 $^{o}$C for 1 hr, heat inactivate at 65 $^{o}$C for 20 min.

2. Digestion of plasmid pIY993 by *PaqC*I

Plasmid pIY993 = 1.5 µL (100 ng/µL)
*PaqC*I buffer = 2 µL
*PaqC*I (NEB #R0745S) = 1 µL
*PaqC*I activator (from the kit) = 0.5 µL
Nuclease-free water (NEB #B1500S) = 15 µL

Incubate at 37 $^{o}$C for 1 hr, heat inactivate at 65 $^{o}$C for 20 min.

**C. Purification by magnetic beads**

1. Shake magnetic bead stock solution (Axygen® AxyPrep Magnetic Bead) before use
2. Add 50 µL magnetic beads for each PCR tube reaction.
3. Mix by pipetting 10 times
4. Incubate at room temperature for 3 min
5. Place the PCR tubes on the magnetic plate (DynaMag-96, Thermo Fisher Scientific)
6. Let the magnetic beads settle for 3 min
7. Completely remove the supernatant while the PCR tubes are still on the magnetic plate
8. Add 130 µL 70% ethanol to each PCR tube
9. Remove the ethanol
10. Add another 130 µL 70% ethanol
11. Completely remove the ethanol
12. Remove the PCR tube from the magnetic plate
13. Elute the DNA from the beads by adding 50 µL nuclease-free water to the PCR tubes
14. Mix by pipetting 10 times
15. Incubate at room temperature for 3 min
16. Place the PCR tubes on the magnetic plate
17. Let the beads settle for 3 min
18. Transfer the supernatant (your DNA) into a fresh PCR tube

**D. Ligation of digested hybridized (double-stranded) oligos and digested plasmid pIY993**

For each reaction:

| | | |
|---|---|---|
| *Bsa*I-digested oligos | = | 3 µL |
| *Paq*CI-digested plasmid pIY993 | = | 1 µL |
| T4 DNA ligase (NEB #M0202S) | = | 0.5 µL |
| T4 DNA ligase buffer (NEB #B0202S) | = | 0.5 µL |

Incubate at 25 $^{o}$C for 15 min.

**E. Transformation**

1. Add 5 µL of ligation reaction to 15 µL of XL1-Blue competent cells.
2. Incubate on ice for 10 min
3. Heat shock at 42 $^{o}$C for 45 sec
4. Put it back on ice for 2 min
5. Add 1 mL of LB and incubate at 37 $^{o}$C for 1 h with shaking
6. Centrifuge at max speed for 1 min
7. Remove 900 µL of the supernatant
8. Resuspend the cell pellet
9. Plate all the cells on LB agar plate containing carbenicillin (100 mg/L)
10. Incubate at 37 $^{o}$C, overnight.

**F. Colony picking and DNA miniprep**

1. Place the agar plate on top of the blue-light illuminator plate
2. Pick the white colony (non GFP colony)
3. Inoculate into 5 mL LB media containing carbenicillin (100 mg/L)
4. Incubate overnight at 37 $^{o}$C
5. Miniprep on the next day and verify the sequence by Sanger sequencing

**Step 2: Creation of CRISPRi plasmids**

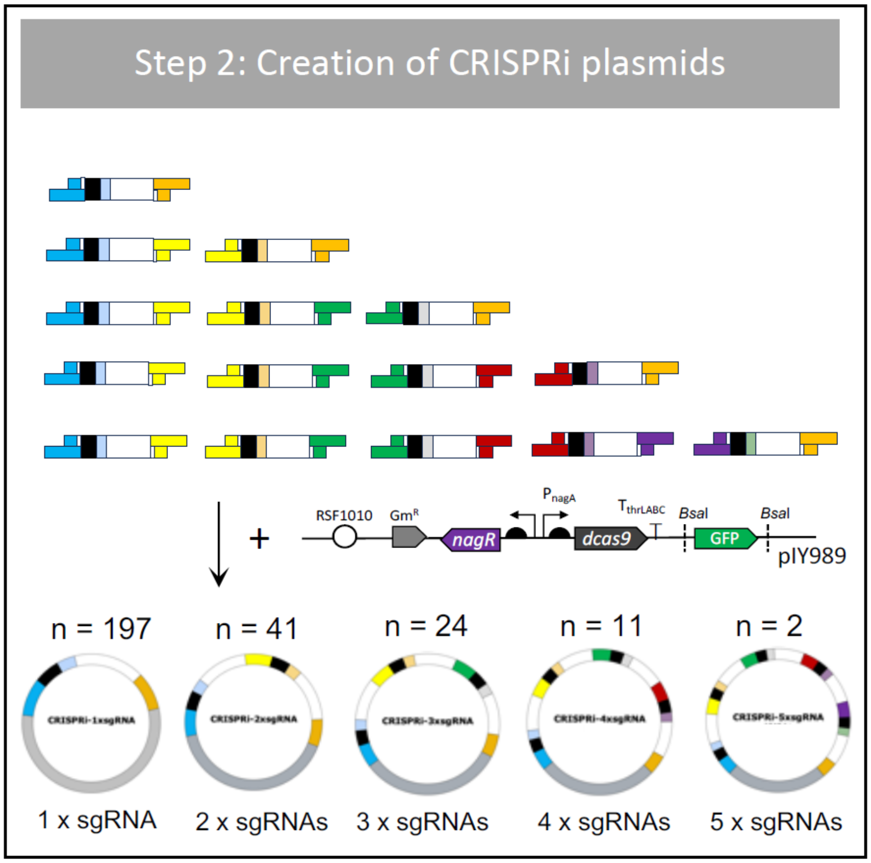


A CRISPRi plasmid consists of a dCas9 protein regulated by the PnagAa promoter and one or more sgRNAs. The sgRNA is cloned into the destination vector pIY989 (pRSF1010-Gm-nagR-dCas9-sfGFP) using a linker-based cloning method (Storch et al., 2015). Both the destination vector and the sgRNA plasmids must be digested and ligated with prefix and suffix linkers before assembly. A detailed protocol for constructing CRISPRi plasmids is provided below.

Prefix and suffix linkers are synthesized as pairs of oligos, with six pairs of linkers available (see the table below). Each prefix and suffix linker pair features complementary overhang sequences. For instance, the overhang sequence of a suffix linker (e.g., 1S) is complementary to that of the corresponding prefix linker (e.g., 1P). To create a functional linker, each oligo must first undergo phosphorylation followed by hybridization. These six pairs of prefix and suffix linkers enable the assembly of up to five sgRNAs in a single one-pot assembly reaction.

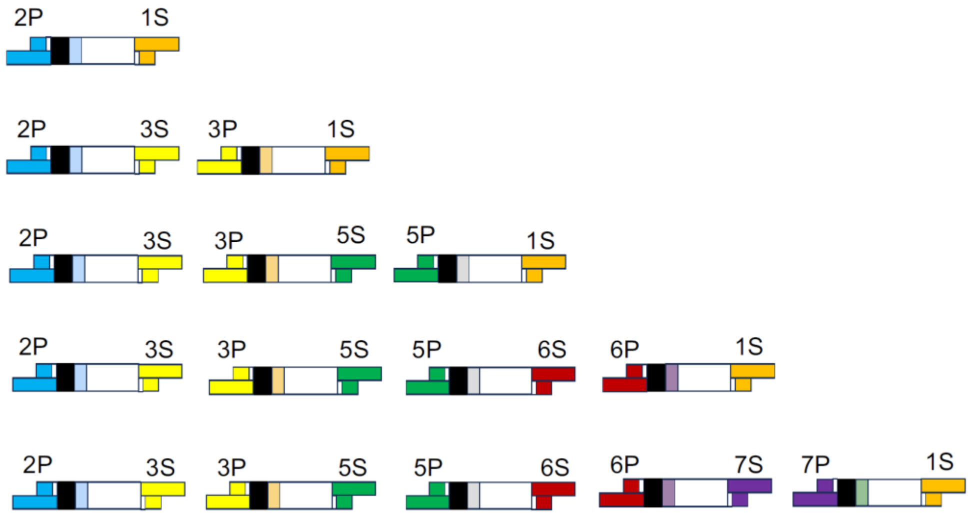


| Oligo Name | Sequence (5' to 3') | Linker Name |
|---|---|---|
| 1S_L | CTCGTTACTTACGACACTCCGAGACAGTCAGAGGGTA | 1S |
| 1S_A | TGTCGTAAGTAA | |
| 1P_L | GGACTAGTTCAATAAATACCCTCTGACTGTCTCGGAG | 1P |
| 1P_A | TTTATTGAACTA | |
| 2S_L | CTCGATCGGTGTGAAAAGTCAGTATCCAGTCGTGTAG | 2S |
| 2S_A | TTTCACACCGAT | |
| 2P_L | GGACAGGTAATAAGAACTACACGACTGGATACTGACT | 2P |
| 2P_A | TTCTTATTACCT | |
| 3S_L | CTCGATCACGGCACTACACTCGTTGCTTTATCGGTAT | 3S |
| 3S_A | TAGTGCCGTGAT | |
| 3P_L | GGACTCTGTAATAACAATACCGATAAAGCAACGAGTG | 3P |
| 3P_A | TGTTATTACAGA | |
| 5S_L | CTCGAGAAGTAGTGCCACAGACAGTATTGCTTACGAG | 5S |
| 5S_A | GGCACTACTTCT | |
| 5P_L | GGACAGGATAAATCAACTCGTAAGCAATACTGTCTGT | 5P |
| 5P_A | TTGATTTATCCT | |
| 6S_L | CTCGGTATTGTAAAGCACGAAACCTACGATAAGAGTG | 6S |
| 6S_A | GCTTTACAATAC | |
| 6P_L | GGACAAGGAGAACTGACACTCTTATCGTAGGTTTCGT | 6P |
| 6P_A | TCAGTTCTCCTT | |
| 7S_L | CTCGAACTTTTACGGGTGCCGACTCACTATTACAGAC | 7S |
| 7S_A | CCCGTAAAAGTT | |
| 7P_L | GGACAGATTGTAGTAAGTCTGTAATAGTGAGTCGGCA | 7P |
| 7P_A | TTACTACAATCT | |

**Part 1: Conditioning of sgRNAs**

**A. Prefix Linker Preparation**

1. Phosphorylation of prefix adapter oligo

| | | |
|---|---|---|
| T4 DNA ligase buffer (NEB #B0202S) | = | 5 μL |
| Adapter oligo (100 uM) (e.g., 2P_A) | = | 5 μL |
| T4 PNK (NEB #M0201S) | = | 1 μL |
| Nuclease-free water (NEB #B1500S) | = | 39 μL |

Incubate at 37 °C for 30 min in the thermocycler.

2. Phosphorylation of prefix linker oligo

| | | |
|---|---|---|
| T4 DNA ligase buffer (NEB #B0202S) | = | 5 μL |
| Linker oligo (100 uM) (e.g., 2P_L) | = | 5 μL |
| T4 PNK (NEB #M0201S) | = | 1 μL |
| $H_2O$ | = | 39 μL |

Incubate at 37 °C for 30 min in the thermocycler.

3. Hybridization of prefix linker

| | | |
|---|---|---|
| Phosphorylated prefix adapter oligo (e.g., 2P_A) | = | 50 μL |
| Phosphorylated prefix linker oligo (e.g., 2P_L) | = | 50 μL |
| Annealing buffer (10mM TRIS-HCl buffer pH7.9, 100mM NaCl, 10mM $MgCl_2$) | = | 400 μL |

Incubate at 95°C for 10 minutes on a heating block, then allow the mixture to cool to room temperature on the same heating block.

Prefix linker No. 2 (2P)

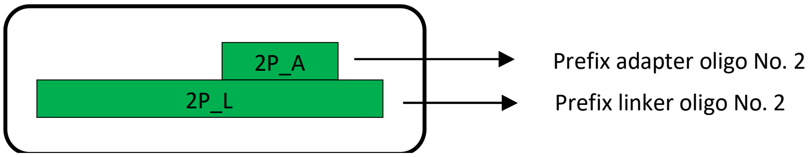


**B. Suffix Linker Preparation**

In general, the protocol is the same as above, except here suffix oligos are used.

1. Phosphorylatin of suffix adapter oligo

| | | |
|---|---|---|
| T4 ligase buffer (NEB #B0202S) | = | 5 μL |
| Adapter oligo (100 uM) (e.g., 1S_A) | = | 5 μL |
| T4 PNK (NEB #M0201S) | = | 1 μL |
| Nuclease-free water (NEB #B1500S) | = | 39 μL |

Incubate at 37 °C for 30 min in the thermocycler.

2. Phosphorylation of suffix linker oligo

| | | |
|---|---|---|
| T4 ligase buffer (NEB #B0202S) | = | 5 µL |
| Linker oligo (100 uM) (e.g., 1S_L) | = | 5 µL |
| T4 PNK (NEB #M0201S) | = | 1 µL |
| Nuclease-free water (NEB #B1500S) | = | 39 µL |

Incubate at 37 °C for 30 min in the thermocycler.

3. Hybridization of suffix linker

| | | |
|---|---|---|
| Phosphorylated suffix adapter oligo (e.g., 1S_A) | = | 50 µL |
| Phosphorylated suffix linker oligo (e.g., 1S_L) | = | 50 µL |
| Annealing buffer (10mM TRIS-HCl buffer pH7.9, 100mM NaCl, 10mM $MgCl_2$) | = | 400 µL |

Incubate at 95 °C for 10 min on a heating block and let them cool down to room temperature on a heating block.

Suffix linker No. 1 (1S)

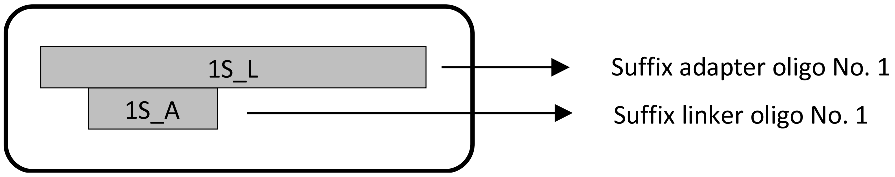


**C. Simultaneous digestion of the sgRNA plasmid (pJET-J23119-sgRNA) and ligation with linkers**

| | | |
|---|---|---|
| Plasmid (100-200 ng/µL) | = | 4 µL |
| Hybridized prefix linker (e.g., 2P) | = | 5 µL |
| Hybridized suffix linker (e.g., 1S) | = | 5 µL |
| T4 ligase buffer (NEB #B0202S) | = | 3 µL |
| *Bsa*I-HFv2 (NEB #R3733S) | = | 0.5 µL |
| T4 DNA ligase (NEB #M0202S) | = | 0.5 µL |
| Nuclease-free water (NEB #B1500S) | = | 12 µL |

Digestion and ligation condition in the thermocycler:

Stage 1 (**20 cycles**)
37 °C for 2 min
20 °C for 1 min

Stage 2 (**1 cycle**)
60 °C for 20 min
Stage 3

Storage condition 4 °C

At this stage, the sgRNA has been ligated to the linkers. However, excess linkers that have not been ligated to the DNA remain in the mixture. These unligated linkers will be removed in the next purification step.

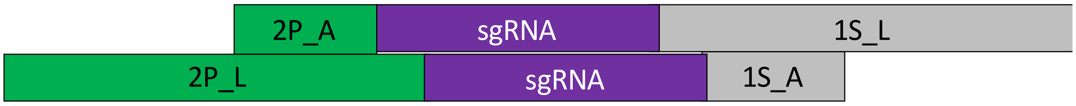


**D. Purification by magnetic beads**

1. Shake magnetic bead stock solution (Axygen® AxyPrep Magnetic Bead) before use
2. Add 50 µL magnetic beads for each PCR tube reaction.
3. Mix by pipetting 10 times
4. Incubate at room temperature for 3 min
5. Place the PCR tubes on the magnetic plate (DynaMag-96, Thermo Fisher Scientific)
6. Let the magnetic beads settle for 3 min
7. Completely remove the supernatant while the PCR tubes are still on the magnetic plate
8. Add 130 µL 70% ethanol to each PCR tube
9. Remove the ethanol
10. Add another 130 µL 70% ethanol
11. Completely remove the ethanol
12. Remove the PCR tube from the magnetic plate
13. Elute the DNA from the beads by adding 50 µL nuclease-free water to the PCR tubes
14. Mix by pipetting 10 times
15. Incubate at room temperature for 3 min
16. Place the PCR tubes on the magnetic plate
17. Let the beads settle for 3 min
18. Transfer the supernatant (your DNA) into a fresh PCR tube

At this point, the sgRNA has been successfully ligated to its corresponding prefix and suffix linkers. The next step is to prepare the destination vector (pIY989).

**Part II: Conditioning of the destination vector (pIY989)**

The protocol is similar to the one described above; however, in this step, different linkers and a different plasmid will be used.

**A. Prefix Linker Preparation**

1. Phosphorylation of prefix adapter oligo

| | | |
|---|---|---|
| T4 DNA ligase buffer (NEB #B0202S) | = | 5 µL |
| Adapter oligo (100 uM) (e.g., 1P_A) | = | 5 µL |
| T4 PNK (NEB #M0201S) | = | 1 µL |
| Nuclease-free water (NEB #B1500S) | = | 39 µL |

Incubate at 37 °C for 30 min in the thermocycler.

2. Phosphorylation of prefix linker oligo

| | | |
|---|---|---|
| T4 DNA ligase buffer (NEB #B0202S) | = | 5 μL |
| Linker oligo (100 uM) (e.g., 1P_L) | = | 5 μL |
| T4 PNK (NEB #M0201S) | = | 1 μL |
| $H_2O$ | = | 39 μL |

Incubate at 37 °C for 30 min in the thermocycler.

3. Annealing of prefix linker

| | | |
|---|---|---|
| Phosphorylated prefix adapter oligo (e.g., 1P_A) | = | 50 μL |
| Phosphorylated prefix linker oligo (e.g., 1P_L) | = | 50 μL |
| Annealing buffer (10mM TRIS-HCl buffer pH7.9, 100mM NaCl, 10mM $MgCl_2$) | = | 400 μL |

Incubate at 95°C for 10 minutes on a heating block, then allow the mixture to cool to room temperature on the same heating block.

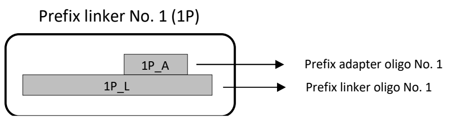


**B. Suffix Linker Preparation**

1. Phosphorylatin of suffix adapter oligo

| | | |
|---|---|---|
| T4 ligase buffer (NEB #B0202S) | = | 5 μL |
| Adapter oligo (100 uM) (e.g., 2S_A) | = | 5 μL |
| T4 PNK (NEB #M0201S) | = | 1 μL |
| Nuclease-free water (NEB #B1500S) | = | 39 μL |

Incubate at 37 °C for 30 min in the thermocycler.

2. Phosphorylation of suffix linker oligo

| | | |
|---|---|---|
| T4 ligase buffer (NEB #B0202S) | = | 5 μL |
| Linker oligo (100 uM) (e.g., 2S_L) | = | 5 μL |
| T4 PNK (NEB #M0201S) | = | 1 μL |
| Nuclease-free water (NEB #B1500S) | = | 39 μL |

Incubate at 37 °C for 30 min in the thermocycler.

3. Hybridization of suffix linker

| | | |
|---|---|---|
| Phosphorylated suffix adapter oligo (e.g., 2S_A) | = | 50 µL |
| Phosphorylated suffix linker oligo (e.g., 2S_L) | = | 50 µL |
| Annealing buffer (10mM TRIS-HCl buffer pH7.9, 100mM NaCl, 10mM $MgCl_2$) | = | 400 µL |

Incubate at 95 °C for 10 min on a heating block and let them cool down to room temperature on a heating block.

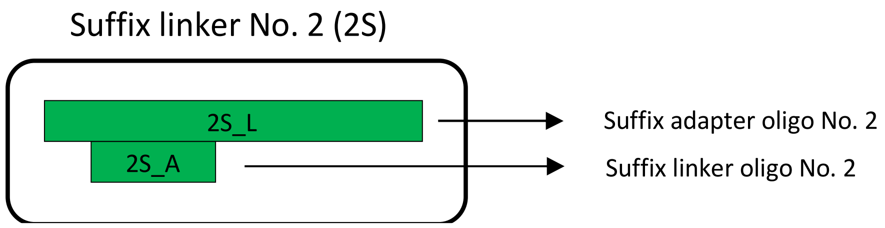


**C. Simultaneous digestion of the CRISPRi plasmid (pIY989: pRSF1010-Gm-nagR-dCas9-sfGFP) and ligation with linkers**

| | | |
|---|---|---|
| Plasmid pIY989 (100-200 ng/µL) | = | 4 µL |
| Hybridized prefix linker (e.g., 1P) | = | 5 µL |
| Hybridized suffix linker (e.g., 2S) | = | 5 µL |
| T4 ligase buffer (NEB #B0202S) | = | 3 µL |
| *Bsa*I-HFv2 (NEB #R3733S) | = | 0.5 µL |
| T4 DNA ligase (NEB #M0202S) | = | 0.5 µL |
| Nuclease-free water (NEB #B1500S) | = | 12 µL |

Digestion and ligation condition in the thermocycler:

Stage 1 (**20 cycles**)
37 °C for 2 min
20 °C for 1 min

Stage 2 (**1 cycle**)
60 °C for 20 min
Stage 3
Storage condition 4 °C

At this stage, the CRISPRi destination vector or plasmid backbone has been ligated to the linkers. However, excess linkers that have not been ligated to the DNA remain in the mixture. These unligated linkers will be removed in the next purification step.

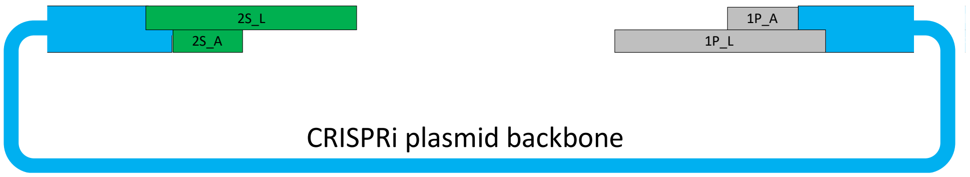


**D. Purification by magnetic beads**

1. Shake magnetic bead stock solution (Axygen® AxyPrep Magnetic Bead) before use
2. Add 50 µL magnetic beads for each PCR tube reaction.
3. Mix by pipetting 10 times
4. Incubate at room temperature for 3 min
5. Place the PCR tubes on the magnetic plate
6. Let the beads settle for 3 min
7. Completely remove the supernatant while the PCR tubes are still on the magnetic plate
8. Add 130 µL 70% ethanol to each PCR tube
9. Remove the ethanol
10. Add another 130 µL 70% ethanol
11. Completely remove the ethanol
12. Remove the PCR tube from the magnetic plate
13. Elute the DNA from the beads by adding 50 µL nuclease-free water to the PCR tubes
14. Mix by pipetting 10 times
15. Incubate at room temperature for 3 min
16. Place the PCR tubes on the magnetic plate
17. Let the beads settle for 3 min
18. Transfer the supernatant (your DNA) into a fresh PCR tube

**Part III: Plasmid Assembly**

In this step, the sgRNA (from Part I) and the digested CRISPRi destination vector (from Part II) undergo isothermal assembly at 50°C for 1 hour.

**A. Plasmid assembly**

| | | |
|---|---|---|
| sgRNA | = | 1 µL |
| Destination vector | = | 1 µL |
| rCutSmart™ Buffer (NEB #B6004) | = | 0.5 µL |
| Nuclease-free water (NEB #B1500S) | = | 2.5 µL |

Incubate at 50 °C for 1 hr in the thermocycler.

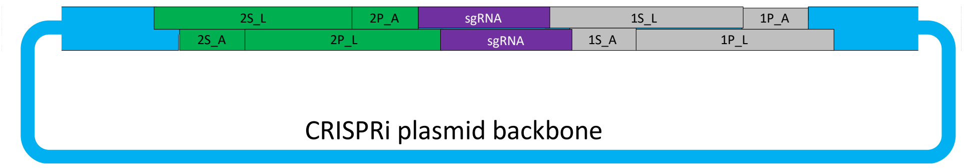


**B. Transformation**

1. Add 5 µL of plasmid assembly product to 15 µL of XL1-Blue competent cells.
2. Incubate on ice for 10 minutes.
3. Perform heat shock at 42°C for 45 seconds.
4. Return the cells to ice for 2 minutes.
5. Add 1 mL of LB medium and incubate at 37°C for 1 hour with shaking.
6. Centrifuge at maximum speed for 1 minute.
7. Discard 900 µL of the supernatant.
8. Resuspend the cell pellet.
9. Plate the entire suspension onto an LB agar plate containing gentamicin (10 mg/L).
10. Incubate overnight at 37°C.

**C. Colony picking**

1. Place the agar plate on top of the blue-light illuminator plate
2. Pick the white colony (non GFP colony)
3. Inoculate into 5 mL LB media containing gentamicin (10 mg/L)
4. Incubate overnight at 37 °C
5. Miniprep on the next day
6. Sequence with primer IY77
